# RNAseq analysis reveals dynamic metaboloepigenetic profiles of human, mouse and bovine pre-implantation embryos

**DOI:** 10.1101/2021.06.25.449773

**Authors:** Marcella Pecora Milazzotto, Michael James Noonan, Marcia de Almeida Monteiro Melo Ferraz

## Abstract

Metaboloepigenetic regulation (metabolites regulating the cellular epigenome inducing long-term changes), has been reported in stem cells, germ cells and tumor cells. Embryonic metaboloepigenetics, however, have just begun to be described. Here we analyzed RNAseq data to characterize the metaboloepigenetic profiles of human, mouse and bovine pre-implantation embryos. In embryos, metaboloepigenetic reprogramming is species specific, varies with the developmental stage and is disrupted with *in vitro* culture. Although the idea that the *in vitro* environment may influence embryo development is not new, there has been little progress on improving pregnancy rates after more than 43 years using *in vitro* fertilization. Hence, the present data on embryonic metaboloepigenetic will contribute to understanding how the *in vitro* manipulation affects the metaboloepigenetic status of early embryos, which can be used to establish culture strategies aimed at improving the *in vitro* environment and, consequently, pregnancy rates and offspring health.

## Introduction

In mammals, epigenetic reprogramming is central to embryonic survival, cell differentiation, and ensuring the proper development of a new organism. Among the different layers of epigenetic control, the methylation of DNA and histone residues are vital to normal cell function and survival. In particular, DNA methylation is involved in the control of transposable elements, X chromosome inactivation, genomic imprinting and cell differentiation (Jansz, 2019; Li and Sasaki, 2011; Sharp et al., 2011). DNA demethylation is also essential for the loss of highly repressive markers on gametes and the establishment of new specific markers, allowing embryonic cells to become totipotent and promoting adequate cell differentiation. Histone methylation also undergoes intense modifications during gametogenesis and early embryogenesis and is crucial for the establishment of totipotency (Ross et al., 2008; Sarmento et al., 2004; Torres-Padilla et al., 2007; Zhang et al., 2012).

During the period over which they undergo epigenetic reprogramming, early embryos are highly sensitive to changes in environmental conditions both *in vivo* (maternal health, diet, medication, etc.) and *in vitro* (imposed by *in vitro* oocyte maturation, *in vitro* fertilization (IVF) and *in vitro* embryo production) (El Hajj and Haaf, 2013). Under *in vitro* conditions, gametes and embryos are washed and exposed to incubations in different culture media, temperatures and oxygen concentrations (Ménézo et al., 2015; Ventura-Juncá et al., 2015). Previous studies have indicated that the dynamics of epigenetics markers during early embryo development are markedly affected by *in vitro* oocyte maturation and *in vitro* embryo culture in various species, including in mice, cows, pigs and humans (Abdalla et al., 2009; Cao et al., 2014; Heras et al., 2017; Lee et al., 2014; Maalouf et al., 2008; Nelissen et al., 2014; Salvaing et al., 2012; Santos et al., 2010). Of special relevance is the fact that metabolites present in the culture media can modulate the *in vitro* embryo development. Consequently, a lot of attention has been devoted to understanding embryonic metabolomics in the past decade (reviewed by Botros et al., 2008; Bracewell-Milnes et al., 2017; Krisher et al., 2015; Nel-Themaat and Nagy, 2011; Singh and Sinclair, 2007).

The importance of metabolites in regulating the cellular epigenome, inducing long-term changes to the cells, the so-called ‘metaboloepigenetic regulation’, has been reported in different cell types, such as stem cells and tumor cells (Donohoe and Bultman, 2012, reviewed by Etchegaray and Mostoslavsky, 2016; Reid et al., 2017; Van Winkle and Ryznar, 2019). More recently, studies have shown that epigenetic reprogramming might also be dependent on the metabolic pattern during the early stages of embryonic development (Ispada et al., 2020; Zhang et al., 2019). In this sense, understanding the metaboloepigenetic profile of *in vivo* embryos and how it differs from *in vitro* derived embryos in different species is essential to guiding future studies on modifying *in vitro* culture systems to improve embryo viability and progeny health. To this end, we analyzed RNAseq data of bovine, human and mouse *in vivo* and *in vitro* derived single oocytes (metaphase II) and early embryos (2-cell, 4-cell, 8-cell, 16-cell, morula and blastocyst) with emphasis on transcripts related to metabolic pathways that are known to influence cellular epigenetics.

## Results

### RNAseq data of bovine, human and mouse oocytes and embryos

Raw RNAseq data from oocytes (metaphase II) and embryos at different stages of development (2-cell, 4-cell, 8-cell, 16-cell, morula, and blastocyst) from *in vivo* and *in vitro* bovines and mice and from *in vitro* humans were obtained from the Gene Expression Omnibus data repository. To allow for interspecific comparisons, raw sequence data were annotated and gene expression quantified using the Galaxy by NetworkAnalyst 3.0 web browser using identical parameters across all species (Zhou et al., 2019). We then used Probabilistic Quotient Normalisation (PQN; Dieterle et al., 2006) to ensure the data were comparable across species and developmental stages. We identified a total of 16,207 bovine, 19,347 human and 22,941 mouse annotated genes in at least one developmental stage. 13,132 genes were common to all three species and 1,208, 3,747 and 7,520 genes were unique to bovines, humans, and mice, respectively (Figure 1a). Only the 13,132 genes detected in all three species were used in subsequent analyses.

**Figure 1.**
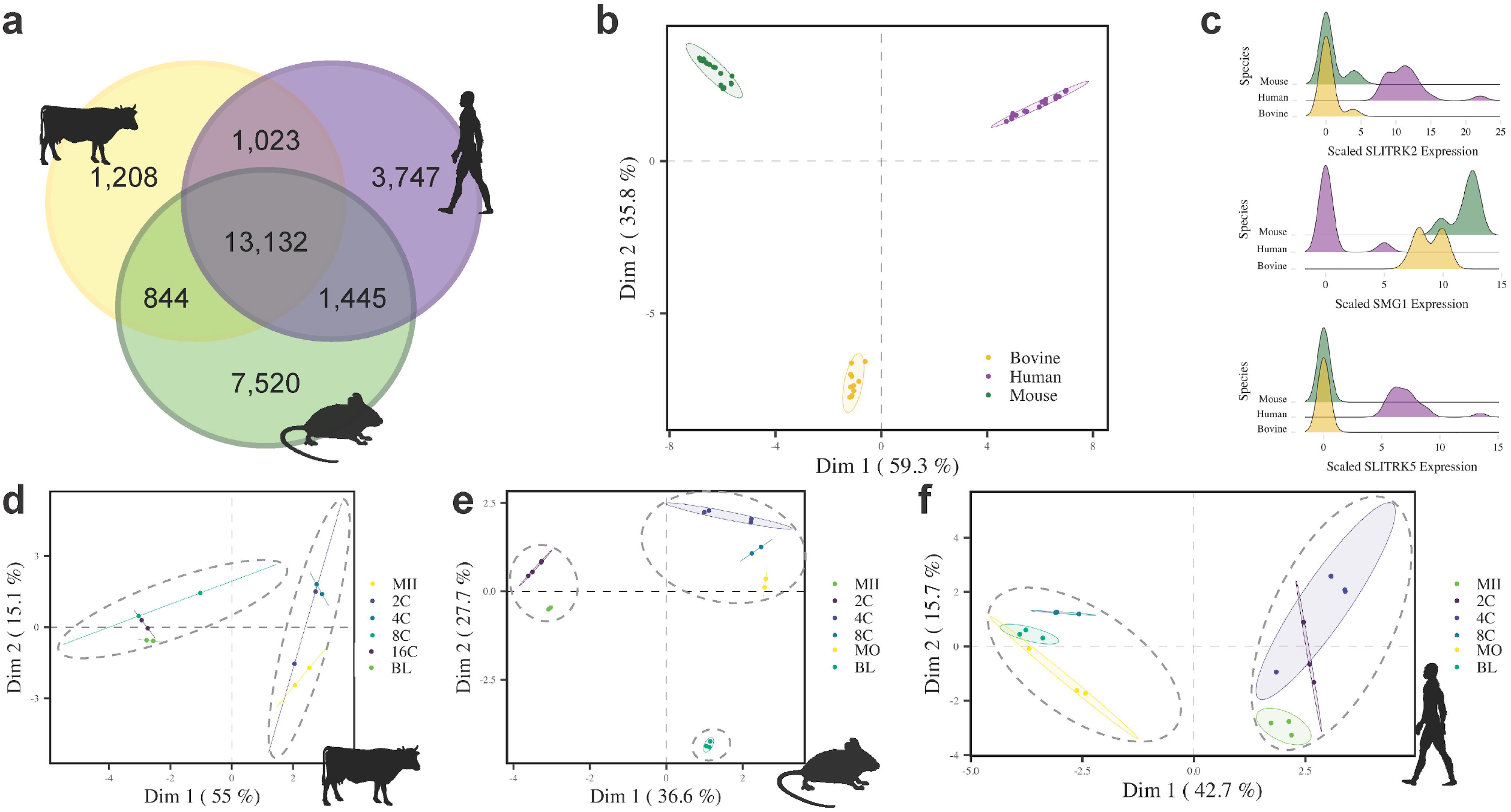
General analysis of *in vivo* bovine and mouse, and *in vitro* human mature oocytes (MII) and embryos (2C, 4C, 8C, 16C, MO and BL) gene expression. The Venn diagram in **a** shows patterns of overlap in genes expressed between bovines, humans and mice. The scatter plot in **b** depicts the first two dimensions (Dim) of a principal component analysis (PCA) across the proximity matrix of a random forest model classifying species based on gene expression profiles. Ellipses depict the means and covariances of the first two dimensions of the PCA for each species. In **c**, density plots of the genes of primary importance for classifying inter-specific variation are shown. PCAs of bovine (**d**), mouse (**e**) and human (**f**) gene expression across different developmental stages are also shown.

Since *in vivo* data from humans could not be obtained, we first compared *in vivo* samples of bovine and mouse and human *in vitro* samples (n = 50). Using these gene expression profiles, a random forest model classified species with an accuracy of 100% (Figure 1b and 1c). Principal component analyses (PCAs) showed clear patterns of (dis)similarities in gene expression across developmental stages for *in vivo* (bovine and mouse) and *in vitro* (human) oocyte and embryos (Figure 1d-f). These PCAs revealed two distinct clusters of *in vivo* bovine and *in vitro* human pre-implantation development: the first including the oocyte (MII), 2-cell (2C) and the 4-cell (4C) stages; and the second including the 8-cell (8C), the 16-cell (16C) or the morula (MO), and the blastocyst (BL) stages (Figures 1d and 1f). In mice, in contrast, *in vivo* pre-implantation development presented three distinct clusters: the first included the MII and the 2C stages; the second the 4C, the 8C and the MO stages; and the third the BL stage (Figure 1e). These results suggest that the embryo genome activation (EGA), occurs around the 4C-8C stage in bovine and human embryos and around the 2C-4C stage in mouse, which is in accordance with previous studies (Duan et al., 2019; Graf et al., 2014a; Guo et al., 2014; Jukam et al., 2017). Additionally, for humans, the transitions 4C-8C (11.94 %) and 8C-MO (10.05%) were the stages with most differentially expressed genes (DEG - Supplementary table 1 and Supplementary data 1). In bovine *in vivo* embryos, the highest number of DEG were observed in the 4C-8C (17.33%) and 16C-BL (3.56%) transitions. Mouse *in vivo* embryos had the highest number of DEG between MII-2C (26.53%), 2C-4C (29.77%, Supplementary table 1), and 16C-BL (41.15%).

### Embryonic metaboloepigenetic profiles are dynamic and distinct in humans, mice and bovines

Metabolism is a relevant aspect for reprogramming and epigenetic control of cells. With that in mind, we performed RNAseq analysis on the different pre-implantation development stages with focus on 117 metabolic and epigenetic pathways (Supplementary table 2) detectable in the different stages and species both *in vitro* and *in vivo*. The rotation gene set testing (ROAST) tool (Wu et al., 2010) was used to assess the significance of changes in these metabolic and epigenetic pathways as a unit (differentially expressed pathways – DEP; Supplementary data 2). In bovine *in vivo* samples, corroborating the EGA that happens around the 8C stage (Graf et al., 2014b; Jiang et al., 2014), the 4C-8C and 16C-BL transitions were the most variable (80 and 70% DEP, respectively; Figure 2 and Supplementary table 3). Similar to bovine, in *in vitro* human embryos, in which EGA occurs at 4C-8C stage (Niakan et al., 2012), the majority of DEP were detected at the transition 4C-8C, 8C-MO and MO-BL (78%, 84% and 65%, respectively; Figure 2 and Supplementary table 3). Mouse samples were the most distinct, with 91% DEP observed in the MII-2C transition and 97% in the transition MO-BL (Figure 2 and Supplementary table 3).

**Figure 2.**
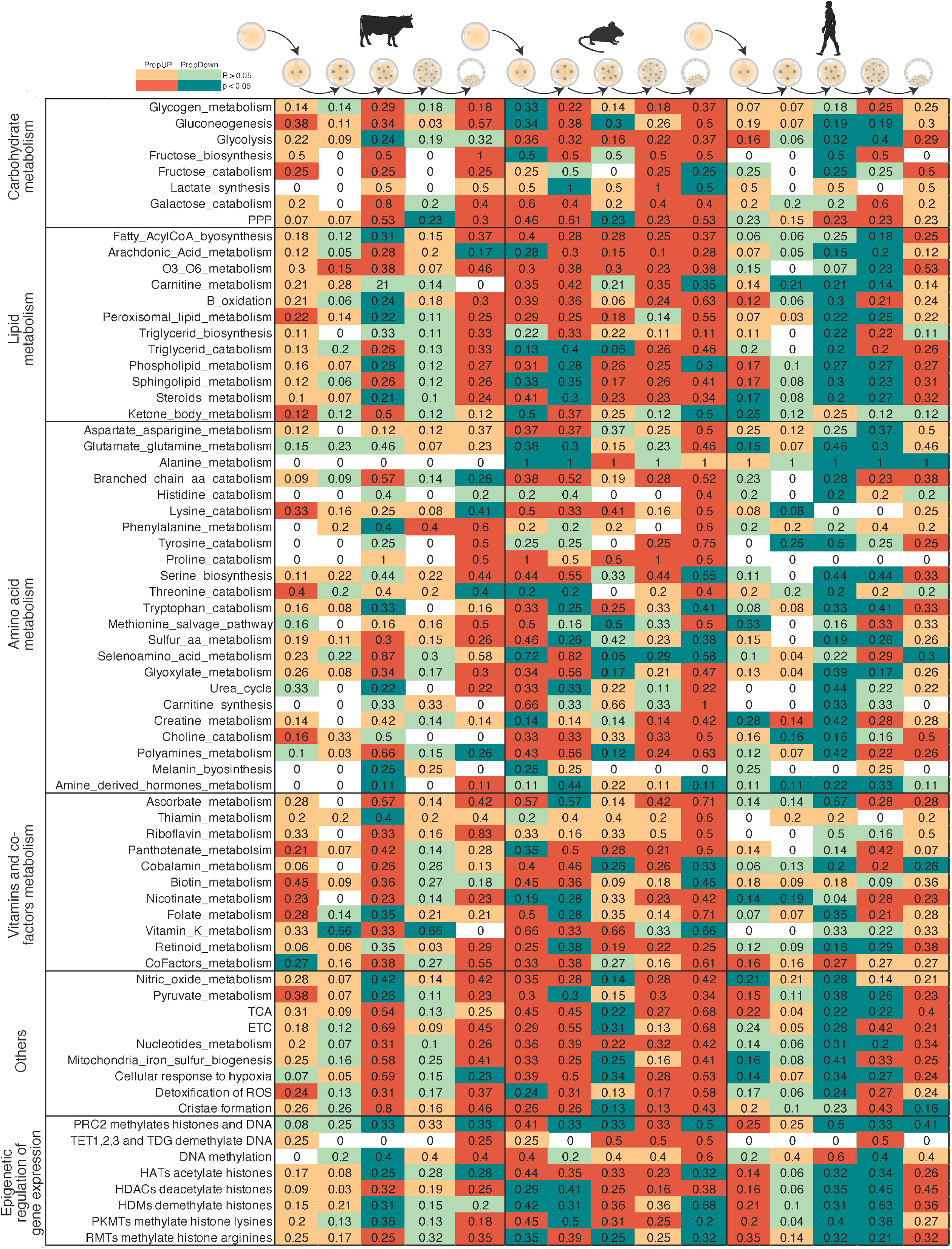
General analysis of human (*in vitro)*, mouse (*in vivo)* and bovine (*in vivo)* mature oocyte (MII) and embryos (2C, 4C, 8C, 16C, MO and BL), showing proportion of up (PropUp, red) and down (PropDown, green) regulated metabolic and epigenetic pathways (part of Reactome terms “Metabolism” and “Epigenetic regulation of gene expression”) in 2C compared to MII, 4C compared to 2C, 8C compared to 4C, 16C compared to 8C and BL compared to 16C. Differences were calculated using ROAST, and significance determined using a one-sided directional p-value < 0.05.

Close to the time of EGA and compaction in bovines and humans, embryos require more energy to support greater transcriptional activity, biosynthesis and cell proliferation, in addition to the blastocele formation and hatching, which marks the transition of an embryo composed of totipotent cells to a blastocyst containing pluripotent (inner cell mass – ICM) and differentiated (trophoblast – TB) cells. This increased metabolic demand can be seen by a change in the expression of the pentose phosphate pathway (PPP), glycolysis, beta-oxidation, electron transport chain (ETC, also known as oxidative phosphorylation) and the tricarboxylic acid cycle (TCA), which concurred with the reported changes in these metabolic pathways (Devreker, 2007; Gardner and Harvey, 2015; Guerif et al., 2013), specifically in the transition 4C-8C in bovine and humans and 2C-4C in mouse (Figure 3). Interestingly, in mice, this pattern was observed at MO-BL interval as well, which indicate that although DNA demethylation and major EGA occur at very early stages of development, the preferred metabolic pathways seem to follow the embryo’s functional requirement for compaction and differentiation, and not the molecular demand.

**Figure 3.**
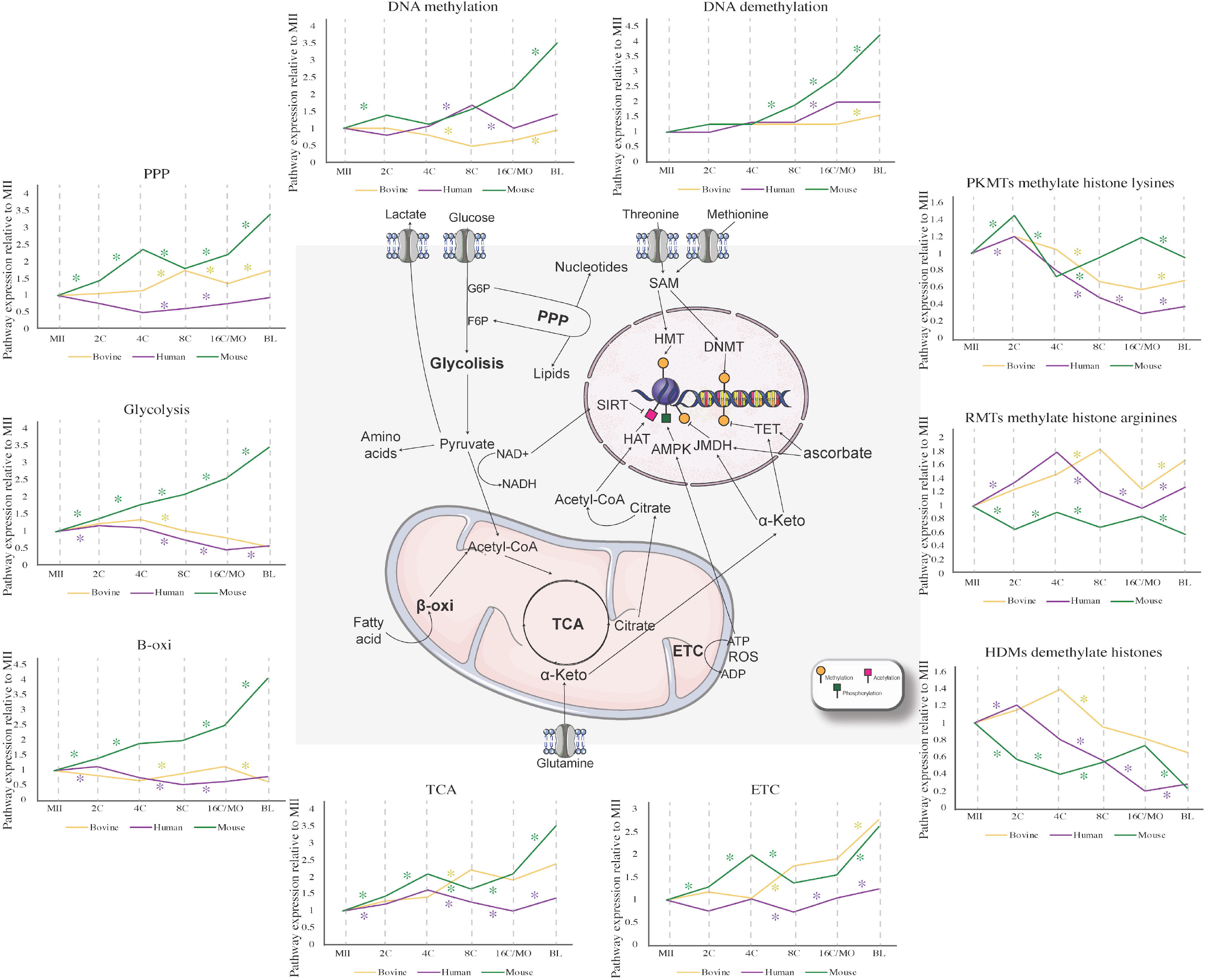
Differences on the metaboloepigenetic pathways profile of human (*in vitro)*, mouse (*in vivo)* and bovine (*in vivo)* pre-implantation embryos (2C, 4C, 8C, 16C/MO and BL). Known metaboloepigenetic pathways are depicted alongside each pathway expression pattern among the different pre-implantation stages (ROAST analysis data relative to MII expression) for human (purple), bovine (yellow) and mouse (green). Significant up and down regulated pathways between stages transition (MII-2C, 2C-4C, 4C-8C, 8C-16C/MO and 16C/MO-BL) intra-specie are marked with “*” (one-sided p-value < 0.05).

Beyond changes in metabolic requirements, pre-implantation embryos also undergo a wave of global epigenetic reprogramming that is in agreement with the dynamic expression of DNA and histone (de)methylation pathways observed (Figure 3). Such dynamics were slightly different between species and likely related to the moment of EGA. In humans, the DNA methylation pathway starts increasing at the 4C, reaching its higher expression at the 8C, which is then reduced at the MO and maintained through the BL (Figure 3), while the DNA demethylation pathway is increased at the MO only (Figure 3). In mice there was a significant increase in DNA methylation pathway expression at the 2C and at the BL stages (Figure 3). In contrast to humans, the mouse DNA demethylation pathway expression demonstrated a significant gradual increase from the 8C to the BL (Figure 3). In bovine a decrease on DNA methylation was observed at the 8C, which was followed by an increase at the BL stage. Bovine DNA demethylation pathway only increased at the BL stage (Figure 3).

It has been previously shown that the lowest levels of global DNA methylation occur around the 16C stage for humans, bovines and mice (Duan et al., 2019; Graf et al., 2014a; Guo et al., 2014; Jukam et al., 2017). More specifically, human embryos exhibit low levels of DNA methylation after the first cell division and remain that way until the MO stage, restoring the methylation levels only in the late BL stage (Guo et al., 2014). In this case, the minor EGA is also initiated during the first cleavage while the major EGA takes place around the 8C stage (Jukam et al., 2017). In bovines the process is similar, with demethylation being more evident from 4C and resumed earlier, still in the MO stage (Duan et al., 2019). As no differences in expression of the DNA demethylation pathway were observed between the 2C, 4C and 8C stages in neither humans nor bovines, nor at the 16C in bovines (Figure 3 and Supplementary data 2), it is clear that other players, such as metabolic factors, regulate such transitions in these species. As seen in Figure 3, the relationship between epigenetic events and metabolic aspects can occur at different levels within the cell.

Mouse embryos seemed to have the most distinct pattern. In this species, the global genome methylation is already low in 2C embryos in accordance with the major EGA, which also occurred at this stage. Unlike in humans and bovines, it was observed that mouse 8C, MO and BL stages have increased expression of the DNA demethylation pathway (Figure 3). This indicates that the action of demethylases was also responsible for such demethylation changes in mice. As for humans, *de novo* methylation will also be restored from the late BL (Jukam et al., 2017; Santos et al., 2002), which can be partially due to a significant increase on the expression of DNA methylation pathway at the BL stage in mice (Figure 3).

The dynamics of histone methylation are also crucial during gametogenesis and early embryogenesis, being the most abundant modification with the functional role dependent on the modified amino acid and the number of CH3 groups inserted (1, 2 or 3; Iwasaki et al., 2013; Ross et al., 2008; Sarmento, 2004; Torres-Padilla et al., 2007; Zhang et al., 2012). The insertion of CH3 groups leads to changes in the protein’s conformation, assembling or removing specific binding sites (Bannister and Kouzarides, 2011). For instance, H3K4 trimethylation (H3K4me3), primarily found around transcription initiate sites, is related to increased gene transcription and euchromatin formation (Sims et al., 2007; Wysocka et al., 2006), while H3K27 trimethylation (H3K27me3) leads to the opposite effect (Bártová et al., 2008). The patterns of H3K4me3 and H3K27me3 undergo dramatic changes during the initial embryogenesis, in accordance with its relevance for molecular events such as EGA and the establishment of totipotency. In humans, the levels of H3K4me3 and H3K27me3 decrease between 4C-8C and increase in the BL. H3K4me3, specifically, also increases after fertilization to the 4C (Zhang et al., 2012). In the same way, in bovine, both marks reach the lowest levels in the 4C-8C and reestablish in the BL (Ross et al., 2008; Wu et al., 2011). Mouse embryos present the most distinct pattern, with a reduction of H3K4me3 in the transition of 2C-4C and of H3K27me3 in the transition of 8C-MO and reestablishment of both marks only takes place in the ICM of the late BL (Wongtawan, 2010). In the present work, we observed a significant decrease in the expression of the histone demethylases (HDMs) pathway during the transition 4C-8C in bovine, while in humans this increased at MII-2C and at the MO-BL transitions and decreased at the 4C-8C and 8C-MO transitions (Figure 3). In mice, the increase was observed in the transition 4C-8C and decreased in MII-2C, 2C-4C and MO-BL transitions (Figure 3). Notably, it is clear from our findings that the embryonic metaboloepigenetic profile is a dynamic process, that varies not only inter-species, but also between the developmental stages intra-specifically.

To better elucidate these variations, we specifically characterized the patterns of transcripts belonging to methylation / demethylation of DNA and H3K4 and H3K27 residues, as well as the metabolic pathways known to be related to these processes: TCA cycle, one-carbon cycle (folate and methionine cycles) and methionine salvage pathway. The levels of these transcripts were analyzed in the 2C-MII, 4C-2C, 8C-4C, 16C-8C/MO-8C and BL-16C/BL-MO intervals in bovine and murine embryos produced *in vivo* and *in vitro* (except for mouse 2C-MII *in vivo*), as well as in the 2C-MII, 4C-2C, 8C-4C, MO-8C and BL-MO intervals in human embryos produced *in vitro*. Human embryos are not discussed in detail due to the limited variation between the different stages, which can be explained by the greater variability and molecular heterogeneity caused by morphophysiological and chromosomal aberrations characteristic of human embryonic samples, which are, normally, surplus of IVF cycles and/or discarded samples (Liu et al., 2009).

### Mouse embryo metaboloepigenetic profile is dynamic between stages and varies between in vivo and in vitro

The relationship between methylation events and metabolic aspects can occur at different levels within the cells. The first is related to the generation of methyl donors. Although DNA and histone methylation processes are catalyzed by different sets of enzymes, both use a common methyl donor, S-adenosylmethionine (SAM). Thus, changes in the intracellular availability of SAM can result in transcriptional changes throughout the genome (Stover and Caudill, 2008). The synthesis of SAM involves the one-carbon cycle metabolism which integrates two main pathways: the folate cycle and the methionine cycle (Figure 4). Briefly, folate can be converted to 5-methylTHF and integrated into the one-carbon cycle. 5-methylTHF is used as a scaffold to transport one-carbon units donated by the interconversion from serine to glycine, with serine being the largest donor of one-carbon in cells (Yang and Vousden, 2016). The resulting methyl group produced by the demethylation of 5-methyl THF is used to methylate the homocysteine, linking the folate cycle to the methionine cycle. Methionine is then adenylated by methionine adenosyl transferase (MAT) to form SAM, the main methyl group donor for epigenetic changes. SAM demethylation results in the formation of S-adenosylhomocysteine (SAH), which is hydrolyzed to homocysteine and can be remethylated to methionine and made available for a late SAM generation cycle (Clare et al., 2019; Stover and Caudill, 2008).

**Figure 4.**
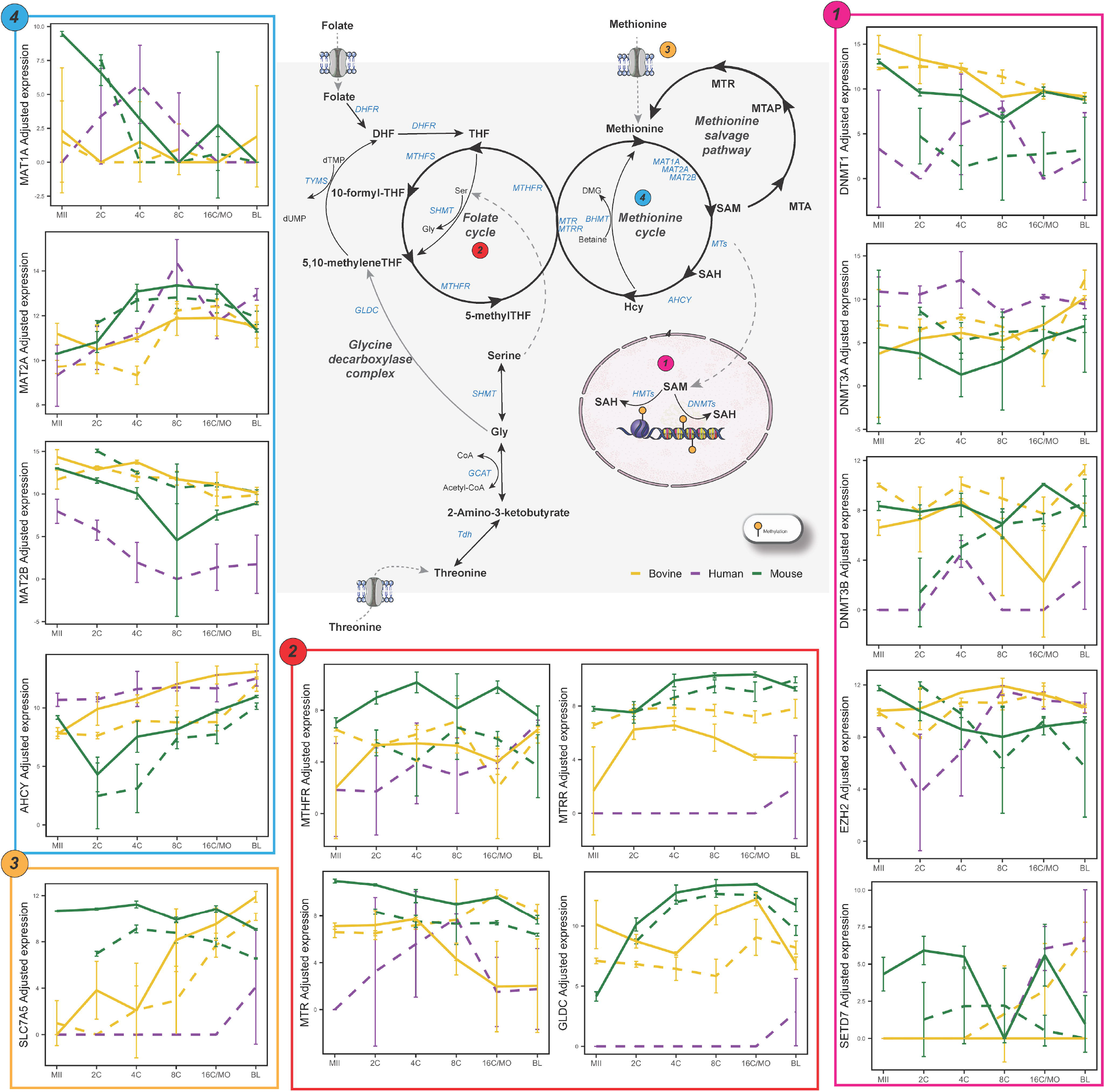
Metaboloepigenetic control of DNA and histone methylation of human (purple), mouse (green) and bovine (yellow) *in vivo* (full lines) and *in vitro* (dashed lines) pre-implantation stages (MII, 2C, 4C, 8C, 16C/MO and BL). Known metaboloepigenetic pathways are depicted alongside selected genes expression pattern among the different pre-implantation stages for each species. For significant up and down regulated genes between stages and species see Supplementary Figure 1.

The synthesis of SAM, therefore, integrates several metabolic aspects of the cell, in addition to external factors, among them vitamins as those of the B complex and, particularly, amino acids, as methionine, serine, glycine and threonine. The deficiency of glycine N-methyltransferase leads to overproduction of SAM, resulting in DNA hypermethylation and inappropriate gene silencing (Mudd et al., 2001). In tumor cells it has been shown that serine provides one-carbon units for the regeneration of methionine from homocysteine, supporting the methionine cycle even in conditions of methionine depletion in the culture media, somehow maintaining SAM generation and DNA methylation (Maddocks et al., 2016). In embryonic stem cells, the maintenance of pluripotency is dependent on H3K4me3 and this mark dramatically decreases when threonine is not available in the culture media, although DNA methylation and other lysine residues remain unchanged, suggesting that in these cells, threonine is metabolized to selectively promote H3K4me3 (Shyh-Chang et al., 2013). In addition, in mice, the restriction of methionine intake also triggers a decrease in H3K4 methylation, reinforcing the sensitive relationship between intracellular SAM levels and epigenetic control (Mentch et al., 2015) (Figure 4).

In addition to the relationship between metabolism and SAM generation, intermediates of metabolic pathways also interfere with the activity of enzymes responsible for DNA and histone demethylation (Figure 5). A key molecule in this process is α-ketoglutarate (α-KG). an intermediate in the tricarboxylic acid cycle. This metabolite can be generated by the deamination of glutamate by the enzyme glutamate dehydrogenase or, in the TCA cycle, by the decarboxylation of the isocitrate by the enzyme isocitrate dehydrogenase. In the TCA cycle, α-KG is decarboxylated in succinyl-CoA and CO_2_ by α-ketoglutarate dehydrogenase, which in turn is converted to succinate (Stryer et al., 2019). In embryonic stem cells the α-KG:succinate ratio affects pluripotency, and the accumulation of succinate and fumarate inhibits the enzymatic activity of TET demethylases, leading to higher levels of DNA methylation and, consequently, the maintenance of a more differentiated state. On the other hand, the high αKG:succinate ratio, results in greater TET activity and reduced DNA methylation, maintaining the less differentiated state. Similar to DNA demethylation, histone demethylase enzymes also use α-KG as a cofactor to remove methyl groups from histones residues. Although α-KG is crucial for histone demethylation, it has been shown that the accumulation of succinate within the cell may antagonize the activity of histone demethylases and promote cell differentiation from embryonic stem cells (Carey et al., 2015; Ispada et al., 2020; TeSlaa et al., 2016).

**Figure 5.**
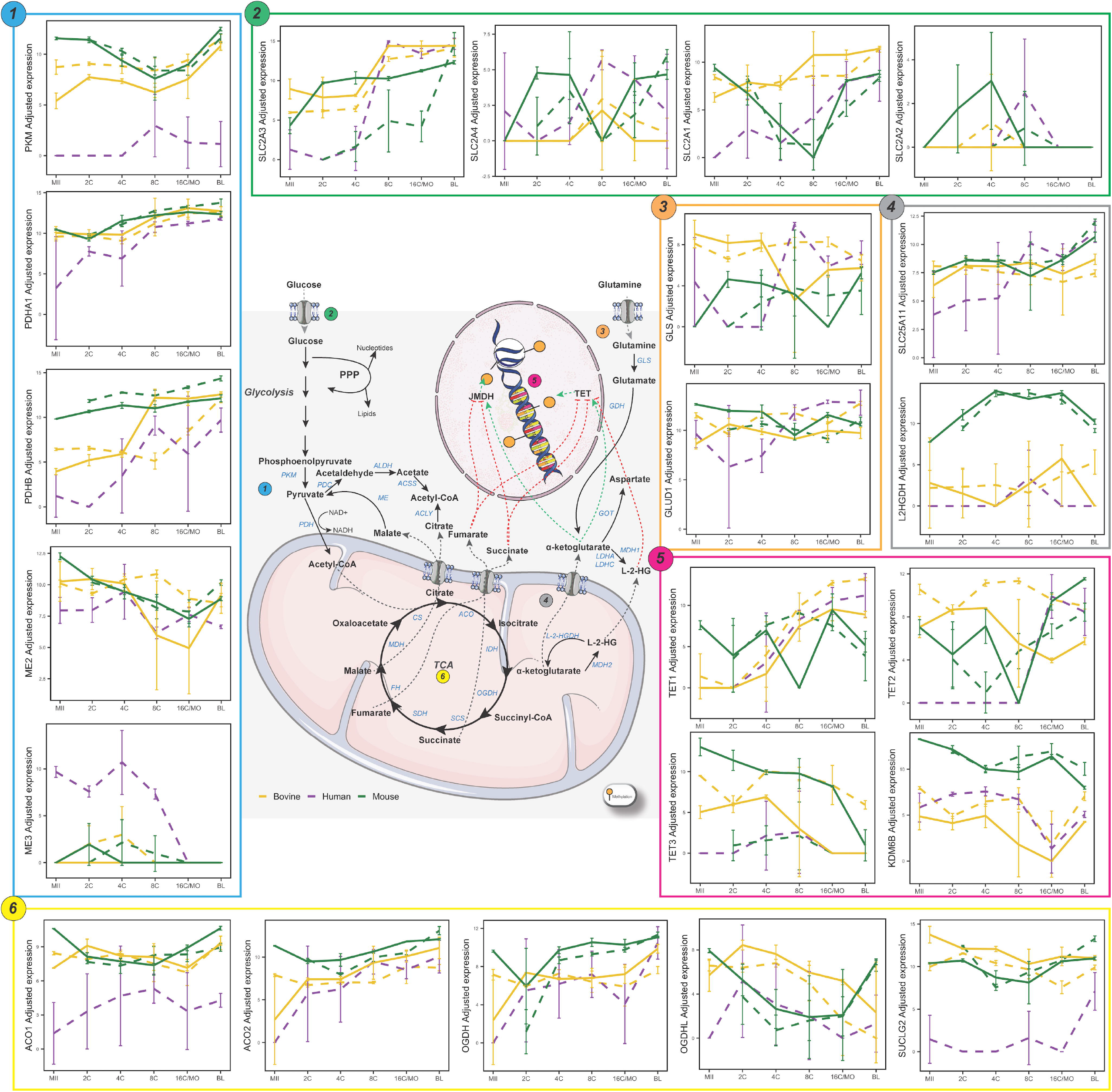
Metaboloepigenetic control of DNA and histone demethylation of human (purple), mouse (green) and bovine (yellow) *in vivo* (full lines) and *in vitro* (dashed lines) pre-implantation stages (MII, 2C, 4C, 8C, 16C/MO and BL). Known metaboloepigenetic pathways are depicted alongside each gene expression pattern among the different pre-implantation stages for each species. For significant up and down regulated genes between stages intra-species see Supplementary Figure 2.

Regarding these related pathways, in mice we verified marked differences in transcripts levels already at the 2C-MII interval, which can be explained by the major EGA that occurs around the 2C stage in this species (26.53%DEG *in vivo*). More specifically, at the beginning of development, *in vivo* embryos present a global scenario of demethylation, both for DNA and K4 (already in 2C) and K27 (from 8C) residues of histone 3. This was corroborated by the higher levels of *Kdm6a*, a H3K27 demethylase. On the other hand, *Tet* levels remained unchanged and there was a decrease in the levels of transcripts related to H3K4 demethylation (*Kdm5a* and *Kdm5b*) (Figure 5). Therefore, changes in both the availability of co-factors for demethylases and methyl donors could explain these demethylation processes *in vivo*. In fact, there was an increase in *L2hgdh*, a FAD-dependent enzyme that oxidizes L-2-hydroxyglutarate to α-KG, together with an increase in *Gls*, that catalyzes the conversion of glutamine in L-glutamate that can be further converted into α-KG, a co-factor of both TETs and JmjC-domain containing demethylases (Figure 5). A high α-KG:succinate ratio promotes DNA and histone demethylases activity, leading to chromatin modifications related to hyper-transcription in embryonic stem cells (Turner, 2008). This possible increase in demethylases activity would be accompanied by the lower availability of SAM, evidenced by a lower *Mtr, Mat1a* and *Mat1b* (Figure 4). The decline of *Mtr* decrease THR to SAM flux and, along with the decrease in *Mat1a* and *Mat1b*, there would be less generation of SAM, the main donor of methyl groups to methylations (Shyh-Chang et al., 2013).

During 4C-2C interval, there was still an intense dynamic in the synthesis of transcripts *in vitro* and *in vivo* (45.87 and 29.77 % DEG, respectively), which decreased in the 8C-4C and MO-8C intervals (*in vitro:* 7.22 and 4.68% and *in vivo*: 2.73 and 4.69%, respectively; Supplementary table 1) suggesting a transcriptional stability phase for these pathways, which underwent marked changes again in the MO-BL range (*in vitro: 31.99*% and *in vivo*: 41.15% DEG). During BL formation, *in vivo*, several histone and DNA methylation processes reinstate, preferentially in the cells of the ICM. Supporting the observed increase in methylation, there was, in general, a reduction in transcripts for demethylases and an increase in transcripts for histone and DNA methylases, except for the increase in *Tet2* (Figure 5). Among the TET enzymes, TET3 is the most involved in the global genome demethylation during the earliest stages of development, while TET1 and, mainly, TET2 participate in the maintenance of pluripotency in cells of ICM (Ito et al., 2010; Koh et al., 2011), explaining the greater levels in this interval (Figure 5). Both for *in vivo* and *in vitro* BL there was an increase in *Ahcy*, which converts SAH, an inhibitor of methyltransferases, to homocysteine, preventing its accumulation (Figure 4). It is worth noting that, in several tissues, such as the liver, it is not the greater availability of SAM that creates a supportive environment for methylation, but rather the SAM:SAH ratio, the methylation index (Hoffman et al., 1980). Thus, for both *in vivo* and *in vitro* embryos, higher levels of *Ahcy* might be decreasing intracellular SAH, providing an environment more prone to methylation. While in *in vivo* BL there was a decrease in *Mat2a* and an increase in *Mat2b*, in *in vitro* BL *Mat2a* remained unchanged and there was a decrease in *Mat2b*, favoring the synthesis of SAM, which could explain the hypermethylated DNA status of the latter in relation to the former (Wright et al., 2011).

### Prior to EGA bovine embryo metaboloepigenetic profiles remained stable in vivo but were divergent in vitro

In bovine embryos, active and passive DNA demethylation also occurs from the first cleavage, reaching its lowest levels between 8C-16C (Duan et al., 2019). Although the major EGA already occurs at 2C in mice, in bovines this occurs only around 8C (Graf et al., 2014a). As expected, there were no marked differences in the transcript levels in 2C-MII and 4C-2C intervals for *in vivo* embryos (0.69 and 0.26% DEG, respectively), and it was only in the 8C-4C interval that substantial differences could be identified (17.33% DEG; Supplementary table 1). The 16C-8C interval showed minimal variation (0.43% DEG), which remained in the BL-16C range (3.56% DEG).

In further contrast to mice, where differences in metaboloepigenetic transcripts were already observed in the early cleavage stages (2C-MII, 4C-2C and 8C-4C), differences were mainly observed when early stages (MII/2C/4C) were compared to the BL stages in bovines, agreeing with the lower differences in number of DEG in early cleavage stages (MII-8C) in this species. The DNA demethylation that remarkedly occurs until the 8-16C stage was confirmed by an increased expression of demethylation factors (*TET2* and *TET3*, Figure 5) and a decrease in the maintenance DNA methylase *DNMT1*, in early stages (MII and 4C) compared to the BL stage (Figure 4). Metabolically, a lower level of the methionine transport Solute Carrier Family 7 Member 5 (*SLC7A5*), was observed in early-stage embryos (MII, 2C and 4C) compared to BL (Figure 4), which would lead to lower methionine levels and, consequently, less SAM as substrate for DNA methylation.

Interestingly, *in vitro* embryos presented relevant changes in both the 2C-MII and 4C-2C intervals (35.18 and 34.66% DEG, respectively). More specifically, in the 2C-MII interval *in vitro*, several enzymes from the TCA cycle appear upregulated, with the exception of *ACO*, that catalyzes the isomerization of citrate to isocitrate, and *OGDH*, that catalyzes the oxidative decarboxylation of α-KG to Succinyl-CoA. *GLS* and *GLUD1*, which participate in the conversion of glutamine and glutamate to α-KG, were downregulated, as well as several histone and DNA methylases (Figures 4 and 5, Supplementary data 1). Surprisingly, in the 4C-2C interval *in vitro*, the global scenario was exactly the opposite, with TCA cycle enzymes being overexpressed, except *GLS, GLUD1* and *OGDHL*, while there was an overall increase in methylases and demethylases.

*In vitro* 16C-8C interval showed slightly variation (4.49% DEG), which increased in the BL-16C range (25.89% DEG). In the latter, there was a substantial increase in transcripts related to the TCA cycle. Still, there were lower levels of *MAT2A* and higher levels of *AHCY* (Figure 4), suggesting the lower synthesis of SAM, but still maintaining the favorable methylation environment, typical of this phase, which is corroborated by the increase in *DNMT3A* and *DNMT3B* (DNA de novo methylation; Figure 4 and Supplementary data 1). It is important to note that for H3K27, which also begins to be globally methylated in this phase *in vitro*, there was a decrease in *EZH2* methylase (Figure 4) and an increase in *KDM6B* demethylase (Figure 5), suggesting an additional control beyond the amount of enzymes for this methylation process. Furthermore, the decrease in *MAT2A* might explain the increase in hypomethylated genomic loci in embryos cultured *in vitro* when compared to *in vivo* (Salilew-Wondim et al., 2018).

Some important elements must be considered in relation to these results: i) there was an inverse relationship between the transcription of TCA cycle enzymes and those related to methylation/demethylation; ii) there was an increase in the levels of transcripts related to these pathways, which is not expected prior to the major EGA. The interpretation of this phenomenon in light of the metabolic and epigenomic changes that take place in this phase is quite challenging as the decrease in the amount of transcripts may be related to less transcription or greater translation. Nevertheless, the increase in transcripts levels prior to the major EGA for *in vitro* embryos and differently from *in vivo* embryos suggests a lack of molecular control with possible consequences to both the metabolic activity and the epigenome reprogramming. The earliest stages of development are extremely sensitive to the microenvironment. *In vivo*, this environment is highly influenced by the maternal state, while *in vitro*, the culture conditions have a remarkable impact on their development potential, impairing their developmental competence, and disturbing the subsequent maternal-embryonic communication, leading to failures in the establishment of pregnancy and long-term implications to the offspring (Krisher et al., 1999; Leroy et al., 2015; Lucy et al., 2014; Thompson et al., 1996; Velazquez, 2015).

This lack of molecular control may be a consequence of *in vitro* culture conditions which, despite presenting an acceptable relative efficiency, still report a high arrest rate up to 16C (Antunes et al., 2010). Thus, the inadequate supply of metabolites in culture media in supra physiological concentrations could require the embryo to activate metabolic pathways in advance, in an attempt to survive. In fact, when compared to oviduct and uterus fluids, SOFaa (Holm et al., 1999), a conventional culture medium for bovine embryos, offers lower amounts of glutamine but reaches five times more glutamate, three times more methionine, and almost twice more threonine (Hugentobler et al., 2007).

The results presented here point to differences in relation to the dynamics of transcript synthesis throughout development, characteristic of both species (mouse *vs* bovine) and microenvironment (*in vivo vs in vitro*). The latter, in particular, can be indicative of the molecular status of embryos and contribute to: i) the understanding of the effect of the environment on the metabolic, molecular and epigenomic status in the early stages of embryonic development; and ii) in the development of new strategies for assisted reproductive technologies (ART) protocols aimed at obtaining blastocysts that are increasingly similar to those *in vivo* for further transfer. Due to this importance, the impact of *in vitro* embryo production (IVP) on the embryonic metaboloepigenetic features were investigated in more detail.

### In vitro environment impairs the metaboloepigenetic profile of pre-implantation embryos

During pre-implantation development, both metabolism and epigenetic reprogramming are considered to be extremely sensitive to changes in environmental conditions, such as those imposed by IVP systems. To better verify the impact of *in vitro* conditions on the proper epigenome reprogramming, we anew assessed the levels of transcripts related to DNA/histone methylation/demethylation, as well as those from metabolic routes related to these modifications. This time, the levels of these transcripts were analyzed throughout the development in mouse and bovine embryos, with special attention to the blastocyst stage.

A clear distinction between *in vivo* and *in vitro* oocytes and embryos from different developmental stages was observed in both bovine and mice (Figure 6a and b, Supplementary Figure 3a and b). Unlike *in vivo*, *in vitro* bovine pre-implantation development presented three distinct segments: the first including the MII, 2C, 4C and 8C stages; the second including the 16C stage; and the third including the BL stage (Supplementary Figure 3a). Similar to *in vivo, in vitro* mouse pre-implantation development presented three distinct segments: the first including the 2C, the second including the 4C, 8C and MO stages, and the third including the BL stage (Supplementary Figure 3b). The *in vivo and in vitro* observed differences of bovine and mouse pre-implantation development was supported by a random forest model, which classified the gene expression of 71 samples (32 from bovine and 39 from mice) with an accuracy of 100% for bovine and 97.5% for mice. Model diagnostics revealed that interspecies variation in the expression of *NRSN2, TXN* and *GSPT1* were the variables of primary importance for differentiating between collection method (*in vitro vs in vivo*; Supplementary Figure 3c). In contrast to *in vivo samples, in vitro* bovine embryos had the highest number of DEG between MII-2C (35.18%), 2C-4C (34.66%) and 16C-BL (25.89%), while mouse *in vitro* had the highest number of DEG between 2C-4C (45.87%) and MO-BL (31.99%, Supplementary table 1). Considering developmental stages individually, *in vitro* produced bovine and mouse embryos had, in average, 4.47 and 9.58% DEG, respectively, compared to *in vivo* (Supplementary table 4). MII and BL were the stages with more DEG in bovine (5.61 and 5.79%, respectively), while in the mouse the 2C, MO and BL had the most DEG (11.77, 12.58 and 12.65%, respectively; Supplementary table 4).

**Figure 6.**
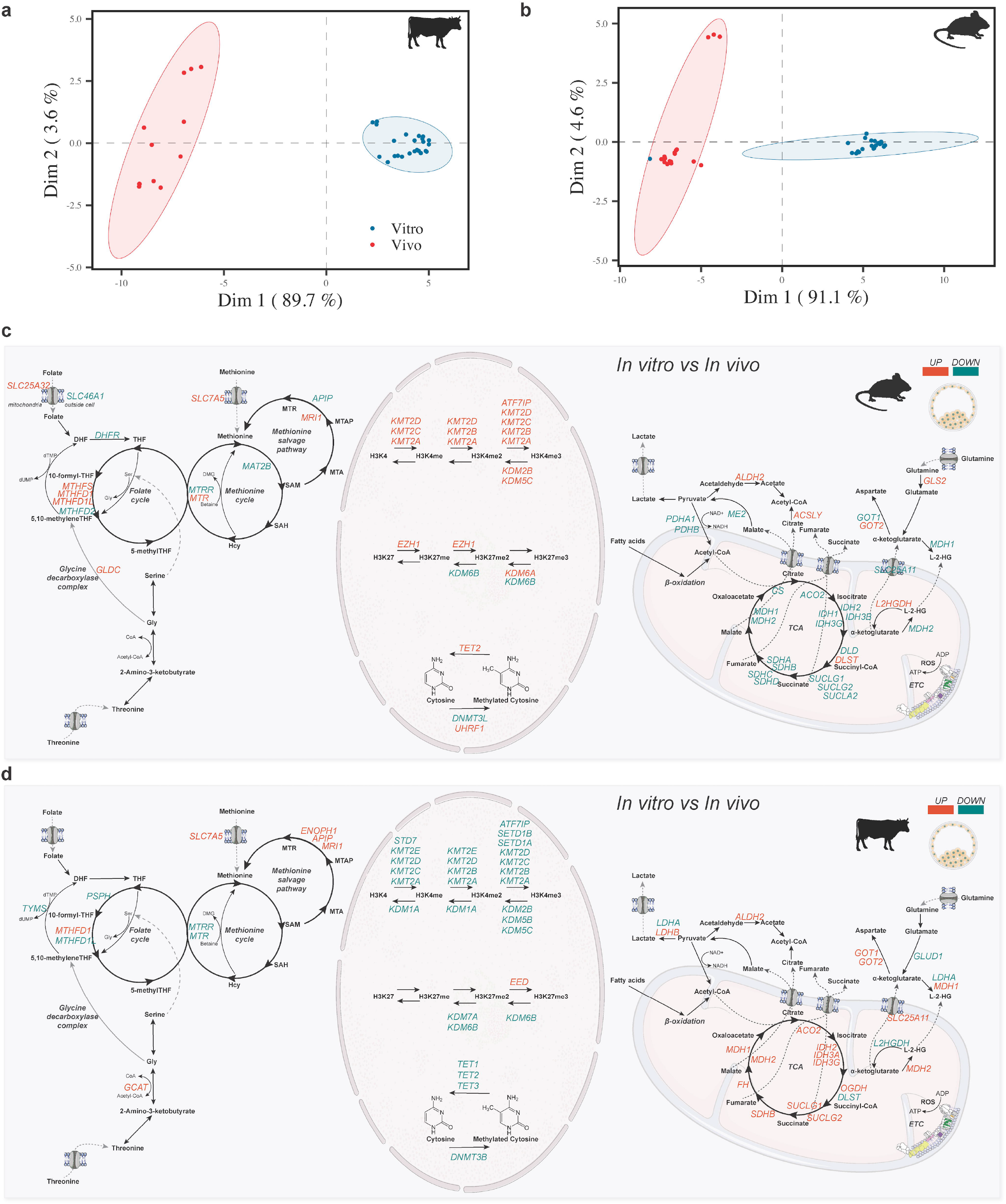
General analysis of *in vitro* bovine and mouse mature oocyte (MII) and embryos (2C, 4C, 8C, 16C, MO and BL) gene expression of metabolism and epigenetic genes. PCA comparing bovine (**a**) and mouse (**b**) *in vivo* and *in vitro* samples. Metaboloepigenetic genes up (red) and down (green) regulated (adjusted p-value < 0.05) comparing *in vitro vs in vivo* blastocysts from bovine (**c**) and mouse (**d**).

DEG for individual stages were mainly intricated in the metaboloepigenetic pathways analyzed (Figure 7), with bovine *in vitro* having 85.34% of DEP and mouse *in vitro* having 85.21% of DEP (Supplementary table 5). In the bovine, BL was the stage with the highest DEP (93.97%), while in mouse it was the 2C (96.52%; Supplementary table 5). Independent of the developmental stage, the majority of metabolic pathways that were up-regulated in bovine *in vitro*, were down regulated in mice *in vitro*: such as TCA, ETC, methionine salvage pathway, gluconeogenesis, beta-oxidation, glutamate and glutamine metabolism, pyruvate metabolism, among others (Supplementary Figure 4). In contrast, epigenetic pathways were mainly down regulated in bovine *in vitro* and up regulated in mouse *in vitro*, also independently of the developmental stage (Supplementary Figure 4). These results indicate that the *in vitro* environment has an important effect on the metaboloepigenetic control of bovine and mouse embryos. Moreover, it corroborates the fact that bovine and mouse embryos have different developmental kinetics and are, therefore, differentially affected by the *in vitro* environment.

**Figure 7.**
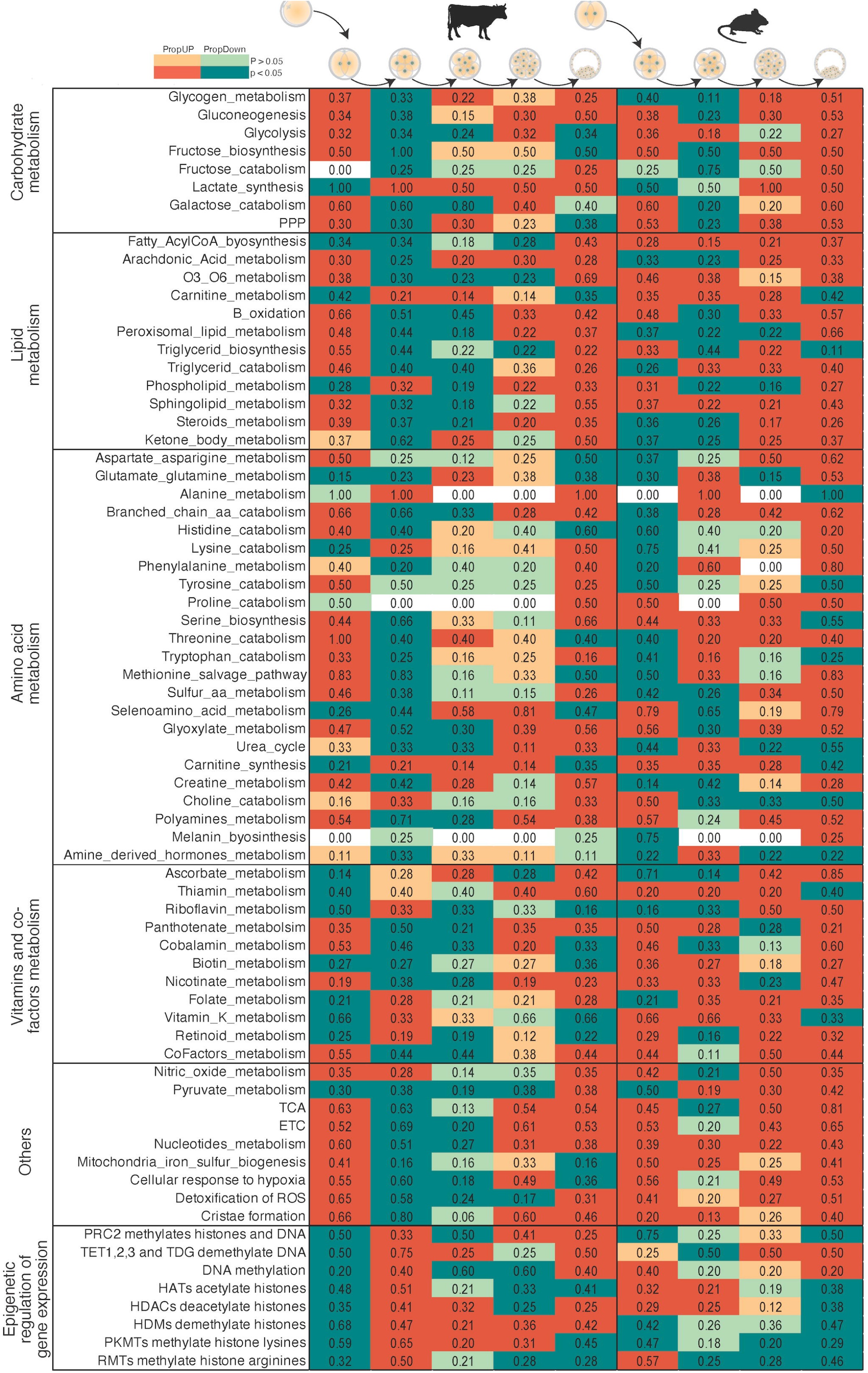
General analysis of *in vitro* bovine and mouse mature oocyte (MII) and embryos (2C, 4C, 8C, 16C, MO and BL), showing proportion of up (PropUp, red) and down (PropDown, green) regulated metabolic and epigenetic pathways (part of Reactome terms “Metabolism” and “Epigenetic regulation of gene expression”) in 2C compared to MII (bovine only), 4C compared to 2C, 8C compared to 4C, 16C compared to 8C and BL compared to 16C. Differences on PropUp and PropDown were calculated using ROAST, and significance determined using a the one-sided directional p-value < 0.05.

Mouse blastocysts produced *in vitro* also showed several metabolic indicators that confirmed their hypermethylated DNA status compared to blastocysts produced *in vivo* (Wright et al., 2011) since this changes was not explained by differences in transcripts related to methylases and demethylases (Figure 6c). Among them, we point out the highest amount of transcripts of the folate cycle and those connecting the folate cycle to methionine cycle, such as *Mthfs, Mthfd, Glcs* and *Mtr* (Figure 6c). In addition, *in vitro* embryos had higher levels of *Slc7a5*, a methionine transporter, and *Ahcy*, together with lower levels of *Mat2b*, reinforcing this environment more conducive to methylation (Figure 6c). In the case of bovine blastocyst, similarly, the differences in methylation reported in the literature were not explained by differences in the transcripts of methylases and demethylases (Figure 6d). In this species, however, unlike the mouse, there were lower levels of *MTR* and *MTRR*, despite the increase in *GCAT, SLC7A5* and transcripts of the methionine salvage pathway (Figure 6d). This scenario could indicate an active metabolism of methionine, however not necessarily related to methylation of DNA and histones. Methionine is a direct target of reactive oxygen species, acting as scavenger and protect cells from oxidative stress (Luo and Levine, 2009). ART protocols are known to induce higher oxidative stress since the gametes and embryos must be manipulated during maturation, fertilization and embryo development in environments that generate reactive oxygen species (ROS) (Torres-Osorio et al., 2019), which could lead to methionine deviation for ROS scavenger, decreasing the availability of SAM for methylation of DNA and histones and conferring a hypomethylated status to these embryos (Salilew-Wondim et al., 2018). This deviation to ROS scavenger can be substantiated by an increase in the metabolic pathway “detoxification of ROS” not only in *in vitro* produced BL, but also in MII, 4C, 8C and 16C embryos (Supplementary Figure 4).

We also observed a relationship between TCA cycle transcripts and those related to epigenetic events. In the case of *in vitro* mouse blastocysts, 16/20 transcripts from the TCA cycle were downregulated and 11/30 methylase and demethylase transcripts were upregulated when compared to *in vivo* ones (Figure 6c). It was also seen in a reduction of the metabolic pathway “TCA” in all developmental stages and in an increase in the epigenetic pathway “TET1, 2, and 3 and TDG demethylate DNA” and in the “DNA methylation” in all but the 8C stage (Supplementary Figure 4). In bovine blastocysts, this relationship was reversed; *in vitro* blastocysts had 12/20 transcripts from the TCA cycle upregulated and 19/30 methylase and demethylase transcripts downregulated when compared to their *in vivo* counterparts (Figure 6d). Oppositely to mouse, these changes could be corroborated by an increase in the metabolic pathway “TCA” in all but the 2C stage, and a reduction in the epigenetic pathway “TET1, 2, and 3 and TDG demethylate DNA” in all stages (Supplementary Figure 4). A comparison of the *in vivo vs in vitro* 2C, 4C, 8C and 16C/MO metaboloepigenetic genes for mice and bovines can be seen in Supplementary Figures 5 and 6, respectively.

These data point to metabolism, especially pathways related to the TCA cycle, as an unequivocal regulator of epigenetic processes in early embryos beyond the synthesis of succinate or α-KG, as already described for other cells and tissues. For instance, acetyl-CoA, generated from oxidation of pyruvate, fatty acid oxidation or amino acid degradation acts as an acetyl donor-group for histone acetylation, affecting global histone acetylation and gene expression, with possible consequences to the methylation status (Moussaieff et al., 2015). Also, 2-Hydroxyglutarate, an α-KG derived metabolite increased in hypoxic conditions, targets α-KG-dependent dioxygenases, inhibiting their function (Xu et al., 2011). Fumarate, another intermediate of TCA cycle, causes DNA hypermethylation by inhibiting TET activity (Sciacovelli et al., 2016). These data reinforce the environment in which embryos are produced as essential for proper metabolic control, which increasingly impose themselves as a determinant of the epigenomic control of the embryo.

## Discussion

As aptly stated by (Harvey et al., 2016), “metabolism is at the heart of cell-sensing mechanisms”. Beyond simply providing ATP to maintain homeostasis and cell replication, metabolism generates intermediate products that form the basic building blocks for cell proliferation, and modulate signalling pathways and gene expression (reviewed by Donohoe and Bultman, 2012; Harvey et al., 2016; Milazzotto et al., 2020; Spyrou et al., 2019). Further, metabolites have the capacity to regulate the cellular epigenome, inducing long-term changes to cells via a process known as metaboloepigenetic regulation (Donohoe and Bultman, 2012). Here we assembled the largest collection of embryonic RNAseq data compared to date in order to understand the molecular control of the metaboloepigenetic profile of bovine, mouse, and human *in vitro* and *in vivo* pre-implantation embryos. Our analyses have shown that the embryonic metaboloepigenetic profile is unique for each species, is highly dynamic between developmental stages, and is modified in *in vitro* cultured embryos.

The idea that the environment where oocytes and embryos develop may influence their outcome is certainly not new. Strategies to boost the numbers of transferable embryos by altering the availability of molecules in culture media in a search for the right formulations or culture conditions that mimic conditions found inside the reproductive tract have been an active area of research since the development of IVF 43 years ago (Cebrian-Serrano et al., 2013; Ferraz et al., 2018; Leese et al., 2008; Menezo et al., 2018; Sturmey et al., 2009). Despite decades of research, however, there have been only limited improvements in terms of developmental rates pre- and post-transfer (Simopoulou et al., 2018). Furthermore, although live births from ARTs (such as hormonal stimulation, embryo production by intracytoplasmic sperm injection and IVF) are now routine, ARTs are increasingly being shown to lead to postnatal ailments including premature births, delayed development, and metabolic disorders such as insulin resistance, and increased fasting blood glucose (Chen et al., 2014, 2013; Coussa et al., 2020). Consequently, understanding the molecular events that coordinate early embryonic development may assist in determining important pathways and processes for this moment, as well as identifying changes induced by ART (Menezo et al., 2018). Of special relevance is the fact that metabolites have been shown to modulate the *in vitro* embryo development, and a lot of attention has been devoted to understanding embryonic metabolomics in the past decade (reviewed by Botros et al., 2008; Bracewell-Milnes et al., 2017; Krisher et al., 2015; Nel-Themaat and Nagy, 2011; Singh and Sinclair, 2007). Interestingly, our findings have shown that, while the average number of DEG between *in vivo* and *in vitro* embryos were relatively low (4.47 and 9.58% for bovines and mice, respectively), these DEG were implicated in metaboloepigenetic pathways, accounting for an average difference of more than 80% DEP for both bovines and mice. These results demonstrate just how sensitive embryonic metabolism is to environmental conditions.

With these findings in mind, it is striking to consider that, even after four decades effort, most of the protocols for embryo culture still rely on a one-step supply of nutrients. Consequently, the dramatic differences we noted between the metaboloepigenetic profiles of *in vivo* and *in vitro* embryos was not overly surprising. As shown here, the embryonic metaboloepigenetic profile is stage dependent, which is not represented by current static IVP protocols. Moreover, our findings clearly show that the mechanisms leading to an incorrect epigenome reprogramming after IVF are under metabolic control. Thus, IVP protocols should also consider the dynamic nature of metabolic profiles in order to further improve their success. This is especially important given that cell metabolism, which has been remarkably affected trough changes in ART systems, seems to be the key in embryo epigenetic reprograming. To this end, and perhaps more important than accurately reproducing the physiological environment, we suggest that the focus should be on identifying systems and/or protocols that allow the embryos to respond the same way they would normally do *in vivo*. The embryonic metaboloepigenetic profiles mapped here are fundamental to developing new dynamic protocols that recreate the *in vivo* metaboloepigenetic profile, becoming a unique tool for the future of *in vitro* production of embryos and the birth of healthy offspring.

## Limitations of Study

For ethical and legal reasons, collecting *in vivo* human embryos is not an option. All of our analysis in humans were therefore based on *in vitro* embryos, which are known to be different from *in vivo*. Our RNAseq analysis was performed by curating available raw RNAseq data obtained from the Gene Expression Omnibus (GEO). As such, it is likely that there was noise in the data related to different embryo collection, culture, RNA extraction protocols, library preparation, and RNAseq analysis. Although this limitation was mostly overcome via our data normalization protocols, a fully controlled study would have ensured that this noise was minimized. Furthermore, while this represents the largest collection of embryos RNAseq data compared to date, the number of samples per cell stage within each species were limited. As such, small sample sizes limited the scope and scale of the analysis and our ability to accurately detect DEG. Therefore, our findings may include false negatives/positives for both DEG and DEP. Despite these limitations, the present results have an invaluable implication to the advances of the IVF field.

## Acknowledgements

MAMMF was supported by the Alexander von Humboldt Foundation in the framework of the Sofja Kovalevskaja Award endowed by the German Federal Ministry of Education and Research. MJN was supported by an NSERC Discovery Grant RGPIN-2021-02758.

## Author Contributions

Conceptualization, M.A.M.M.F.; Methodology, M.A.M.M.F., M.J.N, and M.P.M.; Investigation, M.J.N., M.P.M. and M.A.M.M.F.; Data curation, M.A.M.M.F. and M.J.N.; Writing – Original Draft, M.A.M.M.F. and M.P.M.; Writing – Review & Editing, M.A.M.M.F., M.P.M, and M.J.N.; Funding Acquisition, M.A.M.M.F.

## Methods

### Resource availability

#### Lead contact

Further information and requests for resources should be directed to and will be fulfilled by the lead contact, Marcia de Almeida Monteiro Melo Ferraz.

#### Materials availability

This study did not generate new unique reagents.

#### Data and code availability

*In vivo* and *in vitro* derived oocytes and pre-implantation embryos RNAseq raw data were obtained from Gene Expression Omnibus (GEO) accession numbers are listed in table 1. Oocyte and embryonic stages and number of samples are depicted in table 1.

**Table 1.**
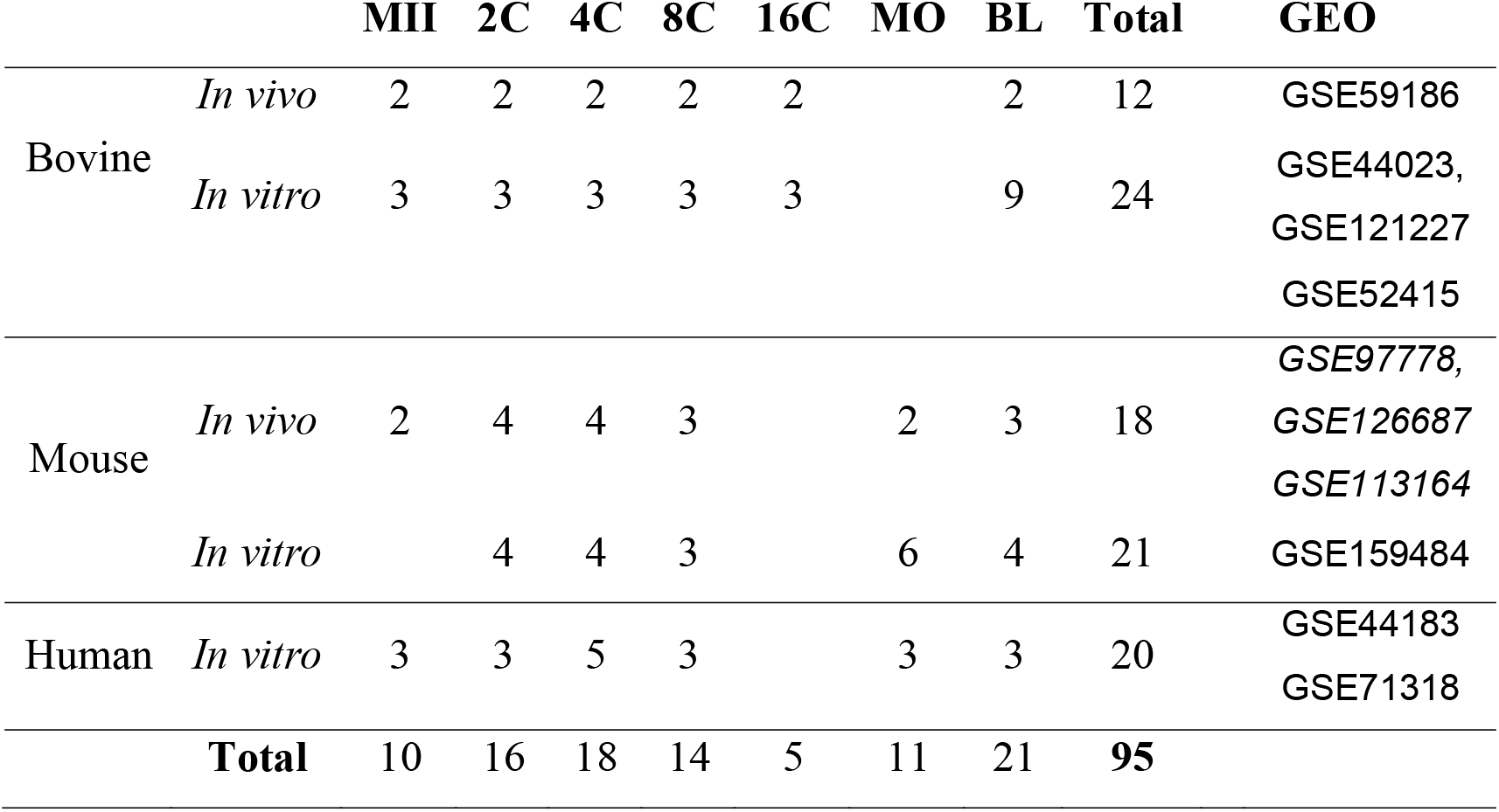
Oocyte and embryo developmental stage, method of collection and number of replicates.

This paper does not report original code and any additional information required to reanalyze the data reported in this paper is available from the lead contact upon request. All software used are freely available and are listed in the key resources table.

### Methods details

#### Data mining

RNAseq raw data were downloaded from Gene Expression Omnibus (GEO), accession numbers are described in the Key Resources table. Raw data was imported using Galaxy by NetworkAnalyst 3.0 web browser (https://galaxy.networkanalyst.ca/) (Zhou et al., 2019). Abundance of RNAseq transcripts was quantified using the function Kallisto (Bray et al., 2016). Genes with low counts (<5) were removed. For further analysis, only genes that were expressed in at least one stage of each species were used (13,132 genes).

#### RNAseq data normalization

Data were transformed using EdgeR: log2(CPM+c) (c = 4). These data were obtained from GEO, and, as such, it is likely that there was noise in the data related to different embryo collection, culture, and RNA extraction protocols, library preparation, and RNAseq analysis. To render the data comparable across species and developmental stages, EdgeR transformed data were scaled using Probabilistic Quotient Normalisation (PQN; (Dieterle et al., 2006)), which calibrates individual gene expression profiles against the median profile. Notably, analyses on PQN transformed data have been shown to have low false-positive rates, and can accurately recover groups of interest without introducing artefactual differences (Noonan et al., 2018).

#### Differentially expressed genes analysis

Differentially expressed genes (DEG) of normalized data were identified using the R package limma (Ritchie et al., 2015). An adjusted p-value to correct for multiple testing was calculated using the Benjamini–Hochberg method (Benjamini and Hochberg, 1995) that aims to control the false discovery rate across significant genes and is the most widely used correction for genomic studies. The R code necessary to reproduce these analyses is presented in Supplementary file S2.

#### Epigenetic and metabolic genes analysis using the rotation gene set testing

We accessed information of all epigenetic and metabolic pathways for humans (Reactome terms: “epigenetic regulation of gene expression”, “metabolism”, “metabolism of proteins” and “metabolism of RNA”) from the Reactome pathways database version 74 (Jassal et al., 2020). Reactome pathways are arranged into several tiers, the Reactome term “epigenetic regulation of gene expression” (Reactome ID: R-HSA-212165.2), included curated pathways involving 122 genes; the Reactome term “metabolism” (Reactome ID: R-HSA-1430728.10) involved 2,210 genes; the curated pathways of the Reactome term “metabolism of proteins” (Reactome ID: R-HSA-392499.7) involved 2,095 genes; and the curated pathways of the Reactome term “metabolism of RNA” (Reactome ID: R-HSA-8953854.4) involved 739 genes. Rotation gene set testing (ROAST) was used to perform self-contained gene set analysis of each metabolic pathway, for the different developmental stages and species (Wu et al., 2010). The ROAST analysis was performed using the mroast() function from the limma R package and the R code necessary to reproduce these analyses is presented in Supplementary file S2.

### Quantification and Statistical Analysis

Number of samples per species, collection method and stage used are depicted in table 1. For DEG, an adjusted p value < 0.05 was considered significant. For DEP, an one-sided directional p-value < 0.05 was considered significant.

A random forest (RF) model (Ho, 1995) was used to classify intra- and inter-species gene expression profiles according to developmental stages and collection method (*in vivo* or *in vitro*), with scaled gene expression values as the prediction variables. These analyses were conducted using the R package randomForest (ver. 4.6-14; RColorBrewer) (Liaw and Wiener, 2018). We chose RF modeling as it does not require any parameter reduction prior to analysis (Cutler et al., 2007), and has been shown to provide reliable results for biomarker identification (Chen et al., 2017), particularly on PQN transformed data (Noonan et al., 2018). We used the RF variable importance values to identify the key genes important for classifying groups of interest (i.e., species, collection type, developmental stage) in each RF model. The R code necessary to reproduce these analyses is presented in Supplementary file S2.

## Supplemental Information

Supplementary data 1: Log2 fold change, p-values and adjusted p-values for all differently expressed genes.

Supplementary data 2: Full ROAST analysis data.

Supplementary file S1: Supplementary tables 1-5 and Supplementary Figures 1-6.

Supplementary file S2: An HTML file detailing the R script required to reproduce the results presented in the main text.

## References

Abdalla, H., Hirabayashi, M., and Hochi, S. (2009). Demethylation dynamics of the paternal genome in pronuclear-stage bovine zygotes produced by in vitro fertilization and ooplasmic injection of freeze-thawed or freeze-dried spermatozoa. J. Reprod. Dev. 55, 433–439.

Antunes, G., Chaveiro, A., Santos, P., Marques, A., Jin, H., and Moreira da Silva, F. (2010). Influence of Apoptosis in Bovine Embryo’s Development. Reprod. Domest. Anim. 45, 26–32.

Bannister, A.J., and Kouzarides, T. (2011). Regulation of chromatin by histone modifications. Cell Res. 21, 381–395.

Bártová, E., Krejčí, J., Harničarová, A., Galiová, G., and Kozubek, S. (2008). Histone Modifications and Nuclear Architecture: A Review. J. Histochem. Cytochem. 56, 711–721.

Benjamini, Y., and Hochberg, Y. (1995). Controlling the False Discovery Rate: A Practical and Powerful Approach to Multiple Testing. J. R. Stat. Soc. Ser. B 57, 289–300.

Botros, L., Sakkas, D., and Seli, E. (2008). Metabolomics and its application for non-invasive embryo assessment in IVF. Mol. Hum. Reprod. 14, 679–690.

Bracewell-Milnes, T., Saso, S., Abdalla, H., Nikolau, D., Norman-Taylor, J., Johnson, M., Holmes, E., and Thum, M.Y. (2017). Metabolomics as a tool to identify biomarkers to predict and improve outcomes in reproductive medicine: A systematic review. Hum. Reprod. Update 23, 723–736.

Bray, N.L., Pimentel, H., Melsted, P., and Pachter, L. (2016). Near-optimal probabilistic RNA-seq quantification. Nat. Biotechnol. 34, 525–527.

Cao, Z., Zhou, N., Zhang, Y., Zhang, Y., Wu, R., Li, Y., Zhang, Y., and Li, N. (2014). Dynamic reprogramming of 5-hydroxymethylcytosine during early porcine embryogenesis. Theriogenology 81, 496–508.

Carey, B.W., Finley, L.W.S., Cross, J.R., Allis, C.D., and Thompson, C.B. (2015). Intracellular α-ketoglutarate maintains the pluripotency of embryonic stem cells. Nature 518, 413–416.

Cebrian-Serrano, a., Salvador, I., García-Roselló, E., Pericuesta, E., Pérez-Cerezales, S., Gutierrez-Adán, a., Coy, P., and Silvestre, M. a. (2013). Effect of the bovine oviductal fluid on in vitro fertilization, development and gene expression of in vitro-produced bovine blastocysts. Reprod. Domest. Anim. 48, 331–338.

Chen, J., Zhang, P., Lv, M., Guo, H., Huang, Y., Zhang, Z., and Xu, F. (2017). Influences of Normalization Method on Biomarker Discovery in Gas Chromatography–Mass Spectrometry-Based Untargeted Metabolomics: What Should Be Considered? Anal. Chem. 89, 5342–5348.

Chen, M., Wu, L., Wu, F., Wittert, G.A., Norman, R.J., Robker, R.L., and Heilbronn, L.K. (2014). Impaired Glucose Metabolism in Response to High Fat Diet in Female Mice Conceived by In Vitro Fertilization (IVF) or Ovarian Stimulation Alone. PLoS One 9, e113155.

Chen, Z., Robbins, K.M., Wells, K.D., and Rivera, R.M. (2013). Large offspring syndrome. Epigenetics 8, 591–601.

Clare, C.E., Brassington, A.H., Kwong, W.Y., and Sinclair, K.D. (2019). One-Carbon Metabolism: Linking Nutritional Biochemistry to Epigenetic Programming of Long-Term Development. Annu. Rev. Anim. Biosci. 7, 263–287.

Coussa, A., Hasan, H.A., and Barber, T.M. (2020). Impact of contraception and IVF hormones on metabolic, endocrine, and inflammatory status. J. Assist. Reprod. Genet. 37, 1267–1272.

Cutler, D.R., Edwards, T.C., Beard, K.H., Cutler, A., Hess, K.T., Gibson, J., and Lawler, J.J. (2007). RANDOM FORESTS FOR CLASSIFICATION IN ECOLOGY. Ecology 88, 2783–2792.

Devreker, F. (2007). Uptake and release of metabolites in human preimplantation embryos. In Human Preimplantation Embryo Selection, J. Cohen, and K. Elder, eds. (CRC Press), pp. 159–168.

Dieterle, F., Ross, A., Schlotterbeck, G., and Senn, H. (2006). Probabilistic Quotient Normalization as Robust Method to Account for Dilution of Complex Biological Mixtures. Application in 1 H NMR Metabonomics. Anal. Chem. 78, 4281–4290.

Donohoe, D.R., and Bultman, S.J. (2012). Metaboloepigenetics: Interrelationships between energy metabolism and epigenetic control of gene expression. J. Cell. Physiol. 227, 3169–3177.

Duan, J.E., Jiang, Z.C., Alqahtani, F., Mandoiu, I., Dong, H., Zheng, X., Marjani, S.L., Chen, J., and Tian, X.C. (2019). Methylome Dynamics of Bovine Gametes and in vivo Early Embryos. Front. Genet. 10, 1–14.

Etchegaray, J.-P., and Mostoslavsky, R. (2016). Interplay between Metabolism and Epigenetics: A Nuclear Adaptation to Environmental Changes. Mol. Cell 62, 695–711.

Ferraz, M.A.M.M., Rho, H.S., Hemerich, D., Henning, H.H.W., van Tol, H.T.A., Hölker, M., Besenfelder, U., Mokry, M., Vos, P.L.A.M., Stout, T.A.E., et al. (2018). An oviduct-on-a-chip provides an enhanced in vitro environment for zygote genome reprogramming. Nat. Commun. 9, 4934.

Gardner, D.K., and Harvey, A.J. (2015). Blastocyst metabolism. Reprod. Fertil. Dev. 27, 638.

Graf, A., Krebs, S., Heininen-Brown, M., Zakhartchenko, V., Blum, H., and Wolf, E. (2014a). Genome activation in bovine embryos: Review of the literature and new insights from RNA sequencing experiments. Anim. Reprod. Sci. 149, 46–58.

Graf, A., Krebs, S., Zakhartchenko, V., Schwalb, B., Blum, H., and Wolf, E. (2014b). Fine mapping of genome activation in bovine embryos by RNA sequencing. Proc. Natl. Acad. Sci. U. S. A. 111, 4139–4144.

Guerif, F., McKeegan, P., Leese, H.J., and Sturmey, R.G. (2013). A Simple Approach for COnsumption and RElease (CORE) Analysis of Metabolic Activity in Single Mammalian Embryos. PLoS One 8, e67834.

Guo, H., Zhu, P., Yan, L., Li, R., Hu, B., Lian, Y., Yan, J., Ren, X., Lin, S., Li, J., et al. (2014). The DNA methylation landscape of human early embryos. Nature 511, 606–610.

El Hajj, N., and Haaf, T. (2013). Epigenetic disturbances in in vitro cultured gametes and embryos: Implications for human assisted reproduction. Fertil. Steril. 99, 632–641.

Harvey, A.J., Rathjen, J., and Gardner, D.K. (2016). Metaboloepigenetic regulation of pluripotent stem cells. Stem Cells Int. 2016.

Heras, S., Smits, K., De Schauwer, C., and Van Soom, A. (2017). Dynamics of 5-methylcytosine and 5-hydroxymethylcytosine during pronuclear development in equine zygotes produced by ICSI. Epigenetics Chromatin 10, 13.

Ho, T.K. (1995). Random decision forests. In Proceedings of 3rd International Conference on Document Analysis and Recognition, (IEEE Comput. Soc. Press), pp. 278–282.

Hoffman, D.R., Marion, D.W., Cornatzer, W.E., and Duerre, J.A. (1980). S-Adenosylmethionine and S-adenosylhomocystein metabolism in isolated rat liver. Effects of L-methionine, L-homocystein, and adenosine. J. Biol. Chem. 255, 10822–10827.

Holm, P., Booth, P.J., Schmidt, M.H., Greve, T., and Callesen, H. (1999). High bovine blastocyst development in a static in vitro production system using sofaa medium supplemented with sodium citrate and myo-inositol with or without serum-proteins. Theriogenology 52, 683–700.

Hugentobler, S.A., Diskin, M.G., Leese, H.J., Humpherson, P.G., Watson, T., Sreenan, J.M., and Morris, D.G. (2007). Amino acids in oviduct and uterine fluid and blood plasma during the estrous cycle in the bovine. Mol. Reprod. Dev. 74, 445–454.

Ispada, J., da Fonseca Junior, A.M., de Lima, C.B., dos Santos, E.C., Fontes, P.K., Nogueira, M.F.G., da Silva, V.L., Almeida, F.N., Leite, S. de C., Chitwood, J.L., et al. (2020). Tricarboxylic Acid Cycle Metabolites as Mediators of DNA Methylation Reprogramming in Bovine Preimplantation Embryos. Int. J. Mol. Sci. 21, 6868.

Ito, S., D’Alessio, A.C., Taranova, O. V., Hong, K., Sowers, L.C., and Zhang, Y. (2010). Role of Tet proteins in 5mC to 5hmC conversion, ES-cell self-renewal and inner cell mass specification. Nature 466, 1129–1133.

Iwasaki, W., Miya, Y., Horikoshi, N., Osakabe, A., Taguchi, H., Tachiwana, H., Shibata, T., Kagawa, W., and Kurumizaka, H. (2013). Contribution of histone N-terminal tails to the structure and stability of nucleosomes. FEBS Open Bio 3, 363–369.

Jansz, N. (2019). DNA methylation dynamics at transposable elements in mammals. Essays Biochem. 63, 677–689.

Jassal, B., Matthews, L., Viteri, G., Gong, C., Lorente, P., Fabregat, A., Sidiropoulos, K., Cook, J., Gillespie, M., Haw, R., et al. (2020). The reactome pathway knowledgebase. Nucleic Acids Res. 48, D498–D503.

Jiang, Z., Sun, J., Dong, H., Luo, O., Zheng, X., Obergfell, C., Tang, Y., Bi, J., O’Neill, R., Ruan, Y., et al. (2014). Transcriptional profiles of bovine in vivo pre-implantation development. BMC Genomics 15, 756.

Jukam, D., Shariati, S.A.M., and Skotheim, J.M. (2017). Zygotic Genome Activation in Vertebrates. Dev. Cell 42, 316–332.

Koh, K.P., Yabuuchi, A., Rao, S., Huang, Y., Cunniff, K., Nardone, J., Laiho, A., Tahiliani, M., Sommer, C.A., Mostoslavsky, G., et al. (2011). Tet1 and Tet2 Regulate 5-Hydroxymethylcytosine Production and Cell Lineage Specification in Mouse Embryonic Stem Cells. Cell Stem Cell 8, 200–213.

Krisher, R.L., Lane, M., and Bavister, B.D. (1999). Developmental Competence and Metabolism of Bovine Embryos Cultured in Semi-Defined and Defined Culture Media1. Biol. Reprod. 60, 1345–1352.

Krisher, R.L., Schoolcraft, W.B., and Katz-Jaffe, M.G. (2015). Omics as a window to view embryo viability. Fertil. Steril. 103, 333–341.

Lee, K., Hamm, J., Whitworth, K., Spate, L., Park, K. wook, Murphy, C.N., and Prather, R.S. (2014). Dynamics of TET family expression in porcine preimplantation embryos is related to zygotic genome activation and required for the maintenance of NANOG. Dev. Biol. 386, 86–95.

Leese, H.J., Hugentobler, S. a, Gray, S.M., Morris, D.G., Sturmey, R.G., Whitear, S.L., and Sreenan, J.M. (2008). Female reproductive tract fluids: composition, mechanism of formation and potential role in the developmental origins of health and disease. Reprod Fertil Dev 20, 1–8.

Leroy, J.L.M.R., Valckx, S.D.M., Jordaens, L., De Bie, J., Desmet, K.L.J., Van Hoeck, V., Britt, J.H., Marei, W.F., and Bols, P.E.J. (2015). Nutrition and maternal metabolic health in relation to oocyte and embryo quality: critical views on what we learned from the dairy cow model. Reprod. Fertil. Dev. 27, 693.

Li, Y., and Sasaki, H. (2011). Genomic imprinting in mammals: its life cycle, molecular mechanisms and reprogramming. Cell Res. 21, 466–473.

Liaw, A., and Wiener, M. (2018). Breiman and Cutler’s Random Forests for Classification and Regression. CRAN Repos.

Liu, W., Yin, Y., Long, X., Luo, Y., Jiang, Y., Zhang, W., Du, H., Li, S., Zheng, Y., Li, Q., et al. (2009). Derivation and characterization of human embryonic stem cell lines from poor quality embryos. J. Genet. Genomics 36, 229–239.

Lucy, M.C., Butler, S.T., and Garverick, H.A. (2014). Endocrine and metabolic mechanisms linking postpartum glucose with early embryonic and foetal development in dairy cows. Animal 8, 82–90.

Luo, S., and Levine, R.L. (2009). Methionine in proteins defends against oxidative stress. FASEB J. 23, 464–472.

Maalouf, W.E., Alberio, R., and Campbell, K.H.S. (2008). Differential acetylation of histone H4 lysine during development of in vitro fertilized, cloned and parthenogenetically activated bovine embryos. Epigenetics 3, 199–209.

Maddocks, O.D.K., Labuschagne, C.F., Adams, P.D., and Vousden, K.H. (2016). Serine Metabolism Supports the Methionine Cycle and DNA/RNA Methylation through De Novo ATP Synthesis in Cancer Cells. Mol. Cell 61, 210–221.

Menezo, Y., Dale, B., and Elder, K. (2018). Time to re-evaluate ART protocols in the light of advances in knowledge about methylation and epigenetics: an opinion paper. Hum. Fertil. 21, 156–162.

Ménézo, Y., Guérin, P., and Elder, K. (2015). The oviduct: A neglected organ due for re-assessment in IVF. Reprod. Biomed. Online 30, 233–240.

Mentch, S.J., Mehrmohamadi, M., Huang, L., Liu, X., Gupta, D., Mattocks, D., Gómez Padilla, P., Ables, G., Bamman, M.M., Thalacker-Mercer, A.E., et al. (2015). Histone Methylation Dynamics and Gene Regulation Occur through the Sensing of One-Carbon Metabolism. Cell Metab. 22, 861–873.

Milazzotto, M.P., Lima, C.B. De, Fonseca Junior, A.M. da, Santos, E.C. dos, and Ispada, J. (2020). Erasing gametes to write blastocysts: metabolism as the new player in epigenetic reprogramming. Anim. Reprod. 17, 1–23.

Moussaieff, A., Rouleau, M., Kitsberg, D., Cohen, M., Levy, G., Barasch, D., Nemirovski, A., Shen-Orr, S., Laevsky, I., Amit, M., et al. (2015). Glycolysis-Mediated Changes in Acetyl-CoA and Histone Acetylation Control the Early Differentiation of Embryonic Stem Cells. Cell Metab. 21, 392–402.

Mudd, S.H., Cerone, R., Schiaffino, M.C., Fantasia, A.R., Minniti, G., Caruso, U., Lorini, R., Watkins, D., Matiaszuk, N., Rosenblatt, D.S., et al. (2001). Glycine N -methyltransferase deficiency: A novel inborn error causing persistent isolated hypermethioninaemia. J. Inherit. Metab. Dis. 24, 448–464.

Nel-Themaat, L., and Nagy, Z.P. (2011). A review of the promises and pitfalls of oocyte and embryo metabolomics. Placenta 32, S257–S263.

Nelissen, E.C.M., Dumoulin, J.C.M., Busato, F., Ponger, L., Eijssen, L.M., Evers, J.L.H., Tost, J., and van Montfoort, A.P.A. (2014). Altered gene expression in human placentas after IVF/ICSI. Hum. Reprod. 29, 2821–2831.

Niakan, K.K., Han, J., Pedersen, R.A., Simon, C., and Pera, R.A.R. (2012). Human pre-implantation embryo development. Development 139, 829–841.

Noonan, M.J., Tinnesand, H. V., and Buesching, C.D. (2018). Normalizing Gas-Chromatography-Mass Spectrometry Data: Method Choice can Alter Biological Inference. BioEssays 40, 1700210.

Reid, M.A., Dai, Z., and Locasale, J.W. (2017). The impact of cellular metabolism on chromatin dynamics and epigenetics. Nat. Cell Biol. 19, 1298–1306.

Ritchie, M.E., Phipson, B., Wu, D., Hu, Y., Law, C.W., Shi, W., and Smyth, G.K. (2015). limma powers differential expression analyses for RNA-sequencing and microarray studies. Nucleic Acids Res. 43, e47–e47.

Ross, P.J., Ragina, N.P., Rodriguez, R.M., Iager, A.E., Siripattarapravat, K., Lopez-Corrales, N., and Cibelli, J.B. (2008). Polycomb gene expression and histone H3 lysine 27 trimethylation changes during bovine preimplantation development. REPRODUCTION 136, 777–785.

Salilew-Wondim, D., Saeed-Zidane, M., Hoelker, M., Gebremedhn, S., Poirier, M., Pandey, H.O., Tholen, E., Neuhoff, C., Held, E., Besenfelder, U., et al. (2018). Genome-wide DNA methylation patterns of bovine blastocysts derived from in vivo embryos subjected to in vitro culture before, during or after embryonic genome activation. BMC Genomics 19, 424.

Salvaing, J., Aguirre-Lavin, T., Boulesteix, C., Lehmann, G., Debey, P., and Beaujean, N. (2012). 5-Methylcytosine and 5-Hydroxymethylcytosine Spatiotemporal Profiles in the Mouse Zygote. PLoS One 7, 20–23.

Santos, F., Hendrich, B., Reik, W., and Dean, W. (2002). Dynamic Reprogramming of DNA Methylation in the Early Mouse Embryo. Dev. Biol. 241, 172–182.

Santos, F., Hyslop, L., Stojkovic, P., Leary, C., Murdoch, A., Reik, W., Stojkovic, M., Herbert, M., and Dean, W. (2010). Evaluation of epigenetic marks in human embryos derived from IVF and ICSI. Hum. Reprod. 25, 2387–2395.

Sarmento, O.F. (2004). Dynamic alterations of specific histone modifications during early murine development. J. Cell Sci. 117, 4449–4459.

Sarmento, O.F., Digilio, L.C., Wang, Y., Perlin, J., Herr, J.C., Allis, C.D., and Coonrod, S.A. (2004). Dynamic alterations of specific histone modifications during early murine development. J. Cell Sci. 117, 4449–4459.

Sciacovelli, M., Gonçalves, E., Johnson, T.I., Zecchini, V.R., da Costa, A.S.H., Gaude, E., Drubbel, A.V., Theobald, S.J., Abbo, S.R., Tran, M.G.B., et al. (2016). Erratum: Corrigendum: Fumarate is an epigenetic modifier that elicits epithelial-to-mesenchymal transition. Nature 540, 150–150.

Sharp, A.J., Stathaki, E., Migliavacca, E., Brahmachary, M., Montgomery, S.B., Dupre, Y., and Antonarakis, S.E. (2011). DNA methylation profiles of human active and inactive X chromosomes. Genome Res. 21, 1592–1600.

Shyh-Chang, N., Locasale, J.W., Lyssiotis, C.A., Zheng, Y., Teo, R.Y., Ratanasirintrawoot, S., Zhang, J., Onder, T., Unternaehrer, J.J., Zhu, H., et al. (2013). Influence of Threonine Metabolism on S-Adenosylmethionine and Histone Methylation. Science (80-.). 339, 222–226.

Simopoulou, M., Sfakianoudis, K., Rapani, A., Giannelou, P., Anifandis, G., Bolaris, S., Pantou, A., Lambropoulou, M., Pappas, A., Deligeoroglou, E., et al. (2018). Considerations regarding embryo culture conditions: From media to epigenetics. In Vivo (Brooklyn). 32, 451–460.

Sims, R.J., Millhouse, S., Chen, C.-F., Lewis, B.A., Erdjument-Bromage, H., Tempst, P., Manley, J.L., and Reinberg, D. (2007). Recognition of Trimethylated Histone H3 Lysine 4 Facilitates the Recruitment of Transcription Postinitiation Factors and Pre-mRNA Splicing. Mol. Cell 28, 665–676.

Singh, R., and Sinclair, K.D. (2007). Metabolomics: Approaches to assessing oocyte and embryo quality. Theriogenology 68, S56–S62.

Spyrou, J., Gardner, D.K., and Harvey, A.J. (2019). Metabolism Is a Key Regulator of Induced Pluripotent Stem Cell Reprogramming. Stem Cells Int. 2019, 1–10.

Stover, P.J., and Caudill, M.A. (2008). Genetic and Epigenetic Contributions to Human Nutrition and Health: Managing Genome–Diet Interactions. J. Am. Diet. Assoc. 108, 1480–1487.

Stryer, L., Berg, J.M., Tymoczko, J.L., and Gatto Jr., G.J. (2019). Biochemistry (W.H. Freeman & Company).

Sturmey, R., Reis, A., Leese, H., and McEvoy, T. (2009). Role of Fatty Acids in Energy Provision During Oocyte Maturation and Early Embryo Development. Reprod. Domest. Anim. 44, 50–58.

TeSlaa, T., Chaikovsky, A.C., Lipchina, I., Escobar, S.L., Hochedlinger, K., Huang, J., Graeber, T.G., Braas, D., and Teitell, M.A. (2016). α-Ketoglutarate Accelerates the Initial Differentiation of Primed Human Pluripotent Stem Cells. Cell Metab. 24, 485–493.

Thompson, J.G., Partridge, R.J., Houghton, F.D., Cox, C.I., and Leese, H.J. (1996). Oxygen uptake and carbohydrate metabolism by in vitro derived bovine embryos. Reproduction 106, 299–306.

Torres-Osorio, V., Urrego, R., Echeverri Zuluaga, J.J., and López-Herrera, A. (2019). Estrés oxidativo y el uso de antioxidantes en la producción in vitro de embriones mamíferos. Revisión. Rev. Mex. Ciencias Pecu. 10, 433–459.

Torres-Padilla, M.-E., Parfitt, D.-E., Kouzarides, T., and Zernicka-Goetz, M. (2007). Histone arginine methylation regulates pluripotency in the early mouse embryo. Nature 445, 214–218.

Turner, B.M. (2008). Open Chromatin and Hypertranscription in Embryonic Stem Cells. Cell Stem Cell 2, 408–410.

Velazquez, M.A. (2015). Impact of maternal malnutrition during the periconceptional period on mammalian preimplantation embryo development. Domest. Anim. Endocrinol. 51, 27–45.

Ventura-Juncá, P., Irarrázaval, I., Rolle, A.J., Gutiérrez, J.I., Moreno, R.D., and Santos, M.J. (2015). In vitro fertilization (IVF) in mammals: epigenetic and developmental alterations. Scientific and bioethical implications for IVF in humans. Biol. Res. 48, 68.

Van Winkle, L.J., and Ryznar, R. (2019). One-Carbon Metabolism Regulates Embryonic Stem Cell Fate Through Epigenetic DNA and Histone Modifications: Implications for Transgenerational Metabolic Disorders in Adults. Front. Cell Dev. Biol. 7.

Wongtawan, T. (2010). Epigenetic and chromatin reprogramming in mouse development and embryonic stem cells. The University of Edinburgh.

Wright, K., Brown, L., Brown, G., Casson, P., and Brown, S. (2011). Microarray assessment of methylation in individual mouse blastocyst stage embryos shows that in vitro culture may have widespread genomic effects. Hum. Reprod. 26, 2576–2585.

Wu, D., Lim, E., Vaillant, F., Asselin-Labat, M.L., Visvader, J.E., and Smyth, G.K. (2010). ROAST: Rotation gene set tests for complex microarray experiments. Bioinformatics 26, 2176–2182.

Wu, X., Li, Y., Xue, L., Wang, L., Yue, Y., Li, K., Bou, S., Li, G.-P., and Yu, H. (2011). Multiple histone site epigenetic modifications in nuclear transfer and in vitro fertilized bovine embryos. Zygote 19, 31–45.

Wysocka, J., Swigut, T., Xiao, H., Milne, T.A., Kwon, S.Y., Landry, J., Kauer, M., Tackett, A.J., Chait, B.T., Badenhorst, P., et al. (2006). A PHD finger of NURF couples histone H3 lysine 4 trimethylation with chromatin remodelling. Nature 442, 86–90.

Xu, W., Yang, H., Liu, Y., Yang, Y., Wang, P., Kim, S.-H., Ito, S., Yang, C., Wang, P., Xiao, M.-T., et al. (2011). Oncometabolite 2-Hydroxyglutarate Is a Competitive Inhibitor of α-Ketoglutarate-Dependent Dioxygenases. Cancer Cell 19, 17–30.

Yang, M., and Vousden, K.H. (2016). Serine and one-carbon metabolism in cancer. Nat. Rev. Cancer 16, 650–662.

Zhang, A., Xu, B., Sun, Y., Lu, X., Gu, R., Wu, L., Feng, Y., and Xu, C. (2012). Dynamic changes of histone H3 trimethylated at positions K4 and K27 in human oocytes and preimplantation embryos. Fertil. Steril. 98, 1009–1016.

Zhang, Z., He, C., Zhang, L., Zhu, T., Lv, D., Li, G., Song, Y., Wang, J., Wu, H., Ji, P., et al. (2019). Alpha-ketoglutarate affects murine embryo development through metabolic and epigenetic modulations. Reproduction 158, 125–135.

Zhou, G., Soufan, O., Ewald, J., Hancock, R.E.W., Basu, N., and Xia, J. (2019). NetworkAnalyst 3.0: A visual analytics platform for comprehensive gene expression profiling and meta-analysis. Nucleic Acids Res. 47, W234–W241.

