## Supplementary file S1 for "RNAseq analysis reveals dynamic metaboloepigenetic profiles of human, mouse and bovine pre-implantation embryos"

**Supplementary table 1.** Overview of up and down significant (adjusted p-value < 0.05) differentially expressed genes (DEG) from Limma analysis of *in vivo* (bovine and mouse) and *in vitro* (bovine, mouse and human) pre-implantation development stages (adjusted p-value < 0.05). Up: number of up-regulated genes; Down: number of down regulated genes.

| **Species** | **Collection** | **Stage** | **Up** | **Down** | **NS** | **DEG (%)** | **Average**  **DEG (%)** |
| --- | --- | --- | --- | --- | --- | --- | --- |
| **Human** | *in vitro* | *2C-MII* | 135 | 25 | 12972 | 1.22 | 5.6 |
|  |  | *4C-2C* | 6 | 110 | 13016 | 0.88 |  |
|  |  | *8C-4C* | 141 | 1427 | 11564 | 11.94 |  |
|  |  | *MO-8C* | 348 | 972 | 11812 | 10.05 |  |
|  |  | *BL-MO* | 419 | 91 | 12622 | 3.88 |  |
| **Bovine** | *in vivo* | *2C-MII* | 88 | 3 | 13041 | 0.69 | 4.45 |
|  |  | *4C-2C* | 27 | 7 | 13098 | 0.26 |  |
|  |  | *8C-4C* | 1151 | 1125 | 10856 | 17.33 |  |
|  |  | *16C-8C* | 30 | 26 | 13076 | 0.43 |  |
|  |  | *BL-16C* | 271 | 196 | 12665 | 3.56 |  |
|  | *in vitro* | *2C-MII* | 2284 | 2336 | 8512 | 35.18 | 20.97 |
|  |  | *4C-2C* | 2373 | 2178 | 8581 | 34.66 |  |
|  |  | *8C-4C* | 400 | 208 | 12524 | 4.63 |  |
|  |  | *16C-8C* | 269 | 320 | 12543 | 4.49 |  |
|  |  | *BL-16C* | 1664 | 1736 | 9732 | 25.89 |  |
| **Mouse** | *in vivo* | *2C-MII* | 2007 | 1477 | 9648 | 26.53 | 21.37 |
|  |  | *4C-2C* | 2035 | 1875 | 9222 | 29.77 |  |
|  |  | *8C-4C* | 480 | 135 | 12517 | 4.68 |  |
|  |  | *MO-8C* | 494 | 122 | 12516 | 4.69 |  |
|  |  | *BL-MO* | 2736 | 2668 | 7728 | 41.15 |  |
|  | *in vitro* | *4C-2C* | 2880 | 3143 | 7109 | 45.87 | 21.95 |
|  |  | *8C-4C* | 561 | 387 | 12184 | 7.22 |  |
|  |  | *MO-8C* | 217 | 142 | 12773 | 2.73 |  |
|  |  | *BL-MO* | 2391 | 1810 | 8931 | 31.99 |  |

**Supplementary table 2.** Metabolic and epigenetic Reactome terms used for ROAST analysis.

|  |  | *Gene list* |
| --- | --- | --- |
| Carbohydrate  metabolism | Glycogen metabolism | *ALDOA; ALDOB; ALDOC; GPI; TPI1; GAPDHS; GAPDH; PGAM1; PGAM2; PGK2; PGK1; ENO2; ENO3; ENO1; PFKL; PFKM; PFKP; ADPGK; GCK; GCKR; NUP98; SEH1L; NUP107; NUP43; NUP160; NUP37; NUP85; NUP133; SEC13; TPR; NUP153; NUP88; NUP188; NUP214; NDC1; NUP210; POM121; POM121C; NUP35; NUP93; NUP155; NUP205; SEH1L; RAE1; NUP98; NUP98; NUP62; NUP58; NUP58; NUP54; NUP50; AAAS; RANBP2; NUP42; GNPDA1; GNPDA2; HK1; HK2; HK3; PKM; PKLR; PKLR; PKM; BPGM; PGM2L1; PGP; PPP2R1A; PPP2R1B; PPP2CA; PPP2CB; PPP2R5D; PFKFB1; PFKFB3; PFKFB4; PFKFB2; PRKACA; PRKACB; PRKACG* |
|  | Gluconeogenesis | *SORD; AKR1B1* |
|  | Glycolysis | *GLYCTK; ALDH1A1; ALDOB; KHK; TKFC* |
|  | Fructose  biosynthesis | *B4GALT1; LALBA; SLC2A1* |
|  | Fructose  catabolism | *GLYCTK; ALDH1A1; ALDOB; KHK; TKFC* |
|  | Lactose  synthesis | *B4GALT1; LALBA; SLC2A1* |
|  | Galactose  catabolism | *GALE; GALK1; GALT; PGM2; PGM2L1* |
|  | PPP | *PGLS; PGD; PGM2; TKT; RPEL1; RPE; TALDO1; G6PD; RPIA; RBKS; SHPK; PRPS2; PRPS1L1; PRPS1; DERA* |
|  | Glycosaminoglycan  metabolism | *HAS2; HAS1; HAS3; CEMIP; ABCC5; HEXB; HEXA; GUSB; HYAL2; HMMR; STAB2; CD44; LYVE1; SLC9A1; CHP1; HYAL3; HYAL1; CHST14; DCN; BCAN; CSPG4; NCAN; BGN; CSPG5; VCAN; UST; DSE; DSEL; CHSY1; CHPF; CHSY3; CSGALNACT1; CSGALNACT2; CHST7; CHST3; CHST11; CHST9; CHST13; CHST12; CHPF2; CHST15; IDS; IDUA; ARSB; B4GALT7; GPC2; AGRN; GPC6; GPC3; SDC1; GPC4; SDC2; GPC1; SDC3; HSPG2; SDC4; GPC5; B3GAT3; B3GALT6; XYLT2; XYLT1; B3GAT1; B3GAT2; SLC35B2; SLC35B3; SLC26A2; SLC26A1; PAPSS1; PAPSS2; HS3ST5; HS3ST1; HS2ST1; SLC35D2; GLCE; NDST3; NDST4; NDST1; NDST2; EXT2; EXT1; HS3ST4; HS3ST3A1; HS3ST3B1; HS3ST2; HS3ST6; HS6ST3; HS6ST1; HS6ST2; GLB1L; GLB1; HPSE; SGSH; NAGLU; HGSNAT; HPSE2; ACAN; FMOD; LUM; OMD; OGN; PRELP; KERA; GALNS; GNS; CHST1; CHST2; CHST5; CHST6; B3GNT2; B3GNT7; B4GAT1; B3GNT4; B3GNT3; ST3GAL2; ST3GAL1; ST3GAL4; ST3GAL6; ST3GAL3; B4GALT1; B4GALT6; B4GALT3; B4GALT4; B4GALT5; B4GALT2* |
| Inositol  Phosphate  metabolism | Inositol  Phosphate  metabolism | *NUDT11; NUDT3; NUDT10; NUDT4; IP6K1; IP6K3; ITPK1; PPIP5K1; PPIP5K2; IPPK; MINPP1; MIOX; INPP5B; INPP5A; IMPA2; IMPA1; INPP4A; INPP4B; MTMR9; MTMR7; INPP1; ISYNA1; SYNJ1; INPP5J; OCRL; PLD4; PLCD4; PLCB4; PLCB3; PLCB2; PLCB1; PLCZ1; PLCD3; PLCH2; PLCE1; PLCG2; PLCH1; PLCG1; PLCD1; PTEN; ITPKA; CALM1; ITPKC; ITPKB; INPP5D; INPPL1; IPMK; IP6K2* |
| Lipids  metabolism | Fatty  AcylCoA  biosynthesis | *FASN; PPT1; OLAH; ACSF3; SLC27A3; ACSBG2; ACSBG1; ACSL3; ACSL4; ELOVL6; ELOVL3; ELOVL7; ELOVL1; ELOVL4; TECRL; TECR; ACSL5; ACSL6; ACSL1; ELOVL2; ELOVL5; HSD17B3; HSD17B12; HACD4; HACD3; HACD1; HACD2; HTD2; SLC25A1; ACACA; PPT2; ACLY; MORC2; CBR4; HSD17B8; SCD; SCD5* |
|  | Arachdonic Acid metabolism | *PON2; PON3; PON1; ALOX5AP; ALOX5; LTC4S; GPX2; GPX4; GPX1; CYP1A2; CYP1B1; CYP1A1; CYP2U1; CYP4A11; CYP4F2; CYP2C8; CYP2C9; CYP2C19; FAAH2; CYP4A22; CYP4F22; CYP4B1; CYP4F11; CYP4F8; CYP4F3; ALOX15; ABCC1; DPEP3; DPEP2; DPEP1; GGT5; GGT1; LTA4H; MAPKAPK2; PTGR1; ALOX12; PLA2G4A; PTGR2; PTGS2; PTGDS; AKR1C3; TBXAS1; PRXL2B; HPGD; PTGES; PTGS1; CBR1; CYP8B1; PTGIS; PTGES3; PTGES2; HPGDS; CYP2J2; EPHX2; ALOX15B; FAAH; ALOXE3; ALOX12B; AWAT1* |
|  | O3 O6  metabolism | *SCP2; HSD17B4; ACOT8; ELOVL2; ELOVL5; FADS2; FADS1; ELOVL1; ELOVL3; ABCD1; ACSL1; ACAA1; ACOX1* |
|  | Carnitine  metabolism | *SLC22A5; PRKAA2; PRKAG2; PRKAB2; ACACB; CPT1A; CPT1B; RXRA; PPARD; CPT2; MID1IP1; THRSP; ACACA; SLC25A20* |
|  | B oxidation | *ACBD7; DBI; ACSF2; ACAD11; MCAT; NDUFAB1; ACOT9; ACOT2; THEM5; THEM4; ACADM; MECR; ECHS1; HADHA; HADHB; HADH; ACADL; ACADS; ACADVL; ACSM6; ACSM3; ECI1; DECR1; MMUT; MMAA; PCCA; PCCB; MCEE; ACAD10; ACOT7L; ACOT7; ACOT11; ACOT12; ACOT13; ACOT1; PCTP; ACBD6; ACAA2* |
|  | Peroxisomal lipid  metabolism | *ACBD4; ACBD5; ACOX1; ECI2; EHHADH; DECR2; HSD17B4; MLYCD; ABCD1; ACOT4; ACOT8; ACOT6; ACAA1; HAO2; SLC25A17; PHYH; SLC27A2; PECR; ALDH3A2; HACL1; ACOX2; ACOXL; SCP2; ACOX3; CRAT; AMACR; CROT; NUDT7; NUDT19* |
|  | Triglyceris biosynthesis | *DGAT2; AGMO; MOGAT1; MOGAT2; MOGAT3; LPIN3; LPIN2; LPIN1; DGAT1; GK2; GK3P; GK; GPAT2; GPAM* |
|  | Triglyceride catabolism | *LIPE; PNPLA4; PLIN3; FABP4; MGLL; PLIN1; ABHD5; GPD2; PPP1CC; PPP1CA; PPP1CB; PNPLA5; PRKACA; PRKACB; PRKACG; CAV1; FABP9; FABP7; FABP2; FABP1; FABP6; FABP3; FABP12; FABP5* |
|  | Phospholipid  metabolism | *MBOAT2; LPCAT3; LPCAT4; MBOAT1; PLA2G2A; PNPLA8; ABHD4; PLA2G4E; PLA2G4A; PLAAT3; PLBD1; PLA2G4F; PLA2G4B; PLA2G4D; PLA2G6; PLA2G2D; PLA2G10; PLA2G3; PLA2G1B; PLA2G2F; PLA2G12A; PLA2G2E; PLA2G5; PLA2R1; PLA2G4C; PLAAT5; PLAAT1; PLAAT4; PLAAT2; STARD7; PHOSPHO1; BCHE; ACHE; MFSD2A; ABHD3; LPCAT1; STARD10; PCTP; CEPT1; CHKA; CHKB; CHAT; CHPT1; CSNK2A1; CSNK2A2; CSNK2B; PEMT; LPIN2; LPIN3; LPIN1; SLC44A2; SLC44A4; SLC44A3; SLC44A5; SLC44A1; PCYT1B; PCYT1A; GPCPD1; OSBPL5; OSBPL8; OSBPL10; PLA1A; AGPAT5; ALPI; GNPAT; AGPAT4; GPAT4; LCLAT1; AGPAT1; AGPAT2; GPAT3; AGPAT3; GPD1L; GPD1; GPD2; GPAT2; GPAM; PLD6; MIGA1; MIGA2; PLD2; PLD1; ACP6; LIPI; LIPH; DDHD1; DDHD2; PITPNB; PTDSS1; PTDSS2; CDIPT; CDS1; PITPNM3; PITPNM1; PITPNM2; LPCAT2; TMEM86B; PLB1; CRLS1; PNPLA3; PNPLA2; DGAT2L6; DGAT2L7P; AWAT2; MGLL; DGAT2; DGAT1; HADHA; HADHB; TAZ; PLD3; PLD4; CDS2; PTPMT1; PGS1; ETNK2; ETNK1; PCYT2; SELENOI; ETNPPL; PISD; LPGAT1; MBOAT7; AGK; PLA2G15; CPNE1; CPNE3; CPNE6; CPNE7; MTMR10; MTMR2; MTMR12; MTMR4; MTM1; PIKFYVE; FIG4; VAC14; INPP5F; PIK3C2A; PIK3C3; PIK3R4; INPP4A; INPP4B; PI4K2A; PI4K2B; MTMR9; MTMR7; PI4KA; ARF3; ARF1; PI4KB; PIK3C2G; TPTE; TPTE2; INPP5E; OCRL; SACM1L; PIP4K2B; PIP4K2C; PIP4K2A; PIP4P1; SBF2; PIP5K1B; PIP5K1A; PTPN13; PLEKHA1; PLEKHA2; MTMR6; INPP5K; INPP5J; INPP5D; INPPL1; MTMR14; SYNJ1; MTMR3; SYNJ2; MTMR1; PTEN; PIP5K1C; PLEKHA3; PLEKHA8; PLEKHA5; PLEKHA6; PLEKHA4; RUFY1; PIK3CA; PIK3R2; PIK3R3; PIK3CB; PIK3R5; PIK3CG; PIK3CD; PIK3R6; PIK3C2B; PIK3R1; MTMR8; BMX; RAB4A; RAB14; RAB5A; GDPD5; GDPD1; PNPLA6; ENPP6; GDE1; GDPD3; PNPLA7; SBF1; TNFAIP8L1; TNFAIP8; TNFAIP8L2; TNFAIP8L3* |
|  | Sphingolipid  metabolism | *SGPP2; SGPP1; VAPB; VAPA; PPM1L; CERT1; ALDH3B2; SPHK1; OSBP; ALDH3A2; SPHK2; SAMD8; ACER2; DEGS1; DEGS2; ACER3; SGMS1; CERS1; CERS6; CERS3; CERS4; CERS2; CERS5; PLPP2; PLPP3; PLPP1; KDSR; SGMS2; SPTLC3; SPTSSB; SPTSSA; SPTLC1; SPTLC2; ORMDL2; ORMDL3; ORMDL1; FA2H; CSNK1G2; SGPL1; ALDH3B1; PRKD1; PRKD3; PRKD2; SPNS2; ACER1; ENPP7; SMPD1; GALC; ESYT2; ESYT3; ESYT1; GLTP; GLA; GLB1L; GLB1; STS; ASAH2; UGT8; CERK; CPTP; SMPD3; SMPD2; ARSA; B4GALNT1; SUMF1; SUMF2; HEXB; HEXA; ASAH1; NEU3; GBA; PSAP; GM2A; SMPD4; B3GALNT1; NEU1; CTSA; NEU4; NEU2; ARSG; ARSJ; ARSB; ARSD; ARSL; ARSF; ARSH; ARSK; ARSI; GBA3; UGCG; GBA2* |
|  | Steroids  metabolism | *GC; CUBN; VDR; CYP24A1; LGMN; CYP27B1; CYP2R1; PIAS4; UBE2I; SUMO2; LRP2; SERPINA6; HSD11B2; HSD11B1; HSD3B2; HSD3B1; CYP17A1; CYP21A2; CYP11B2; CYP11B1; POMC; HSD17B12; HSD17B3; CGA; LHB; SRD5A2; SRD5A3; SRD5A1; AKR1B15; CYP19A1; HSD17B1; HSD17B14; HSD17B11; HSD17B2; TSPO; TSPOAP1; STAR; FDX1; FDX2; CYP11A1; FDXR; STARD4; STARD6; STARD3; STARD3NL; AKR1B1; SLCO1A2; ALB; ABCB11; SLCO1B1; NR1H4; SLC27A5; BAAT; SLC10A2; FABP6; RXRA; NCOA1; NCOA2; ABCC3; SLC10A1; SLCO1B3; STARD5; CYP7A1; CYP7B1; OSBPL2; OSBPL6; OSBPL3; OSBPL9; OSBPL7; OSBPL1A; OSBP; SCP2; HSD17B4; CYP27A1; SLC27A2; AMACR; ACOT8; AKR1C3; AKR1C2; AKR1C1; AKR1C4; AKR1D1; HSD3B7; PTGIS; CYP8B1; ACOX2; CYP46A1; CYP39A1; CH25H; PMVK; HMGCR; HMGCR; CYP51A1; MVD; ARV1; DHCR7; SC5D; EBP; DHCR24; LBR; MSMO1; FDFT1; NSDHL; HMGCS1; FDPS; GGPS1; MVK; IDI2; IDI1; ACAT2; HSD17B7; TM7SF2; LSS; SQLE; PLPP6; SREBF1; SREBF2; SREBF1; KPNB1; MBTPS2; SCAP; SAR1B; SEC24C; SEC24D; SEC24A; SEC24B; SEC23A; INSIG2; INSIG1; MBTPS1; RAN; NFYA; NFYB; NFYC; SP1; ACACA; FASN; TBL1XR1; HELZ2; CHD9; TGS1; CREBBP; NCOA6; SMARCD3; CARM1; MED1; PPARA; TBL1X; SCD; ACACB; MTF1; GPAM; ELOVL6; MVK; CYP51A1; SQLE; ACACA; FASN; SC5DL; FDFT1; HMGCS1; TM7SF2; SCD; HMGCR; IDI1; GGPS1; ACACB; LSS; DHCR7; GPAM; FDPS; PMVK; MVD; ELOVL6* |
|  | Ketone  Body metabolism | *OXCT1; OXCT2; BDH1; ACAT1; HMGCLL1; HMGCL; AACS; ACSS3; BDH2; HMGCS2* |
| Nitric oxide  metabolism | Nitric oxide metabolism | *NOS3; NOSIP; CAV1; NOSTRIN; WASL; DNM2; HSP90AA1; CALM1; AKT1; CYGB; DDAH1; DDAH2; ZDHHC21; SPR; LYPLA1* |
| Pyruvate  metabolism | Pyruvate metabolism | *DLD; GLO1; HAGH; MPC1; MPC2; PDHB; PDHA2; PDHA1; DLAT; PDHX; LDHAL6B; BSG; SLC16A3; SLC16A8; SLC16A1; LDHAL6A; LDHB; LDHA; LDHC; ME1; GSTZ1; PDK2; PDK4; PDK3; PDK1; RXRA; PPARD; PDP1; PDPR; PDP2; VDAC1* |
| TCA | TCA | *SDHA; SDHB; SDHD; SDHC; MDH2; ME2; FH; OGDH; DLST; DLD; IDH2; SUCLA2; SUCLG1; ME3; IDH3B; IDH3A; IDH3G; ACO2; FAHD1; NNT; CS; SUCLG2* |
| ETC | ETC | *COQ10A; COQ10B; NDUFS5; NDUFS6; NDUFV1; NDUFV3; NDUFA12; NDUFS4; NDUFS1; NDUFV2; MT-ND4; MT-ND5; MT-ND1; NDUFA9; NDUFS3; NDUFS8; NDUFS7; NDUFS2; NDUFB8; NDUFA13; NDUFB10; NDUFB7; NDUFA10; NDUFC1; NDUFB2; NDUFB5; NDUFB9; NDUFA11; NDUFA2; NDUFA7; NDUFA8; NDUFA1; NDUFA3; NDUFC2; NDUFB1; NDUFAB1; NDUFA5; NDUFB3; NDUFA6; NDUFB4; NDUFB11; NDUFB6; MT-ND2; MT-ND6; MT-ND3; ETFDH; ETFB; ETFA; NDUFAF2; NDUFAF7; NDUFAF3; NDUFAF4; TIMMDC1; TMEM126B; ECSIT; NDUFAF1; ACAD9; NDUFAF5; NDUFAF6; NUBPL; UQCRFS1; MT-CYB; UQCR10; UQCRQ; UQCRC2; CYC1; UQCRH; UQCRB; UQCRC1; UQCR11; CYCS; COX6C; COX6B1; NDUFA4; MT-CO2; COX6A1; COX7C; COX5B; COX4I1; COX7A2L; COX5A; COX7B; MT-CO1; MT-CO3; COX8A; COX18; COX14; COX16; COX20; COX11; COX19; SCO2; TACO1; SCO1; SURF1; LRPPRC; SDHA; SDHB; SDHD; SDHC; TRAP1; ATP5PO; ATP5MC3; MT-ATP6; ATP5ME; ATP5MF; MT-ATP8; ATP5MC1; ATP5MG; ATP5PF; ATP5PD; ATP5PB; ATP5MC2; ATP5F1B; ATP5F1C; ATP5F1D; ATP5F1E; ATP5F1A; DMAC2L; SLC25A14; SLC25A27; UCP1; UCP2; UCP3; PM20D1* |
| Nucleotides  metabolism | Nucleotides metabolism | *RRM1; RRM2; GLRX; DCTD; RRM2B; TXN; GUK1; NME1; NME2; CMPK1; AK1; NME4; AK8; AK7; AK9; AK5; NME2P1; NME3; DTYMK; DCTPP1; CTPS2; TXNRD1; CTPS1; AK6; AK4; NUDT13; TYMS; DUT; GSR; AK2; ITPA; PNP; NT5C1B; XDH; NT5C2; NT5C1A; GDA; NUDT15; NUDT18; NUDT1; NUDT1; NUDT1; NUDT1; NUDT9; NUDT16; ADPRM; NUDT5; DNPH1; GPX1; NT5E; NT5C; NT5C3A; TYMP; UPP1; UPP2; NT5M; DPYD; DPYS; UPB1; AGXT2; ENTPD1; ENTPD3; ENTPD7; ENTPD6; ENTPD4; ENTPD8; ENTPD2; ENTPD5; SAMHD1; DHODH; CAD; UMPS; GMPS; IMPDH2; IMPDH1; ATIC; ADSS2; ADSS1; ADSL; PAICS; GART; PPAT; PFAS; LHPP; ADA; ADAL; APRT; DCK; HPRT1; ADK; GMPR2; GMPR; DGUOK; AMPD1; AMPD3; AMPD2; PUDP; TK1; CDA; TK2; UCK1; UCK2; UCKL1* |
| Vitamins and  cofactors  metabolism | Ascorbate metabolism | *SLC2A3; SLC2A1; SLC23A1; SLC23A2; CYB5R3; CYB5A; GSTO1; GSTO2; LOC785216* |
|  | Thiamin metabolism | *THTPA; SLC19A2; SLC19A3; SLC25A19; TPK1* |
|  | Riboflavin metabolism | *ENPP1; FLAD1; ACP5; RFK; SLC52A2; SLC52A1; SLC52A3* |
|  | Panthotenate metabolism | *AASDHPPT; FASN; ENPP2; ENPP3; ENPP1; VNN2; VNN1; SLC5A6; PDZD11; PANK3; PANK4; PANK1; COASY; PPCS; SLC25A16; PANK2; PPCDC* |
|  | Cobalamin metabolism | *MMACHC; MTR; MTRR; PRSS1; PRSS3; CTRB2; CTRB1; TCN1; ABCC1; MMUT; MMAA; TCN2; MMAB; AMN; CUBN; CBLIF; ABCD4; CD320; CTRC; MMADHC; LMBRD1* |
|  | Biotin metabolism | *SLC5A6; PDZD11; HLCS; PC; BTD; ACACB; ACACA; MCCC2; MCCC1; PCCA; PCCB* |
|  | Nicotinate metabolism | *NMRK2; NMRK1; NT5E; NMNAT1; NADK; BST1; NMNAT2; NAXD; NUDT12; PARP14; PARP6; PARP8; PARP4; PARP10; PARP9; PARP16; NAMPT; NAPRT; CYP8B1; PTGIS; NAXE; RNLS; SLC5A8; SLC22A13; NNMT; PTGS2; NADSYN1; CD38; QPRT; NMNAT3; NADK2* |
|  | Folate metabolism | *SHMT2; FOLR2; SLC25A32; FPGS; FPGS; MTHFD1; MTHFR; SHMT1; SLC46A1; MTHFS; MTHFD1L; MTHFD2L; MTHFD2; ALDH1L1; DHFR2; DHFR; SLC19A1; ALDH1L2* |
|  | Vitamin K metabolism | *VKORC1L1; UBIAD1; VKORC1* |
|  | Retinoid metabolism | *AKR1B10; AKR1C3; AKR1C4; AKR1C1; RBP2; LRAT; RBP1; APOE; GPC2; AGRN; GPC4; SDC4; GPC3; SDC3; GPC1; GPC6; SDC2; HSPG2; GPC5; SDC1; APOC2; APOA2; APOA4; APOC3; APOA1; APOB; PNLIP; CLPS; PLB1; BCO2; APOM; BCO1; LPL; GPIHBP1; LDLR; LRP12; LRP10; LRP1; LRP2; LRP8; RBP4; RETSAT; RDH11; TTR* |
|  | Vitamin e metabolism | *TTPA* |
|  | CoFactors metabolism | *SPR; PRKG2; PTS; HSP90AA1; AKT1; CALM1; NOS3; GCH1; GCHFR; DHFR; IDH1; COQ2; COQ3; COQ9; COQ7; COQ6; COQ5; PDSS1; PDSS2* |
| Amino  Acids  metabolism | Aspartate asparigine metabolism | *GOT1; FOLH1; NAALAD2; FOLH1B; SLC25A13; SLC25A12; GOT2; ASPA; ASNS; GADL1; ASPG; NAT8L* |
|  | Glutamate glutamine metabolism | *PYCR2; RIMKLA; RIMKLB; PYCR1; GLUD2; GLUD1; OAT; ALDH18A1; ALDH18A1; GLUL; GLS; GLS2; PYCR3; GOT2* |
|  | Alanine metabolism | *GPT2; GPT* |
|  | Branched chain aa catabolism | *AUH; HIBCH; BCAT1; HIBADH; BCAT2; BCKDK; DBT; BCKDHB; BCKDHA; DLD; ACAD8; ALDH6A1; SLC25A44; ACADSB; MCCC2; MCCC1; HSD17B10; PPM1K; ECHS1; IVD; ACAT1* |
|  | Histidine catabolism | *HAL; CARNS1; CARNMT1; UROC1; FTCD; AMDHD1; HDC; HNMT* |
|  | Lysine catabolism | *HYKK; SLC25A21; GCDH; AADAT; PHYKPL; PIPOX; CRYM; AASS; ALDH7A1; OGDH; DLST; DLD* |
|  | Phenylalanine metabolism | *KYAT1; ASRGL1; PCBD1; QDPR; IL4I1; PAH* |
|  | Tyrosine catabolism | *TAT; GSTZ1; HPD; FAH; HGD* |
|  | Proline catabolism | *PRODH2; ALDH4A1; PRODH* |
|  | Serine biosynthesis | *PSPH; SRR; PHGDH; PSAT1; SERINC1; SERINC2; SERINC4; SERINC3; SERINC5* |
|  | Threonine catabolism | *RIDA; SDS; SDSL; GCAT; TDH* |
|  | Tryptophan catabolism | *IDO1; KMO; AADAT; KYAT3; IDO2; SLC7A5; SLC3A2; KYNU; KYAT1; HAAO; AFMID; TDO2; ACMSD; SLC36A4* |
|  | Methionine salvage pathway | *MRI1; APIP; MTAP; ENOPH1; GOT1; ADI1* |
|  | Sulfur aa metabolism | *BHMT2; AHCY; CBS; CBSL; CTH; MTR; MTRR; MAT1A; BHMT; TST; CSAD; CDO1; MPST; SQOR; ETHE1; SUOX; SLC25A10; TSTD1; ADO; GADL1; TXN2; GOT2; MRI1; APIP; MTAP; ENOPH1; GOT1; ADI1* |
|  | Selenoamino acid metabolism | *TXNRD1; PAPSS1; PAPSS2; GSR; DARS1; EPRS1; LARS1; RARS1; EEF1E1; KARS1; QARS1; IARS1; MARS1; AIMP1; AIMP2; AHCY; MAT1A; HNMT; GNMT; NNMT; SCLY; CBS; CTH; SECISBP2; RPL29; RPL28; RPL27A; RPL27; RPL3L; RPL3; RPL24; RPL10L; RPL10; RPL23A; RPL23; RPL21; RPL19; RPL22L1; RPL22; RPL18A; RPL18; RPL17; RPL15; RPL14; RPL13A; RPL26L1; RPL26; RPL13; RPL12; RPL11; RPL10A; RPL36AL; RPL36A; RPLP2; RPLP1; RPLP0; RPL39L; RPL39; RPL9; RPL8; RPL7A; RPL7; RPL6; RPL5; RPL41; UBA52; RPL4; RPL38; RPL37A; RPL37; RPL36; RPL35A; RPL35; RPL34; RPL32; RPL31; RPL30; RPS7; RPS6; RPS5; RPS3A; FAU; RPS3; RPS29; RPS28; RPS27A; RPS4Y2; RPS4X; RPS4Y1; RPS26; RPS25; RPS24; RPS27L; RPS27; RPS23; RPS21; RPS20; RPS2; RPS15A; RPS19; RPS18; RPS17; RPS16; RPS15; RPS14; RPS13; RPS12; RPS11; RPSA; RPS10; RPS9; RPS8; EEFSEC; SARS1; SEPHS2; PSTK; SEPSECS; INMT; rRNA; rRNA; rRNA; rRNA* |
|  | Glyoxylate metabolism | *HOGA1; HAO1; PRODH2; GRHPR; ALDH4A1; GNMT; AGXT2; GOT2; LIAS; GCSH; AGXT; LIPT2; NDUFAB1; DLD; GLDC; AMT; DAO; DDO; LIPT1; BCKDHB; BCKDHA; DBT; DLST; OGDH; DHTKD1; PDHB; PDHA2; PDHA1; PDHX; DLAT; PXMP2* |
|  | Urea cycle | *NMRAL1; ASS1; OTC; ARG2; ARG1; CPS1; SLC25A2; SLC25A15; ASL; NAGS* |
|  | Carnitine synthesis | *SHMT1; TMLHE; BBOX1; ALDH9A1* |
|  | Creatine metabolism | *CKMT2; CKMT1A; GAMT; GATM; CKM; CKB; SLC6A12; SLC6A11; SLC6A7; SLC6A8* |
|  | Choline catabolism | *DMGDH; SLC44A1; ALDH7A1; BHMT; CHDH; SARDH* |
|  | Polyamines metabolism | *AZIN2; AGMAT; AMD1; ODC1; SAT1; PAOX; SMOX; PSMD3; PSMA6; PSMD2; PSMA5; PSMD13; PSMA4; PSMD12; PSMA3; PSMD11; PSMA2; SEM1; PSMD10; PSMA1; PSMD1; PSMD14; PSMC6; PSMC5; PSMC4; PSMC3; PSMC2; PSMC1; PSMB9; PSMB8; PSMB11; PSME4; PSMB7; PSMB6; PSMA8; PSMF1; PSME3; PSMB5; PSME2; PSMB4; PSME1; PSMD9; PSMB3; PSMD8; PSMB2; PSMD7; PSMD6; PSMB10; PSMD5; PSMB1; PSMD4; PSMA7; OAZ2; OAZ1; OAZ3; NQO1; AZIN1; SRM; SMS* |
|  | Melanin biosynthesis | *TYR; OCA2; SLC45A2; DCT; TYRP1* |
|  | Amine derived hormones metabolism | *DBH; DDC; TH; PNMT; TPO; IYD; CGA; TSHB; SLC5A5; DIO3; DIO2; DIO1; DUOX1; DUOX2; AANAT; TPH1; TPH2; ASMT* |
| Mitochondria iron sulfur  biogenesis | Mitochondria iron sulfur biogenesis | *FXN; ISCU; NFS1; LYRM4; SLC25A37; SLC25A28; FDXR; FDX1; FDX2; GLRX5; ISCA2; ISCA1; HSCB* |
| Reversible  Hydration  Of CO2 | Reversible hydration of CO2 | *CA5A; CA5B; CA3; CA1; CA13; CA2; CA7; CA9; CA12; CA4; CA14; CA6* |
| Protein  metabolism | tRNA Aminoacylation | *HARS1; DARS1; EPRS1; LARS1; RARS1; EEF1E1; KARS1; QARS1; IARS1; MARS1; AIMP1; AIMP2; TARS1; YARS1; CARS1; WARS1; FARSA; FARSB; VARS1; GARS1; PPA1; NARS1; AARS1; SARS1; DARS2; VARS2; PARS2; RARS2; NARS2; SARS2; TARS2; LARS2; FARS2; HARS2; YARS2; AARS2; IARS2; PPA2; MARS2; CARS2; WARS2; EARS2* |
|  | Eukaryotic Translation Initiation | *RPL29; RPL28; RPL27A; RPL27; RPL3L; RPL3; RPL24; RPL10L; RPL10; RPL23A; RPL23; RPL21; RPL19; RPL22L1; RPL22; RPL18A; RPL18; RPL17; RPL15; RPL14; RPL13A; RPL26L1; RPL26; RPL13; RPL12; RPL11; RPL10A; RPL36AL; RPL36A; RPLP2; RPLP1; RPLP0; RPL39L; RPL39; RPL9; RPL8; RPL7A; RPL7; RPL6; RPL5; RPL41; UBA52; RPL4; RPL38; RPL37A; RPL37; RPL36; RPL35A; RPL35; RPL34; RPL32; RPL31; RPL30; RPS7; RPS6; RPS5; RPS3A; FAU; RPS3; RPS29; RPS28; RPS27A; RPS4Y2; RPS4X; RPS4Y1; RPS26; RPS25; RPS24; RPS27L; RPS27; RPS23; RPS21; RPS20; RPS2; RPS15A; RPS19; RPS18; RPS17; RPS16; RPS15; RPS14; RPS13; RPS12; RPS11; RPSA; RPS10; RPS9; RPS8; EIF1AX; EIF3H; EIF3A; EIF3D; EIF3C; EIF3E; EIF3L; EIF3K; EIF3M; EIF3J; EIF3I; EIF3G; EIF3F; EIF3B; EIF5B; EIF2S2; EIF2S3; EIF2S1; EIF4A1; EIF4A2; EIF4E; EIF4G1; EIF4B; EIF4H; EIF5; PABPC1; EIF4EBP1; EIF2B1; EIF2B2; EIF2B4; EIF2B5; EIF2B3; 28S rRNA; 5.8s rRNA; 5s rRNA; 18s rRNA; M13699* |
|  | SRP-dependent co translational protein targeting to membrane | *SRPRA; SRPRB; SRP68; SRP54; SRP19; SRP14; SRP72; SRP9; RPL29; RPL28; RPL27A; RPL27; RPL3L; RPL3; RPL24; RPL10L; RPL10; RPL23A; RPL23; RPL21; RPL19; RPL22L1; RPL22; RPL18A; RPL18; RPL17; RPL15; RPL14; RPL13A; RPL26L1; RPL26; RPL13; RPL12; RPL11; RPL10A; RPL36AL; RPL36A; RPLP2; RPLP1; RPLP0; RPL39L; RPL39; RPL9; RPL8; RPL7A; RPL7; RPL6; RPL5; RPL41; UBA52; RPL4; RPL38; RPL37A; RPL37; RPL36; RPL35A; RPL35; RPL34; RPL32; RPL31; RPL30; RPS7; RPS6; RPS5; RPS3A; FAU; RPS3; RPS29; RPS28; RPS27A; RPS4Y2; RPS4X; RPS4Y1; RPS26; RPS25; RPS24; RPS27L; RPS27; RPS23; RPS21; RPS20; RPS2; RPS15A; RPS19; RPS18; RPS17; RPS16; RPS15; RPS14; RPS13; RPS12; RPS11; RPSA; RPS10; RPS9; RPS8; RPN1; DDOST; TRAM1; SSR2; SSR4; SSR3; SSR1; SEC61B; SEC61A1; SEC61A2; SEC61G; RPN2; SPCS2; SPCS3; SEC11A; SEC11C; SPCS1; 7SL RNA; 7SL RNA; 28S rRNA; 5.8s rRNA; 5s rRNA; 18s rRNA* |
|  | Eukaryotic Translation Elongation | *EEF1D; EEF1B2; EEF1G; EEF1A1; EEF2; RPL29; RPL28; RPL27A; RPL27; RPL3L; RPL3; RPL24; RPL10L; RPL10; RPL23A; RPL23; RPL21; RPL19; RPL22L1; RPL22; RPL18A; RPL18; RPL17; RPL15; RPL14; RPL13A; RPL26L1; RPL26; RPL13; RPL12; RPL11; RPL10A; RPL36AL; RPL36A; RPLP2; RPLP1; RPLP0; RPL39L; RPL39; RPL9; RPL8; RPL7A; RPL7; RPL6; RPL5; RPL41; UBA52; RPL4; RPL38; RPL37A; RPL37; RPL36; RPL35A; RPL35; RPL34; RPL32; RPL31; RPL30; RPS7; RPS6; RPS5; RPS3A; FAU; RPS3; RPS29; RPS28; RPS27A; RPS4Y2; RPS4X; RPS4Y1; RPS26; RPS25; RPS24; RPS27L; RPS27; RPS23; RPS21; RPS20; RPS2; RPS15A; RPS19; RPS18; RPS17; RPS16; RPS15; RPS14; RPS13; RPS12; RPS11; RPSA; RPS10; RPS9; RPS8; EEF1A1P5; EEF1A2; 28S rRNA; 5.8s rRNA; 5s rRNA; 18s rRNA* |
|  | Eukaryotic Translation Termination | *RPL29; RPL28; RPL27A; RPL27; RPL3L; RPL3; RPL24; RPL10L; RPL10; RPL23A; RPL23; RPL21; RPL19; RPL22L1; RPL22; RPL18A; RPL18; RPL17; RPL15; RPL14; RPL13A; RPL26L1; RPL26; RPL13; RPL12; RPL11; RPL10A; RPL36AL; RPL36A; RPLP2; RPLP1; RPLP0; RPL39L; RPL39; RPL9; RPL8; RPL7A; RPL7; RPL6; RPL5; RPL41; UBA52; RPL4; RPL38; RPL37A; RPL37; RPL36; RPL35A; RPL35; RPL34; RPL32; RPL31; RPL30; RPS7; RPS6; RPS5; RPS3A; FAU; RPS3; RPS29; RPS28; RPS27A; RPS4Y2; RPS4X; RPS4Y1; RPS26; RPS25; RPS24; RPS27L; RPS27; RPS23; RPS21; RPS20; RPS2; RPS15A; RPS19; RPS18; RPS17; RPS16; RPS15; RPS14; RPS13; RPS12; RPS11; RPSA; RPS10; RPS9; RPS8; GSPT2; GSPT1; ETF1; N6AMT1; TRMT112; APEH; 28S rRNA; 5.8s rRNA; 5s rRNA; 18s rRNA* |
|  | Mitochondrial translation | *MRPL41; MRPL21; MRPL53; MRPL39; MRPL57; MRPL45; MRPL55; GADD45GIP1; MRPL36; MRPL11; MRPL28; MRPL46; MRPL13; MRPL44; MRPL3; MRPL20; MRPL37; MRPL18; MRPL34; MRPL38; MRPL1; OXA1L; MRPL32; MRPL15; MRPL40; MRPL14; MRPL16; MRPL17; MRPL19; MRPL4; MRPL22; MRPL33; MRPL48; MRPL2; MRPL51; MRPL54; MRPL27; MRPL52; MRPL30; MRPL10; MRPL43; MRPL47; MRPL50; MRPL23; MRPL24; MRPL9; MRPL49; MRPL12; MRPL35; MRPL58; MRPL42; DAP3; MRPS18A; MRPS18B; MRPS18C; MRPS11; MRPS15; MRPS33; PTCD3; MRPS16; MRPS2; MRPS12; MRPS6; MRPS10; MRPS7; MRPS34; MRPS35; MRPS23; MRPS5; MRPS17; MRPS36; MRPS25; AURKAIP1; MRPS28; MRPS26; MRPS31; MRPS21; MRPS9; MRPS30; MRPS22; MRPS14; ERAL1; MRPS27; MRPS24; CHCHD1; TUFM; TSFM; GFM1; MTRF1L; MRRF; GFM2; MTFMT; MTIF3; MTIF2; MT-TV; MT-RNR2; MT-RNR1* |
|  | protein folding | *CCT8; TCP1; CCT2; CCT7; CCT5; CCT3; CCT6B; CCT6A; CCT4; ARFGEF2; FBXW2; USP11; XRN2; FBXO6; FBXO4; KIFC3; GAPDHS; FBXW4; FBXW7; FBXW10; FKBP9; FBXW5; NOP56; KIF13A; GBA; FBXL5; SKIV2L; FBXL3; AP3M1; LONP2; FBXW9; WRAP53; TP53; STAT3; HDAC3; DCAF7; CCNE2; CCNE1; SPHK1; PFDN1; VBP1; PFDN4; PFDN5; PFDN2; PFDN6; TUBA1C; TUBB2B; TUBA4A; TUBB3; TUBB6; TUBA1A; ACTB; TUBB1; TUBA3C; TUBA3D; TUBB2A; TUBB4B; TUBB4A; TUBA1B; TUBA8; TUBA4B; TUBAL3; TUBA3E; GNB5; GNB3; GNB2; GNB4; GNB1; PDCL; GNG8; GNGT2; GNGT1; GNG7; GNG13; GNG3; GNG10; GNG11; GNG4; GNG5; GNG12; GNG2; RGS11; RGS6; RGS7; RGS9; GNAT1; GNAI3; GNAI1; GNAZ; GNAI2; GNAT2; GNAO1; GNAT3; CSNK2A1; CSNK2A2; CSNK2B; TBCA; TBCB; TBCD; TBCE; TBCC; ARL2* |
|  | Gamma carboxylation, hypusine formation and arylsulfatase activation | *FN3KRP; DHPS; EIF5A2; EIF5A; DOHH; FN3K; TPST2; TPST1; F8; SUMF1; ARSG; ARSJ; ARSB; ARSD; ARSL; STS; ARSF; ARSH; ARSA; ARSK; ARSI; SUMF2; PROC; F2; PROS1; PROZ; GAS6; F7; BGLAP; F9; F10; FURIN; GGCX; DPH5; EEF2; DPH6; DPH1; DPH2; DPH3; DNAJC24; DPH7; ICMT* |
|  | Post-translational modification: synthesis of GPI-anchored proteins | *PIGF; PIGG; PIGB; DPM2; PIGC; PIGH; PIGA; PIGP; PIGY; PIGQ; PIGW; PIGL; PIGV; PIGM; PIGX; PIGN; PIGO; PIGZ; PIGU; PIGS; GPAA1; PIGK; PIGT; PLAUR; PGAP1; DPM1; DPM3; GPLD1; LY6E; BST1; LY6K; LYPD4; FCGR3B; ULBP2; IZUMO1R; CNTN4; NTNG1; RAET1L; VNN3; ART4; XPNPEP2; LYPD2; GP2; TECTA; MSLN; NRN1; LYPD6B; PLET1; NTM; PRSS21; NEGR1; MELTF; MDGA1; NTNG2; LYPD8; CD52; NRN1L; CD109; ALPI; ART3; SPRN; SPACA4; CEACAM7; OTOA; CNTN5; CPM; OPCML; PRSS41; LY6G6C; LY6D; RTN4RL2; PRND; LY6H; VNN1; RECK; PSCA; LSAMP; ALPG; LYPD3; VNN2; LY6G6D; LYPD1; LYPD5; ALPL; GPIHBP1; THY1; CNTN3; TEX101; RTN4RL1; TECTB; CEACAM5; FOLR2; RAET1G; MDGA2* |
|  | Asparagine N-linked glycosylation | *FUT3; CHST8; LHB; CGA; CHST10; MAN2A1; MAN2A2; FUCA1; MGAT2; FUT8; ST6GAL1; MGAT4A; MGAT4C; MGAT4B; B4GALT1; B4GALT6; B4GALT3; B4GALT4; B4GALT5; B4GALT2; MGAT5; ST8SIA3; ST8SIA6; ST8SIA2; ST3GAL4; MGAT3; LMAN1; MCFD2; NSF; GOSR2; STX5; GOSR1; BET1L; BET1; YKT6; NAPG; NAPB; NAPA; ANK2; ANK1; ANK3; TMED3; TMED2; TMED10; TMED9; TMED7; SPTBN1; SPTBN2; SPTB; SPTBN5; SPTBN4; SPTAN1; SPTA1; COPE; COPB2; COPA; COPZ2; COPZ1; COPB1; COPG1; COPG2; ARCN1; ARF3; ARF1; ARF5; ARF4; RAB1A; RAB1B; GBF1; USO1; KDELR3; KDELR2; INS; KDELR1; CD55; FOLR1; CD59; ARFGAP1; ARFGAP3; ARFGAP2; GORASP1; GOLGA2; COG3; COG7; COG5; COG6; COG4; COG1; COG2; COG8; TMEM115; GOLGB1; TUBB6; TUBB2A; TUBB4B; TUBB1; TUBB3; TUBB2B; TUBB8; TUBB4A; TUBB8B; TUBA8; TUBA4B; TUBAL3; TUBA3E; TUBA1B; TUBA1A; TUBA1C; TUBA3C; TUBA3D; TUBA4A; DYNC1I2; DYNC1I1; DYNC1H1; DYNC1LI1; DYNC1LI2; DYNLL1; DYNLL2; DCTN4; DCTN6; DCTN5; ACTR10; ACTR1A; CAPZA3; CAPZB; CAPZA2; CAPZA1; DCTN2; DCTN3; DCTN1; SEC24C; PREB; MIA2; COL7A1; MIA3; SEC23A; SEC22B; SEC24B; SEC24A; F5; F8; LMAN2L; LMAN2; LMAN1L; CTSC; CTSZ; SERPINA1; SEC24D; TGFA; AREG; CNIH1; GRIA1; CNIH2; CNIH3; SAR1B; SEC22A; SEC22C; SCFD1; STX17; PPP6C; ANKRD28; PPP6R3; PPP6R1; TRAPPC1; TRAPPC10; TRAPPC6B; TRAPPC4; TRAPPC5; TRAPPC6A; TRAPPC3; TRAPPC2L; TRAPPC2; TRAPPC9; SEC31A; SEC13; SEC23IP; TFG; SEC16B; SEC16A; TBC1D20; CSNK1D; MAN1A1; MAN1A2; MAN1C1; MANEA; MGAT1; ASGR1; ASGR2; TUSC3; MAGT1; DAD1; RPN1; DDOST; STT3A; RPN2; ALG9; MPDU1; RFT1; ALG2; DPAGT1; ALG3; ALG8; ALG1; TSTA3; SLC35C1; FCSK; GMDS; FUOM; FPGT; MVD; DOLK; SRD5A3; NUS1; DHDDS; DOLPP1; GNPNAT1; GFPT1; GFPT2; RENBP; AMDHD2; UAP1; NAGK; PGM3; GNE; NANP; ST3GAL5; ST3GAL2; ST3GAL1; ST3GAL6; ST3GAL3; CMAS; ST6GALNAC3; ST6GALNAC6; ST6GALNAC5; ST6GALNAC4; ST6GALNAC2; ST6GALNAC1; ST8SIA4; ST8SIA1; ST8SIA5; NANS; SLC17A5; SLC35A1; ST6GAL2; NEU1; CTSA; GLB1; NEU4; NEU2; NEU3; NPL; DPM1; DPM2; DPM3; GMPPB; GMPPA; MPI; PMM2; PMM1; ALG5; NUDT14; ALG11; ALG12; ALG10B; ALG10; ALG14; ALG13; ALG6; B4GALNT2; UMOD; MLEC; PRKCSH; GANAB; ENGASE; UBC; UBB; UBA52; RPS27A; AMFR; PSMC1; NGLY1; RAD23B; UBXN1; DERL1; VCP; MOGS; CANX; CALR; PDIA3; EDEM2; MAN1B1; EDEM3; EDEM1; UGGT2; UGGT1; SEL1L; OS9; SYVN1; RNF5; RNF139; TRIM13; RNF103; RNF185; MARCHF6; DERL2* |
|  | O-linked glycosylation | *B3GLCT; ADAMTS8; ADAMTS14; THBS2; SBSPON; ADAMTS3; SSPOP; ADAMTS20; ADAMTS17; SEMA5B; THSD4; ADAMTS16; THSD1; ADAMTSL2; ADAMTSL3; ADAMTS15; ADAMTS1; ADAMTS12; ADAMTS9; SPON2; ADAMTS10; ADAMTSL5; ADAMTS18; ADAMTS19; ADAMTS4; SEMA5A; ADAMTS6; THSD7A; THSD7B; ADAMTS2; SPON1; ADAMTSL1; CFP; ADAMTS7; ADAMTS5; ADAMTSL4; ADAMTS13; THBS1; POFUT2; A4GNT; MUC16; MUC1; MUC3A; MUC21; MUC3B; MUC7; MUC17; MUC13; MUC19; MUC4; MUC5AC; MUC20; MUC2; MUC12; MUC5B; MUC6; MUCL1; MUC15; ST6GAL1; ST3GAL2; ST3GAL1; ST3GAL4; ST3GAL3; ST6GALNAC2; ST6GALNAC3; ST6GALNAC4; C1GALT1; C1GALT1C1; GALNTL6; GALNT16; GALNT17; GALNT18; GALNTL5; GALNT4; GALNT5; GALNT15; GALNT6; GALNT10; GALNT14; GALNT1; GALNT8; GALNT7; GALNT12; GALNT13; GALNT3; GALNT11; GALNT9; GALNT2; B4GALT6; B4GALT5; GCNT3; GCNT7; GCNT4; GCNT1; B3GNT9; B3GNTL1; B3GNT2; B3GNT7; B3GNT8; B3GNT4; B3GNT5; B3GNT3; B3GNT6; CHST4; POMT2; POMT1; DAG1; LARGE2; B4GAT1; LARGE1; B3GALNT2; POMGNT1; POMGNT2; POMK* |
|  | SUMOylation | *PIAS1; L3MBTL2; UBE2I; SUMO2; BMI1; PCGF2; SCMH1; RNF2; RING1; CBX2; CBX8; CBX4; PHC1; PHC3; PHC2; SUMO1; H4C1; ZBED1; CHD3; NUP98; SEH1L; NUP107; NUP43; NUP160; NUP37; NUP85; NUP133; SEC13; TPR; NUP153; NUP88; NUP188; NUP214; NDC1; NUP210; POM121; POM121C; NUP35; NUP93; NUP155; NUP205; SEH1L; RAE1; NUP98; NUP98; NUP62; NUP58; NUP58; NUP54; NUP50; AAAS; RANBP2; NUP42; HDAC4; SATB1; PIAS2; SUZ12; HDAC2; SUMO3; HDAC1; SATB2; CBX5; DNMT1; DNMT3A; DNMT3B; HNRNPK; NOP58; HNRNPC; TFAP2A; FOXL2; PIAS3; MITF; SP3; TFAP2B; PIAS4; TP53BP1; CDKN2A; MDM2; TP53; PIAS2; HIC1; MTA1; TFAP2C; RORA; THRB; NR1I2; PPARA; NR4A2; PPARG; RARA; NR1H4; PGR; NR5A1; ESR1; NR2C1; VDR; NR3C2; THRA; NR3C1; AR; NR5A2; NR1H3; NR1H2; RXRA; TDG; MDC1; PARP1; BRCA1; PML; NSMCE2; NSMCE4A; EID3; NSMCE1; SMC5; NSMCE3; SMC6; RAD21; SMC1A; SMC3; STAG2; STAG1; RAD52; CETN2; BLM; XRCC4; HDAC7; SP100; WRN; HERC2; RNF168; XPC; RPA1; PCNA; TOP2A; TOP1; TOP2B; AURKA; RANGAP1; INCENP; AURKB; BIRC5; CDCA8; NFKBIA; RELA; IKBKG; TOPORS; IKBKE; NFKB2; TRIM28; ZNF350; ING2; NCOA2; NPM1; NCOR2; DDX17; PARK7; CREBBP; CASP8AP2; NRIP1; DDX5; EP300; CTBP1; SAFB; MRTFA; DAXX; SIN3A; PPARGC1A; HIPK2; NCOA1; MBD1; UHRF2; ZNF131; VHL; TRIM27; UBA2; SAE1; RWDD3; SENP5; SENP1; SENP2* |
|  | Deubiquitination | *PRKN; UBB; UBC; UBA52; RPS27A; ATXN3; RAD23A; RAD23B; VCP; ATXN3L; JOSD2; JOSD1; KDM1B; HCFC1; MBD6; MBD5; BAP1; ASXL1; ASXL2; FOXK2; FOXK1; H2AC7; H2AC4; H2AC12; H2AC1; H2AC11; H2AC6; H2AC14; H2AC20; H2AC18; H2AC21; H2AW; BARD1; PSMD3; PSMA6; PSMD2; PSMA5; PSMD13; PSMA4; PSMD12; PSMA3; PSMD11; PSMA2; SEM1; PSMD10; PSMA1; PSMD1; PSMD14; PSMC6; PSMC5; PSMC4; PSMC3; PSMC2; PSMC1; PSMB9; PSMB8; PSMB11; PSME4; PSMB7; PSMB6; PSMA8; PSMF1; PSME3; PSMB5; PSME2; PSMB4; PSME1; PSMD9; PSMB3; PSMD8; PSMB2; PSMD7; PSMD6; PSMB10; PSMD5; PSMB1; PSMD4; PSMA7; ADRM1; UCHL5; YY1; OGT; UCHL1; UCHL3; USP15; SMAD7; TGFBR2; TGFB1; TGFBR1; INO80D; INO80E; ACTR5; INO80B; ACTR8; MCRS1; TFPT; INO80C; ACTB; RUVBL1; INO80; NFRKB; ACTL6A; SENP8; NEDD8; EP300; KAT2B; MYSM1; STAMBP; STAM; BABAM1; BABAM2; BRCC3; UIMC1; ABRAXAS1; BRCA1; STAMBPL1; ABRAXAS2; NLRP3; USP30; IDE; VDAC1; PTRH2; VDAC3; VDAC2; MUL1; FKBP8; MAT2B; TOMM70; RHOT1; TOMM20; USP5; SNX3; CFTR; USP10; USP21; USP16; DDX58; RIPK1; H2BC18; H2BC11; H2BC17; H2BC12; H2BC5; H2BC1; H2BC13; H2BC9; H2BC3; H2BC14; H2BC15; H2BC4; H2BC21; H2BU1; TADA2B; TAF9B; TADA3; TRRAP; ATXN7; USP22; KAT2A; TAF10; USP3; USP13; BECN1; USP24; DDB2; USP48; TRAF2; GATA3; IL33; USP11; NFKBIA; SIAH2; USP44; CDC20; SMAD4; USP9X; USP37; CCNA1; CCNA2; USP33; ARRB1; CCP110; ARRB2; USP47; POLB; IKBKG; TRAF6; CYLD; SMAD2; SMAD1; SMAD3; KEAP1; SMURF2; USP42; TP53; USP25; USP2; MDM2; MDM4; USP34; AXIN1; AXIN2; RNF146; TNKS; TNKS2; USP20; ADRB2; USP4; USP49; USP14; USP19; RNF123; USP17L17; USP17L22; USP17L5; USP17L1; USP17L12; USP17L4; USP17L3; USP17L15; USP17L8; USP17L24; USP17L21; USP17L13; USP17L18; USP17L19; USP17L11; USP17L10; USP17L20; USP17L2; RCE1; IFIH1; CDC25A; USP26; WDR20; WDR48; USP12; AR; FOXO4; PTEN; USP7; UFD1; SKP2; MYC; CLSPN; USP28; SUDS3; USP8; HGS; STAM2; HIF1A; BIRC3; BIRC2; RNF128; OTUB1; USP18; TAB1; MAP3K7; ZRANB1; TNFAIP3; OTUD7B; UBE2D1; ESR1; OTUB2; NOD2; NOD1; RIPK2; YOD1; TNIP3; TNIP1; TNIP2; CDK1; VCPIP1; RHOA; TRAF3; OTUD5; RNF135; TRIM25; MAVS; APC; OTUD7A; OTUD3* |
|  | Protein ubiquitination | *HLA-A; HLA-A; HLS-B60; US11; SELENOS; TMEM129; VCP; UBE2J2; UBC; UBB; UBA52; RPS27A; DERL1; UBE2J2; UBE2N; RNF152; RRAGA; UBE2V2; HLTF; RAD18; UBE2B; PCNA; UBE2D2; UBE2D1; UBE2D3; UBE2E1; BCL10; RNF181; PEX5; PEX5; PEX10; PEX2; PEX12; PEX13; PEX14; SHPRH; UBE2L3; RNF144A; PRKDC; H2BC4; H2BC15; H2BC13; H2BC17; H2BC12; H2BC9; H2BC5; H2BC3; H2BC1; H2BC11; H2BC14; CTR9; RTF1; PAF1; CDC73; LEO1; WDR61; UBE2A; RNF40; RNF20; WAC; UBA1; UBA6; UBE2S; UBE2T; UBE2C; CDC34; UBE2E3; UBE2W; UBE2R2; UBE2G1; UBE2K; UBE2Q2; UBE2G2; UBE2H; OTULIN; USP5; USP7; UCHL3; USP9X; UBE2Z* |
|  | RAB geranylgeranylation | *RAB2B; RAB29; RAB19; RAB3B; RAB24; RAB40B; RAB33A; RAB42; RAB27B; RAB7B; RAB4A; RAB20; RAB32; RAB8B; RAB44; RAB3D; RAB39A; RAB14; RAB33B; RAB18; RAB17; RAB41; RAB31; RAB35; RAB23; RAB40C; RAB40A; RAB36; RAB4B; RAB15; RAB25; RAB3C; RAB26; RAB22A; RAB21; RAB12; RAB34; RAB30; RAB37; RAB39B; RAB10; RAB9A; RAB9B; RAB43; RAB27A; RAB5B; RAB5C; RAB7A; RAB6A; RAB6B; RAB11B; RAB5A; RAB8A; RAB2A; RAB11A; RAB38; RAB1B; RAB1A; RAB3A; RAB13; CHML; CHM; RABGGTB; RABGGTA; PTP4A2* |
|  | Protein methylation | *PRMT3; RPS2; METTL21A; HSPA8; EEF1AKMT2; EEF1A1; ETFBKMT; ETFB; CAMKMT; CALM1; EEF2KMT; EEF2; METTL22; KIN; VCPKMT; VCP; EEF1AKMT1* |
|  | Carboxyterminal post-translational modifications of tubulin | *TTL; TUBB6; TUBB2A; TUBB4B; TUBB1; TUBB3; TUBB2B; TUBB8; TUBB4A; TUBB8B; TUBA1A; TUBA1C; TUBA1B; TUBA3D; TUBA3C; TUBA3E; TTLL1; TPGS1; TPGS2; LRRC49; NICN1; TUBA8; TUBA4B; TUBAL3; TUBA4A; TTLL9; TTLL12; TTLL2; TTLL7; TTLL6; TTLL11; TTLL4; TTLL5; TTLL13P; TTLL10; TTLL3; TTLL8; AGBL5; AGTPBP1; AGBL3; AGBL4; AGBL2; AGBL1* |
|  | Neddylation | *RBX1; CUL9; NEDD8; UBE2M; COMMD5; COMMD6; COMMD2; COMMD10; COMMD8; COMMD9; COMMD4; COMMD3; COMMD7; COMMD1; CCDC22; DDB1; RBBP7; ERCC8; DDA1; DCAF17; COP1; DCAF8; DCAF16; DCAF10; DDB2; DCAF6; DCAF11; RBBP5; WDTC1; DCAF13; WDR5; DCAF7; DCAF5; DTL; DCAF4; CUL4A; CUL4B; DCUN1D2; DCUN1D4; DCUN1D5; DCUN1D1; BIRC5; UBC; UBB; UBA52; RPS27A; OBSL1; CCDC8; CUL7; DCUN1D3; UBE2F; CUL5; SPSB2; ASB3; NEURL2; ASB17; ASB6; ASB11; ASB12; ASB10; ASB16; ASB4; SPSB3; ASB18; ASB14; WSB2; ASB5; ASB2; ASB8; CISH; SOCS2; TULP4; WSB1; SPSB4; SPSB1; SOCS5; SOCS3; ASB7; ASB9; ASB13; ASB15; ASB1; SOCS6; ANKRD9; LRRC41; ELOC; ELOB; RNF7; UCHL3; SENP8; NAE1; UBA3; CAND1; UBD; PSMD3; PSMA6; PSMD2; PSMA5; PSMD13; PSMA4; PSMD12; PSMA3; PSMD11; PSMA2; SEM1; PSMD10; PSMA1; PSMD1; PSMD14; PSMC6; PSMC5; PSMC4; PSMC3; PSMC2; PSMC1; PSMB9; PSMB8; PSMB11; PSME4; PSMB7; PSMB6; PSMA8; PSMF1; PSME3; PSMB5; PSME2; PSMB4; PSME1; PSMD9; PSMB3; PSMD8; PSMB2; PSMD7; PSMD6; PSMB10; PSMD5; PSMB1; PSMD4; PSMA7; NUB1; NUB1; CUL1; SKP1; FBXW9; FBXL3; LMO7; FBXL5; FBXL20; FBXW5; FBXW11; FBXO17; FBXO27; FBXL18; FBXL8; FBXL15; FBXO7; FBXO2; FBXW8; FBXL13; BTRC; FBXO10; FBXO30; FBXO11; FBXO15; FBXO41; FBXO31; FBXO9; FBXL7; FBXL16; FBXL12; FBXL4; FBXO40; FBXW12; FBXO44; FBXO22; FBXW10; FBXW7; CCNF; FBXW4; FBXL14; FBXL21P; SKP2; FBXL19; FBXO21; FBXO4; FBXO6; FBXL22; FBXW2; FBXO32; CUL2; FEM1A; FEM1B; FEM1C; VHL; LRR1; CUL3; GAN; ZBTB16; KLHL11; KLHL42; BTBD1; KBTBD13; KLHL13; KEAP1; KLHL22; KLHL20; BTBD6; KLHL5; KLHL41; KLHL25; KBTBD6; KLHL21; KLHL9; KLHL2; KCTD7; KBTBD7; KLHL3; KCTD6; KBTBD8; HIF1A; HIF3A; EPAS1; UBXN7; COPS8; COPS2; COPS6; GPS1; COPS7A; COPS7B; COPS4; COPS5; COPS3; UBE2D3; UBE2D2; UBE2D1; PUM2; DCUN1D3* |
|  | Post-translational protein phosphorylation | *APOA1; VCAN; TNC; SPP2; GAS6; GPC3; CHGB; MBTPS1; TMEM132A; SERPIND1; PRKCSH; IGFBP7; BMP15; SCG2; LGALS1; GOLM1; SERPINA1; FN1; APOL1; LAMB1; STC2; AMBN; MATN3; AMELX; C4A; APOA5; MFGE8; FGA; SPP1; LAMC1; SCG3; CCN1; KTN1; MXRA8; FGG; TIMP1; ALB; FSTL3; TGOLN2; MELTF; MSLN; IGFBP1; DMP1; FGF23; APP; P4HB; PDIA6; F5; HSP90B1; APOB; PENK; FBN1; FUCA2; MEPE; PCSK9; NOTUM; WFS1; IGFBP3; FSTL1; HRC; NUCB1; CP; CKAP4; IGFBP5; SERPINA10; CST3; SDC2; ANO8; DNAJC3; MIA3; PRSS23; CDH2; AHSG; TF; IL6; QSOX1; KNG1; LAMB2; VWA1; IGFBP4; BPIFB2; CHRDL1; SPARCL1; RCN1; AFP; PNPLA2; ENAM; CALU; CSF1; AMTN; APOA2; MGAT4A; SHISA5; ITIH2; ADAM10; VGF; LTBP1; APOE; EVA1A; PROC; SERPINC1; C3; BMP4; APLP2; MEN1; FAM20C; FAM20A* |
|  | Peptide hormone metabolism | *PCSK1; GHRL; SPCS2; SPCS3; SEC11A; SEC11C; SPCS1; GHRL; GH1; GCG; INS; LEP; IGF1; MBOAT4; ACHE; BCHE; PLA2G7; KLF4; UCN; CRHR2; POMC; CGA; CGB3; LHB; FSHB; INHA; INHBC; INHBE; INHBA; INHBB; TSHB; AOPEP; AGT; MME; ATP6AP2; CTSD; REN; ACE; CPB2; CPB1; CPA3; ACE2; GZMH; CTSG; CTSZ; CMA1; ANPEP; CES1; ENPEP; CPE; EXOC8; EXOC5; EXOC2; EXOC6; EXOC3; EXOC1; EXOC4; EXOC7; SLC30A5; SLC30A8; SLC30A6; SLC30A7; VAMP2; STX1A; CLTRN; KIF5A; KIF5C; KIF5B; MYRIP; MYO5A; RAB27A; PCSK2; ERO1B; ERO1A; GIP; ISL1; GATA4; PAX6; DPP4; GPR119; FFAR1; CTNNB1; TCF7L2; CDX2; GRP; FFAR4; GNB3; GNG13; GNAT3; GHRL; GIP; GCG* |
|  | Regulation of Insulin-like Growth Factor (IGF) transport and uptake by Insulin-like Growth Factor Binding Proteins (IGFBPs) | *IGF1; IGF2; IGFBP2; IGFBP4; F2; IGFBP3; IGFALS; PAPPA; PAPPA2; IGFBP5; APOA1; VCAN; TNC; SPP2; GAS6; GPC3; CHGB; MBTPS1; TMEM132A; SERPIND1; PRKCSH; IGFBP7; BMP15; SCG2; LGALS1; GOLM1; SERPINA1; FN1; APOL1; LAMB1; STC2; AMBN; MATN3; AMELX; C4A; APOA5; MFGE8; FGA; SPP1; LAMC1; SCG3; CCN1; KTN1; MXRA8; FGG; TIMP1; ALB; FSTL3; TGOLN2; MELTF; MSLN; IGFBP1; DMP1; FGF23; APP; P4HB; PDIA6; F5; HSP90B1; APOB; PENK; FBN1; FUCA2; MEPE; PCSK9; NOTUM; WFS1; FSTL1; HRC; NUCB1; CP; CKAP4; SERPINA10; CST3; SDC2; ANO8; DNAJC3; MIA3; PRSS23; CDH2; AHSG; TF; IL6; QSOX1; KNG1; LAMB2; VWA1; BPIFB2; CHRDL1; SPARCL1; RCN1; AFP; PNPLA2; ENAM; CALU; CSF1; AMTN; APOA2; MGAT4A; SHISA5; ITIH2; ADAM10; VGF; LTBP1; APOE; EVA1A; PROC; SERPINC1; C3; BMP4; APLP2; MEN1; FAM20C; FAM20A; PLG; GZMH; CTSG; IGFBP6; MMP2; MMP1; KLK13; KLK2; KLK1; KLK3* |
|  | Unfolded Protein Response (UPR) | *MBTPS1; CREB3L1; CREBRF; CREB3; CREB3L4; CREB3L3; CREB3L2; MBTPS2; DCSTAMP; EIF2AK3; EIF2S2; EIF2S3; EIF2S1; CXCL8; ATF4; EXOSC3; EXOSC1; EXOSC9; EXOSC6; EXOSC5; EXOSC7; EXOSC4; EXOSC8; DIS3; EXOSC2; KHSRP; DCP2; PARN; ASNS; CEBPG; CEBPB; IGFBP1; DDIT3; NFYA; NFYB; NFYC; ATF6; CCL2; HERPUD1; ATF3; HSPA5; ERN1; XBP1; ADD1; YIF1A; TLN1; TPP1; SRPRA; DNAJB11; SYVN1; EXTL3; PREB; EXTL1; TATDN2; DCTN1; DNAJC3; CXXC1; GFPT1; GOSR2; EDEM1; SSR1; MYDGF; SERP1; PPP2R5B; WIPI1; SULT1A3; WFS1; DDX11; GSK3A; HDGF; KLHDC3; FKBP14; SRPRB; DNAJB9; ATP6V0D1; TSPYL2; ZBTB17; PDIA5; CUL7; KDELR3; SEC31A; ACADVL; SHC1; PLA2G4B; EXTL2; LMNA; ARFGAP1; HYOU1; CTDSP2; PDIA6; HSP90B1; CALR; IL8; ASNS; IGFBP1; DDIT3; CCL2; HERPUD1; ATF3; ATF4; (unspliced); (spliced); ADD1; YIF1A; TLN1; TPP1; SRPR; DNAJB11; SYVN1; EXTL3; PREB; EXTL1; TATDN2; DCTN1; DNAJC3; CXXC1; GFPT1; GOSR2; EDEM1; SSR1; C19orf10; SERP1; PPP2R5B; WIPI1; SULT1A3; WFS1; DDX11; GSK3A; HDGF; KLHDC3; gene; SRPRB; DNAJB9; ATPV0D1; TSPYL2; ZBTB17; PDIA5; CUL7; KDELR3; SEC31A; ACADVL; SHC1; PLA2G4B; EXTL2; LMNA; ARFGAP1; HYOU1; CTDSP2; PDIA6; HSP90B1; CALR; HSPA5; XBP1* |
|  | Protein repair | *PCMT1; MSRB2; MSRB3; MSRB1; TXN; MSRA* |
|  | Surfactant metabolism | *SFTPD; ZDHHC2; CKAP4; ADA2; SFTPA2; SFTPA1; SFTA3; GATA6; ADORA2A; ADORA2B; SFTPC; CCDC59; TTF1; CSF2RB; CSF2RA; SFTPB; ADGRF5; ADRA2C; ADRA2A; P2RY2; LMCD1; DMBT1; PGA4; PGA3; CTSH; NAPSA; PGA5; ABCA3; SLC34A1; SLC34A2; SFTPD; SFTA3; SFTPA1; SFTPA2; SFTPC; SFTPB* |
|  | Amyloid fiber formation | *SORL1; APP; UBE2L6; SNCA; SIAH1; USP9X; UBC; UBB; UBA52; RPS27A; APCS; H2BC1; H2BC9; H2BC13; H2BC12; H2BU1; H2BC21; H2BC3; H2BC4; H2BC11; H2BC14; H2BC5; H2BC15; H2BS1; H2BC17; H2AX; H2AC20; H2AC6; H2AZ1; H2AB1; H2AC14; H2AC7; H2AC4; H2AC18; H3-3A; H3C15; H3C1; H4C1; SIAH2; BACE1; ADAM10; TSPAN5; TSPAN14; TSPAN15; TSPAN33; FURIN; GGA1; GGA3; GGA2; FGA; LYZ; APOA1; APOA4; GSN; CST3; ITM2B; SAA1; B2M; TTR; PRL; ODAM; LTF; NPPA; MFGE8; SEMG1; TGFBI; IAPP; CALCA; INS; HSPG2; PRKN; SNCAIP; NAT8B; NAT8; SNCAIP; PSENEN; NCSTN; APH1A; APH1B; CALB1* |
| Metabolism  of RNA | mRNA Capping | *RNGTT; GTF2H4; ERCC3; GTF2H3; GTF2H2; GTF2H1; GTF2H5; CDK7; MNAT1; CCNH; ERCC2; POLR2B; POLR2E; POLR2K; POLR2C; POLR2L; POLR2H; POLR2F; POLR2D; POLR2I; POLR2J; POLR2A; POLR2G; GTF2F1; GTF2F2; SUPT5H; RNMT; NCBP2; NCBP1* |
|  | Processing of Capped Intron-Containing Pre-mRNA | *HNRNPF; RBMX; HNRNPH1; HNRNPH2; HNRNPK; HNRNPL; HNRNPM; HNRNPR; PCBP1; HNRNPU; PCBP2; PTBP1; NCBP2; NCBP1; GTF2F1; GTF2F2; POLR2B; POLR2E; POLR2K; POLR2C; POLR2L; POLR2H; POLR2F; POLR2D; POLR2I; POLR2J; POLR2G; POLR2A; HNRNPA0; HNRNPA1; HNRNPA2B1; HNRNPA3; HNRNPC; HNRNPD; DDX39B; DDX39A; POLDIP3; FYTTD1; LUZP4; ALYREF; ZC3H11A; SRSF2; SRSF3; SRSF9; SLU7; U2AF2; CLP1; PCF11; CSTF1; CSTF2; CSTF2T; CSTF3; PABPN1; CPSF4; SYMPK; CPSF1; CPSF2; CPSF3; WDR33; FIP1L1; PAPOLA; NUDT21; CPSF7; RNPS1; UPF3B; RBM8A; MAGOH; MAGOHB; EIF4A3; CASC3; SRRM1; SRSF4; U2AF1; U2AF1L4; SRSF11; SRSF6; SRSF7; SRSF5; DHX38; SRSF1; CDC40; CHTOP; SARNP; THOC5; THOC3; THOC2; THOC1; THOC7; THOC6; NXF1; NUP98; SEH1L; NUP107; NUP43; NUP160; NUP37; NUP85; NUP133; SEC13; TPR; NUP153; NUP88; NUP188; NUP214; NDC1; NUP210; POM121; POM121C; NUP35; NUP93; NUP155; NUP205; SEH1L; RAE1; NUP98; NUP98; NUP62; NUP58; NUP58; NUP54; NUP50; AAAS; RANBP2; NUP42; EIF4E; SLBP; NXF2; NXT1; GLE1; CCAR1; SRRM2; RBM5; AQR; SRRT; ISY1; SUGP1; FUS; YBX1; PPIE; XAB2; DNAJC8; PRCC; CWC25; CRNKL1; ELAVL2; BUD31; PPWD1; HSPA8; CWC15; BCAS2; CDC5L; PQBP1; PLRG1; WBP11; PRPF19; CTNNBL1; ELAVL1; SMNDC1; RBM17; SF3B1; SF3B6; DDX42; SF3B2; SF3B3; SF3B4; SF3B5; CHERP; SNRPE; SNRPF; SNRPG; SNRPD1; SNRPB; SNRPN; SNRPD3; SNRPD2; PUF60; DDX46; SNRPA1; DHX15; SNRPB2; U2SURP; PHF5A; SF3A1; SF3A2; SF3A3; TRA2B; TXNL4A; PRPF8; SNRNP40; DDX23; LSM2; PRPF6; EFTUD2; SNRNP200; SNU13; SYF2; HNRNPUL1; SNW1; RBM22; PPIL3; PPIL6; PPIL4; PPIL1; CWC22; DDX5; DHX16; GPKOW; DHX9; CD2BP2; MTREX; GCFC2; TFIP11; SNRPC; SNRNP70; SNRPA; SART1; SNRNP27; PRPF3; PRPF4; PPIH; PRPF31; LSM3; LSM4; LSM7; LSM5; LSM8; LSM6; USP39; PRPF38A; WBP4; CWC27; SF1; PRPF40A; SRSF10; RNPC3; ZCRB1; ZMAT5; SNRNP25; SNRNP35; PDCD7; SNRNP48; ZRSR2; WTAP; METTL3; METTL14; snRNA; snRNA; snRNA; snRNA; snRNA; RNU12; RNU11; RNU4ATAC* |
|  | Processing of Capped Intronless Pre-mRNA | *SLBP; NCBP2; NCBP1; ZNF473; SNRPE; SNRPF; LSM10; SNRPG; SNRPB; LSM11; SNRPD3; CPSF4; SYMPK; CPSF1; CPSF2; CPSF3; WDR33; FIP1L1; CSTF1; CSTF2; CSTF2T; CSTF3; CLP1; PCF11; PABPN1; PAPOLA; NUDT21; CPSF7* |
|  | mRNA Editing | *A1CF; APOBEC3H; APOBEC3C; APOBEC2; APOBEC3B; APOBEC3A; APOBEC4; APOBEC1; ADAR; ADARB1* |
|  | Regulation of mRNA stability by proteins that bind AU-rich elements | *YWHAB; ZFP36L1; AKT1; MAPKAPK2; EXOSC3; EXOSC1; EXOSC9; EXOSC6; EXOSC5; EXOSC7; EXOSC4; EXOSC8; DIS3; EXOSC2; DCP2; XRN1; DCP1A; ZFP36; TNPO1; KHSRP; PARN; MAPK14; MAPK11; YWHAZ; HNRNPD; HNRNPD; PABPC1; HSPB1; HSPA8; EIF4G1; HSPA1A; UBC; UBB; UBA52; RPS27A; PSMD3; PSMA6; PSMD2; PSMA5; PSMD13; PSMA4; PSMD12; PSMA3; PSMD11; PSMA2; SEM1; PSMD10; PSMA1; PSMD1; PSMD14; PSMC6; PSMC5; PSMC4; PSMC3; PSMC2; PSMC1; PSMB9; PSMB8; PSMB11; PSME4; PSMB7; PSMB6; PSMA8; PSMF1; PSME3; PSMB5; PSME2; PSMB4; PSME1; PSMD9; PSMB3; PSMD8; PSMB2; PSMD7; PSMD6; PSMB10; PSMD5; PSMB1; PSMD4; PSMA7; PRKCA; ELAVL1; PRKCD; NUP214; ANP32A; SET; XPO1; TNFSF13; ENPP2; GPRC5A* |
|  | Insulin-like Growth Factor-2 mRNA Binding Proteins (IGF2BPs/IMPs/VICK | *IGF2BP2; IGF2BP3; IGF2BP1; IGF2; IGF2; CD44; CD44; CD44; CD44; CD44; ACTB; H19; MYC* |
|  | Deadenylation-dependent mRNA decay | *NT5C3B; EXOSC3; EXOSC1; EXOSC9; EXOSC6; EXOSC5; EXOSC7; EXOSC4; EXOSC8; DIS3; EXOSC2; HBS1L; WDR61; SKIV2L; TTC37; DCPS; DDX6; DCP2; EDC3; DCP1B; DCP1A; EDC4; LSM3; LSM4; LSM6; PATL1; LSM5; LSM7; LSM1; LSM2; XRN1; PAN3; PAN2; PABPC1; EIF4E; PAIP1; EIF4A3; EIF4A1; EIF4A2; EIF4G1; EIF4B; CNOT4; CNOT10; CNOT1; CNOT3; CNOT2; CNOT7; CNOT8; CNOT9; CNOT11; CNOT6; CNOT6L; TNKS1BP1; TUT4; TUT7; PARN* |
|  | Nonsense-Mediated Decay (NMD) | *GSPT2; GSPT1; ETF1; PABPC1; EIF4G1; NCBP2; NCBP1; RPL29; RPL28; RPL27A; RPL27; RPL3L; RPL3; RPL24; RPL10L; RPL10; RPL23A; RPL23; RPL21; RPL19; RPL22L1; RPL22; RPL18A; RPL18; RPL17; RPL15; RPL14; RPL13A; RPL26L1; RPL26; RPL13; RPL12; RPL11; RPL10A; RPL36AL; RPL36A; RPLP2; RPLP1; RPLP0; RPL39L; RPL39; RPL9; RPL8; RPL7A; RPL7; RPL6; RPL5; RPL41; UBA52; RPL4; RPL38; RPL37A; RPL37; RPL36; RPL35A; RPL35; RPL34; RPL32; RPL31; RPL30; RPS7; RPS6; RPS5; RPS3A; FAU; RPS3; RPS29; RPS28; RPS27A; RPS4Y2; RPS4X; RPS4Y1; RPS26; RPS25; RPS24; RPS27L; RPS27; RPS23; RPS21; RPS20; RPS2; RPS15A; RPS19; RPS18; RPS17; RPS16; RPS15; RPS14; RPS13; RPS12; RPS11; RPSA; RPS10; RPS9; RPS8; UPF1; SMG7; CASC3; EIF4A3; MAGOH; MAGOHB; RBM8A; UPF3A; UPF3B; UPF2; RNPS1; UPF3A; SMG5; SMG6; PPP2R2A; PPP2CA; PPP2R1A; SMG8; SMG9; SMG1; DCP1A; PNRC2; 28S rRNA; 5.8S rRNA; 5S rRNA; 18S rRNA* |
|  | rRNA processing in the nucleus and cytosol | *BUD23; TRMT112; DIMT1; PNO1; NOP2; TSR3; FBL; NOP58; SNU13; NOP56; NAT10; THUMPD1; NOC4L; NOL11; UTP18; WDR3; UTP6; WDR36; TBL3; PWP2; FCF1; DCAF13; KRR1; WDR46; DHX37; RPS14; DDX47; RPS6; EMG1; DDX49; UTP14C; UTP14A; RPS9; RPS2; RRP9; RCL1; BMS1; PDCD11; NOP14; RRP7A; NOL6; RRP36; RPS7; UTP11; MPHOSPH10; IMP4; IMP3; UTP3; DDX52; UTP4; UTP15; HEATR1; WDR43; WDR75; DIEXF; UTP20; NHP2; GAR1; DKC1; NOP10; NOB1; RIOK3; CSNK1D; CSNK1E; RPS16; RPS13; RPP40; RIOK2; RPS24; RPS5; TSR1; RPS15A; RPS21; RPS10; RPS27A; RPS3A; LTV1; RPSA; RPS29; RPS20; RPS4Y2; RPS4Y1; RPS4X; RPS3; RPP30; RPS12; FAU; RPS17; RPP25; RPS15; RPS25; RPS26; RPS19; RPS27L; RPS27; RPS23; RPP14; RPS11; RPS18; RPP21; RPS28; RIOK1; RPS8; RPP38; BYSL; MTREX; EXOSC7; EXOSC6; EXOSC3; EXOSC8; EXOSC4; EXOSC2; EXOSC5; EXOSC1; EXOSC9; MPHOSPH6; EXOSC10; C1D; NIP7; FTSJ3; SENP3; LAS1L; TEX10; PELP1; WDR18; NOL9; PES1; BOP1; WDR12; EBNA1BP2; GNL3; DDX21; RPL29; RPL28; RPL27A; RPL27; RPL3L; RPL3; RPL24; RPL10L; RPL10; RPL23A; RPL23; RPL21; RPL19; RPL22L1; RPL22; RPL18A; RPL18; RPL17; RPL15; RPL14; RPL13A; RPL26L1; RPL26; RPL13; RPL12; RPL11; RPL10A; RPL36AL; RPL36A; RPLP2; RPLP1; RPLP0; RPL39L; RPL39; RPL9; RPL8; RPL7A; RPL7; RPL6; RPL5; RPL41; UBA52; RPL4; RPL38; RPL37A; RPL37; RPL36; RPL35A; RPL35; RPL34; RPL32; RPL31; RPL30; ERI1; ISG20L2; DIS3; NOL12; XRN2; RBM28; RRP1; NCL; RNA45S5; 28S rRNA; SNORD3A; 18S rRNA; 5.8S rRNA; 5S rRNA* |
|  | rRNA processing in the mitochondrion | *TRMT10C; PRORP; HSD17B10; ELAC2; MRM2; MRM1; MTERF4; NSUN4; TFB1M; MRM3; MT-CO2; MT-TG; MT-TV; MT-TK; MT-ND2; MT-TH; MT-TT; MT-ND4L; MT-TR; MT-CYB; MT-TL1; MT-TF; MT-TW; MT-RNR1; MT-ATP6; MT-ND3; MT-ND4; MT-ND5; MT-CO3; MT-ATP8; MT-CO1; MT-TM; MT-TS2; MT-TL2; MT-ND1; MT-TD; MT-RNR2; MT-TI* |
|  | tRNA processing in the nucleus | *TRNT1; RAN; XPOT; FAM98B; RTRAF; DDX1; C2orf49; RTCB; ZBTB8OS; ELAC2; CLP1; CPSF4; CSTF2; TSEN15; CPSF1; TSEN34; TSEN2; TSEN54; RPP38; POP4; POP1; POP7; RPP40; RPP25; POP5; RPP21; RPP14; RPP30; NUP98; SEH1L; NUP107; NUP43; NUP160; NUP37; NUP85; NUP133; SEC13; TPR; NUP153; NUP88; NUP188; NUP214; NDC1; NUP210; POM121; POM121C; NUP35; NUP93; NUP155; NUP205; SEH1L; RAE1; NUP98; NUP98; NUP62; NUP58; NUP58; NUP54; NUP50; AAAS; RANBP2; NUP42; RPPH1* |
|  | tRNA modification in the nucleus and cytosol | *ADAT3; ADAT2; TRMT44; THG1L; LAGE3; OSGEP; TP53RK; TPRKB; PUS1; QTRT2; QTRT1; NSUN2; NSUN6; FTSJ1; THADA; PUS3; TRMT10A; TRMT13; URM1; CTU1; CTU2; TRMT9B; TRMT11; TRMT112; ALKBH8; TRMT61A; TRMT6; ADAT1; WDR4; METTL1; TRIT1; LCMT2; TYW1; TYW3; TYW5; TRMT12; TRMT5; CDKAL1; PUS7; TRDMT1; TRMT1; DUS2; EPRS1* |
|  | tRNA processing in the mitochondrion | *MT-ND6; MT-TA; MT-TN; MT-TP; MT-TS1; MT-TC; MT-TE; MT-TQ; MT-TY; MT-TI; MT-TD; MT-TW; MT-TF; MT-TL2; MT-TL1; MT-TR; MT-TS2; MT-TM; MT-TT; MT-TH; MT-TK; MT-TV; MT-TG; MT-CO2; MT-ND2; MT-ND4L; MT-CYB; MT-RNR1; MT-ATP6; MT-ND3; MT-ND4; MT-ND5; MT-CO3; MT-ATP8; MT-CO1; MT-ND1; MT-RNR2* |
|  | tRNA modification in the mitochondrion | *TRMU; TRMT10C; PRORP; HSD17B10; TRMT61B; TRIT1; PUS1; MTO1; GTPBP3* |
|  | Metabolism of non-coding RNA | *SNRPD2; SNRPG; SNRPE; SNRPD3; SNRPB; SNRPD1; SNRPF; GEMIN8; GEMIN7; GEMIN2; GEMIN6; GEMIN5; DDX20; SMN1; GEMIN4; SNUPN; NCBP2; NCBP1; PHAX; NUP98; SEH1L; NUP107; NUP43; NUP160; NUP37; NUP85; NUP133; SEC13; TPR; NUP153; NUP88; NUP188; NUP214; NDC1; NUP210; POM121; POM121C; NUP35; NUP93; NUP155; NUP205; SEH1L; RAE1; NUP98; NUP98; NUP62; NUP58; NUP58; NUP54; NUP50; AAAS; RANBP2; NUP42; TGS1; PRMT5; CLNS1A; WDR77; U1A snRNA; U2 snRNA; U4 snRNA; U6 snRNA; U5 snRNA* |
| Epigenetic  regulation of  gene  expression | PRC2 methylates histones and DNA | *H2BC14; H2BC1; H2BC5; H2BC9; H2BC21; H2BC17; H2BS1; H2BC15; H2BC4; H2BC11; H2BC3; H2BC12; H2BC13; H2BU1; H3-3A; H3C1; H3C15; H2AB1; H2AZ2; H2AZ1; H2AC7; H2AX; H2AJ; H2AC20; H2AC6; H2AC4; H2AC18; H2AC14; H4C1; SUZ12; EZH2; EED; RBBP7; RBBP4; PHF19; PHF1; JARID2; MTF2; AEBP2; DNMT1; DNMT3A; DNMT3B* |
|  | TET1,2,3 and TDG demethylate DNA | *TDG; TET1; TET2; TET3* |
|  | Positive epigenetic regulation of rRNA expression | *H2BC14; H2BC1; H2BC5; H2BC9; H2BC21; H2BC17; H2BS1; H2BC15; H2BC4; H2BC11; H2BC3; H2BC12; H2BC13; H2BU1; H3C1; H3C15; H3-3A; H2AB1; H2AZ2; H2AZ1; H2AC7; H2AX; H2AJ; H2AC20; H2AC6; H2AC4; H2AC18; H2AC14; H4C1; CBX3; TTF1; EHMT2; ERCC6; RBBP7; GATAD2A; GATAD2B; MBD3; CHD3; CHD4; HDAC2; HDAC1; RBBP4; MTA3; MTA2; MTA1; EP300; KAT2A; SF3B1; SMARCA5; DEK; BAZ1B; MYO1C; MYBBP1A; DDX21; POLR2E; POLR2L; POLR2H; POLR2F; POLR1A; POLR1B; POLR1F; POLR1C; POLR1G; POLR1D; POLR1E; POLR1H; POLR2K; TAF1C; TBP; TAF1A; TAF1B; TAF1D; KAT2B; ACTB; GSK3B; 45S pre-rRNA* |
|  | Negative epigenetic regulation of rRNA expression | *POLR2E; POLR2L; POLR2H; POLR2F; POLR1A; POLR1B; POLR1F; POLR1C; POLR1G; POLR1D; POLR1E; POLR1H; POLR2K; UBTF; GTF2H4; ERCC3; GTF2H3; GTF2H2; GTF2H1; GTF2H5; CDK7; MNAT1; CCNH; ERCC2; TAF1C; TBP; TAF1A; TAF1D; TAF1B; TTF1; SAP130; SUDS3; SAP30BP; SAP30L; SAP30; SIN3B; SIN3A; ARID4B; SAP18; DNMT1; HDAC2; SMARCA5; BAZ2A; HDAC1; DNMT3B; H2BC14; H2BC1; H2BC5; H2BC9; H2BC21; H2BC17; H2BS1; H2BC15; H2BC4; H2BC11; H2BC3; H2BC12; H2BC13; H2BU1; H3C1; H3C15; H3-3A; H2AB1; H2AZ2; H2AZ1; H2AC7; H2AX; H2AJ; H2AC20; H2AC6; H2AC4; H2AC18; H2AC14; H4C1; MBD2; RRP8; SUV39H1; SIRT1; 18S rRNA; 45S pre-rRNA; 28S rRNA; 5S rRNA; 5.8S rRNA* |
|  | DNA methylation | *H2BC14; H2BC1; H2BC5; H2BC9; H2BC21; H2BC17; H2BS1; H2BC15; H2BC4; H2BC11; H2BC3; H2BC12; H2BC13; H2BU1; H3-3A; H3C1; H3C15; H2AB1; H2AZ2; H2AZ1; H2AC7; H2AX; H2AJ; H2AC20; H2AC6; H2AC4; H2AC18; H2AC14; H4C1; DNMT3L; DNMT3B; DNMT1; UHRF1; DNMT3A* |
|  | HATs acetylate histones | *H3C15; H3C1; WDR5; RBBP7; H4C1; H2AC18; H2AC20; H2AC21; H2AC14; H2AC12; H2AC4; H2AC6; H2AC1; H2AC11; H2AC7; H2AW; ACTL6A; ING5; MEAF6; BRD1; BRPF3; BRPF1; KAT6A; KAT6B; DMAP1; EPC1; KAT5; BRD8; MORF4L1; YEATS4; EP400; ING3; MORF4L2; ACTB; RUVBL1; TRRAP; RUVBL2; VPS72; MRGBP; SGF29; TADA3; MBIP; KAT2A; KAT2B; YEATS2; TADA2A; ZZZ3; KAT14; DR1; SUPT7L; TADA2B; SAP130; TADA1; TAF9; USP22; TAF6L; TAF10; SUPT3H; TAF12; TAF5L; ATXN7; ENY2; SUPT20H; ATXN7L3; EP300; H2BC18; H2BU1; H2BC4; H2BC15; H2BC13; H2BC17; H2BC12; H2BC9; H2BC5; H2BC3; H2BC1; H2BC11; H2BC14; H2BC21; CREBBP; MSL1; MSL3; KAT8; MSL2; CLOCK; ELP6; ELP4; ELP5; ELP2; ELP1; ELP3; HAT1; NCOA2; ATF2; ING4; KAT7; JADE2; JADE3; JADE1; KANSL1; KANSL2; OGT; KANSL3; HCFC1; MCRS1; PHF20; PAX3; NCOA1* |
|  | HDACs deacetylate histones | *H3C15; H3C1; RBBP7; RBBP4; H4C1; KDM1A; H2AC18; H2AC20; H2AC21; H2AC14; H2AC12; H2AC4; H2AC6; H2AC1; H2AC11; H2AC7; H2AW; H2BC18; H2BU1; H2BC4; H2BC15; H2BC13; H2BC17; H2BC12; H2BC9; H2BC5; H2BC3; H2BC1; H2BC11; H2BC14; H2BC21; NCOR1; NCOR2; HDAC3; TBL1X; TBL1XR1; GPS2; HDAC8; HDAC10; BRMS1; ARID4A; SAP30L; SUDS3; SAP30; ARID4B; HDAC2; HDAC1; SAP18; RCOR1; REST; GATAD2A; GATAD2B; MBD3; CHD3; CHD4; MTA3; MTA2; MTA1; HMG20B; PHF21A* |
|  | HDMs demethylate histones | *H3C15; H3C1; H4C1; KDM6B; KDM7A; UTY; KDM6A; JMJD6; KDM5C; KDM5A; KDM5B; KDM5D; KDM2A; KDM4A; KDM2B; PHF2; ARID5B; KDM3A; PHF8; KDM3B; KDM1B; KDM1A; KDM4D; KDM4C; KDM4B; RIOX2* |
|  | PKMTs methylate histone lysine | *H3C15; H3C1; ASH2L; RBBP5; WDR5; DPY30; SETD1B; KMT2D; SETD1A; KMT2B; KMT2A; KMT2C; KMT2E; NSD3; SETD3; SMYD3; SETD7; PRDM9; MECOM; PRDM16; AEBP2; SUZ12; EZH2; EED; RBBP7; RBBP4; EHMT2; EHMT1; NSD2; SETD2; KMT5A; H4C1; KMT5B; KMT5C; DOT1L; SETD6; NFKB1; NFKB2; RELA; ASH1L; NSD1; SMYD2; SETDB1; ATF7IP; SUV39H1; SUV39H2; SETDB2* |
|  | RMTs methylate histone arginine | *H3C15; H3C1; SMARCA4; WDR5; RBBP7; H4C1; PRMT1; H2AC18; H2AC20; H2AC21; H2AC14; H2AC12; H2AC4; H2AC6; H2AC1; H2AC11; H2AC7; H2AW; COPRS; CDK4; CCND1; PRMT5; WDR77; H2AB1; H2AZ2; H2AZ1; H2AX; H2AJ; PRMT7; SMARCB1; SMARCC2; SMARCC1; SMARCA2; ARID2; SMARCD1; PBRM1; SMARCD2; SMARCD3; ARID1B; ACTL6A; ARID1A; SMARCE1; ACTL6B; DNMT3A; PRMT6; JAK2; CARM1; PRMT3; RPS2* |
| Cellular  response to  hypoxia | Cellular response to hypoxia | *CUL2; RBX1; ELOC; ELOB; VHL; EGLN1; EGLN3; WTIP; AJUBA; LIMD1; EPAS1; HIF1A; HIF3A; UBE2D3; UBE2D2; UBE2D1; UBC; UBB; UBA52; RPS27A; EGLN2; PSMD3; PSMA6; PSMD2; PSMA5; PSMD13; PSMA4; PSMD12; PSMA3; PSMD11; PSMA2; SEM1; PSMD10; PSMA1; PSMD1; PSMD14; PSMC6; PSMC5; PSMC4; PSMC3; PSMC2; PSMC1; PSMB9; PSMB8; PSMB11; PSME4; PSMB7; PSMB6; PSMA8; PSMF1; PSME3; PSMB5; PSME2; PSMB4; PSME1; PSMD9; PSMB3; PSMD8; PSMB2; PSMD7; PSMD6; PSMB10; PSMD5; PSMB1; PSMD4; PSMA7; HIF1AN; HIGD1A; ARNT; EP300; CREBBP; CITED2; CA9; EPO; VEGFA; HIGD1A; CA9; EPO; VEGFA* |
| Detoxification  of ROS | Detoxification of Reactive Oxygen Species | *CAT; GPX1; SOD3; SOD1; CCS; NCF4; NCF1; NCF2; CYBA; CYBB; GSR; TXNRD1; TXN; GPX2; PRDX5; PRDX1; PRDX2; GPX6; GPX5; GSTP1; PRDX6; GPX3; NOX4; NOX5; PRDX5; TXN2; CYCS; TXNRD2; ATP7A; ATOX1; NUDT2; SOD2; AQP8; GSR; ERO1A; GPX8; GPX7; P4HB; PRDX3* |
| Cristae  formation | Cristae formation | *MICOS13; TMEM11; HSPA9; SAMM50; MTX1; MTX2; APOOL; CHCHD3; CHCHD6; MICOS10; IMMT; DNAJC11; APOO; ATP5PO; ATP5MC3; MT-ATP6; ATP5ME; ATP5MF; MT-ATP8; ATP5MC1; ATP5MG; ATP5PF; ATP5PD; ATP5PB; ATP5MC2; ATP5F1B; ATP5F1C; ATP5F1D; ATP5F1E; ATP5F1A; DMAC2L* |

**Supplementary table 3.** Overview of up and down differentially expressed pathways (DEP) from ROAST analysis of *in vivo* (bovine and mouse) and *in vitro* (bovine, mouse and human) pre-implantation development stages (one-sided directional p-value < 0.05). Up: number of up-regulated pathways; Down: number of down regulated pathways; NS: number of non-significant pathways.

| **Species** | **Collection** | **Stage** | **Up** | **Down** | **NS** | **DEP (%)** | **Average**  **DEP (%)** |
| --- | --- | --- | --- | --- | --- | --- | --- |
| **Human** | *in vitro* | *2C-MII* | 24 | 21 | 71 | 38.79 | 55.17 |
|  |  | *4C-2C* | 2 | 8 | 106 | 8.62 |  |
|  |  | *8C-4C* | 4 | 87 | 25 | 78.45 |  |
|  |  | *MO-8C* | 30 | 68 | 18 | 84.48 |  |
|  |  | *BL-MO* | 67 | 9 | 40 | 65.52 |  |
| **Bovine** | *in vivo* | *2C-MII* | 15 | 2 | 99 | 14.66 | 34.07 |
|  |  | *4C-2C* | 0 | 1 | 115 | 0.86 |  |
|  |  | *8C-4C* | 64 | 29 | 23 | 80.17 |  |
|  |  | *16C-8C* | 2 | 2 | 112 | 3.45 |  |
|  |  | *BL-16C* | 58 | 24 | 34 | 70.69 |  |
|  | *in vitro* | *2C-MII* | 66 | 40 | 10 | 91.38 | 83.45 |
|  |  | *4C-2C* | 29 | 80 | 7 | 93.97 |  |
|  |  | *8C-4C* | 35 | 42 | 39 | 66.38 |  |
|  |  | *16C-8C* | 60 | 20 | 36 | 68.97 |  |
|  |  | *BL-16C* | 64 | 48 | 4 | 96.55 |  |
| **Mouse** | *in vivo* | *2C-MII* | 74 | 32 | 10 | 91.38 | 82.06 |
|  |  | *4C-2C* | 67 | 36 | 13 | 88.79 |  |
|  |  | *8C-4C* | 43 | 35 | 38 | 67.24 |  |
|  |  | *MO-8C* | 64 | 12 | 40 | 65.52 |  |
|  |  | *BL-MO* | 79 | 34 | 3 | 97.41 |  |
|  | *in vitro* | *4C-2C* | 67 | 46 | 3 | 97.41 | 81.89 |
|  |  | *8C-4C* | 42 | 43 | 31 | 73.28 |  |
|  |  | *MO-8C* | 49 | 18 | 49 | 57.76 |  |
|  |  | *BL-MO* | 86 | 29 | 1 | 99.14 |  |

**Supplementary table 4.** Overview of up and down differentially expressed genes (DEG) from Limma analysis of *in vivo* *versus* *in vitro* bovine and mouse pre-implantation development stages (adjusted p-value < 0.05). Up: number of up-regulated genes; Down: number of down regulated genes.

| **Species** | **Stage** | **Up** | **Down** | **DEG %** | **Average DEG (%)** |
| --- | --- | --- | --- | --- | --- |
| **Bovine** | *MII* | 3317 | 1104 | 5.61 | 4.47 |
|  | *2C* | 1557 | 1480 | 3.85 |  |
|  | *4C* | 1720 | 1021 | 3.48 |  |
|  | *8C* | 1786 | 1011 | 3.55 |  |
|  | *16C* | 2450 | 1136 | 4.55 |  |
|  | *BL* | 2879 | 1680 | 5.79 |  |
| **Mouse** | *2C* | 2258 | 5467 | 11.77 | 9.58 |
|  | *4C* | 1885 | 5126 | 10.68 |  |
|  | *8C* | 54 | 98 | 0.23 |  |
|  | *MO* | 1328 | 6934 | 12.58 |  |
|  | *BL* | 4126 | 4177 | 12.65 |  |

**Supplementary table 5.** Overview of up and down differentially expressed pathways (DEP) from ROAST analysis comparing *in vivo* and *in vitro* bovine and mouse pre-implantation development stages (one-sided directional p-value < 0.05). Up: number of up-regulated pathways; Down: number of down regulated pathways; NS: number of non-significant pathways.

| **Species** | **Stage** | **Up** | **Down** | **NS** | **DEP (%)** | **Average DEP (%)** |
| --- | --- | --- | --- | --- | --- | --- |
| **Bovine** | *MII* | 51 | 51 | 14 | 87.93 | 85.35 |
|  | *2C* | 37 | 49 | 30 | 74.14 |  |
|  | *4C* | 75 | 22 | 19 | 83.62 |  |
|  | *8C* | 67 | 29 | 20 | 82.76 |  |
|  | *16C* | 66 | 38 | 12 | 89.66 |  |
|  | *BL* | 77 | 32 | 7 | 93.97 |  |
| **Mouse** | *2C* | 48 | 63 | 5 | 96.52 | 85.22 |
|  | *4C* | 58 | 51 | 7 | 94.78 |  |
|  | *8C* | 18 | 35 | 63 | 46.09 |  |
|  | *MO* | 64 | 43 | 9 | 93.04 |  |
|  | *BL* | 51 | 59 | 6 | 95.65 |  |

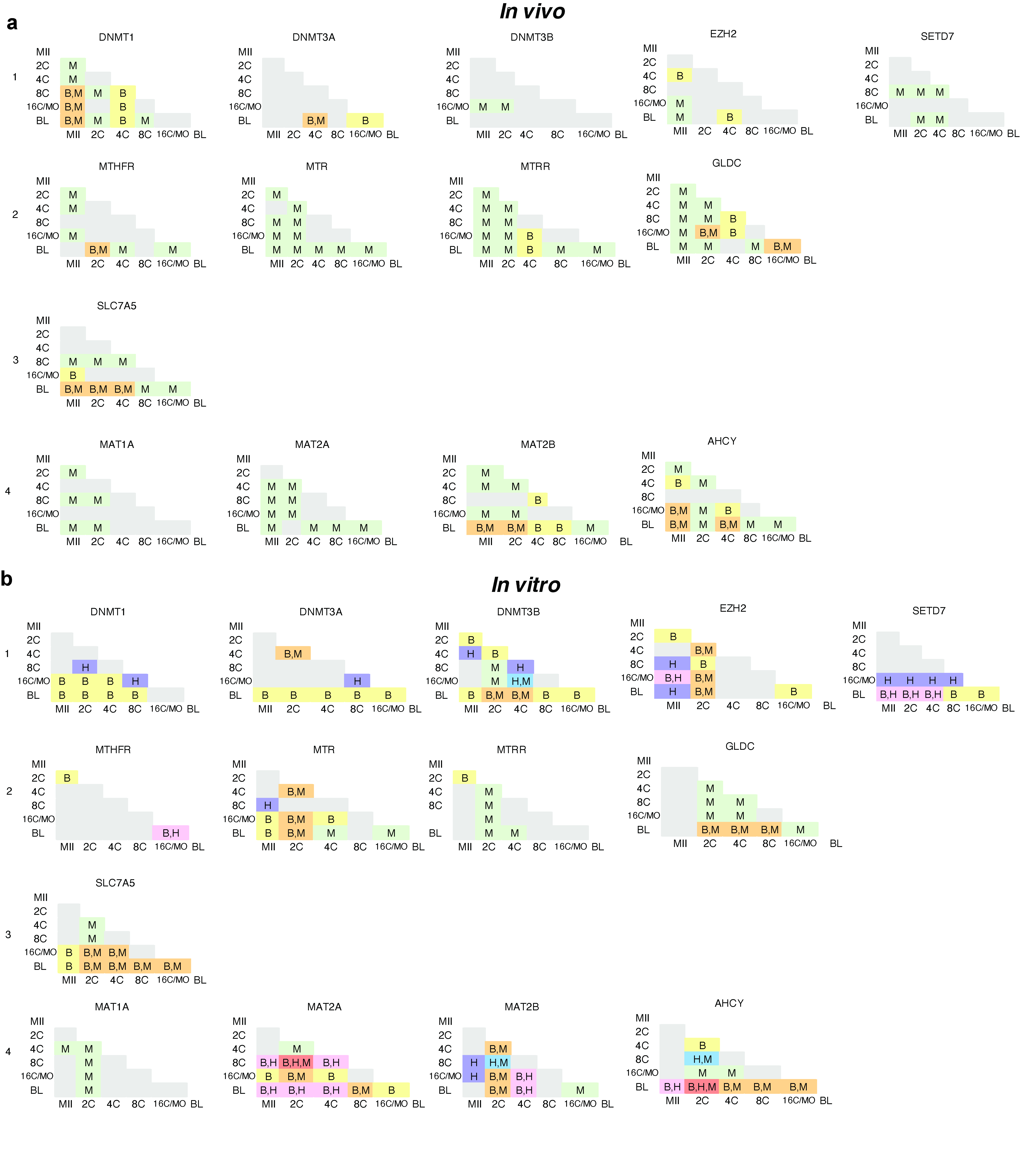
**Supplementary Figure 1.** Significance of up and down regulated genes presented in Figure 5 comparing different stages for each species: human (H), bovine (B) and mouse (M) *in vivo* (a) and *in vitro* (b). Boxes marked with the species letter (H, B and/or M) indicates significant difference between stages for the corresponding specie(s) (Limma, adjusted p-value <0.05)

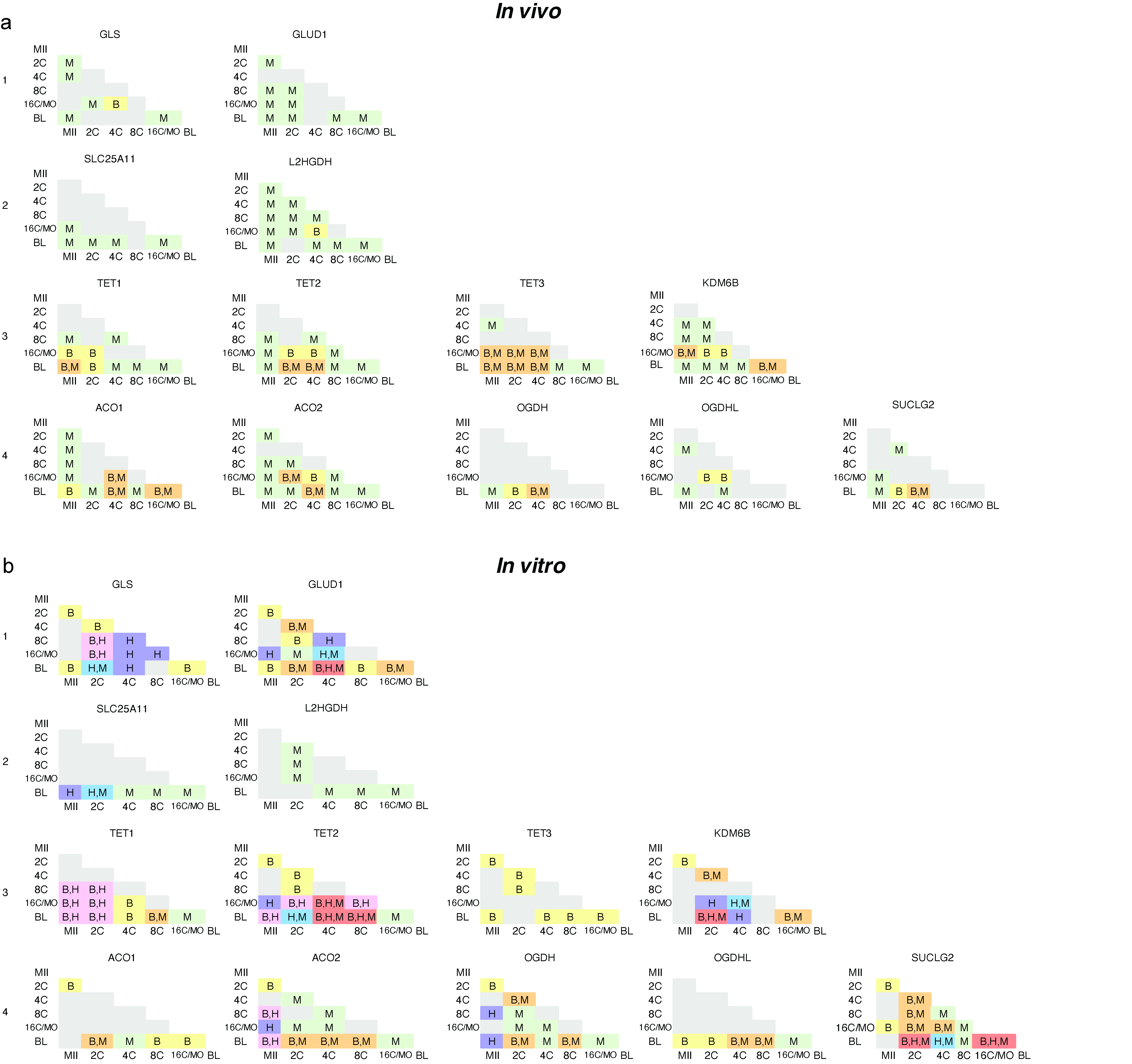
**Supplementary Figure 2.** Significance of up and down regulated genes presented in Figure 6 comparing different stages for each species: human (H), bovine (B) and mouse (M) *in vivo* (a) and *in vitro* (b). Boxes marked with the species letter (H, B and/or M) indicates significant difference between stages for the corresponding specie(s) (Limma, adjusted p-value <0.05)

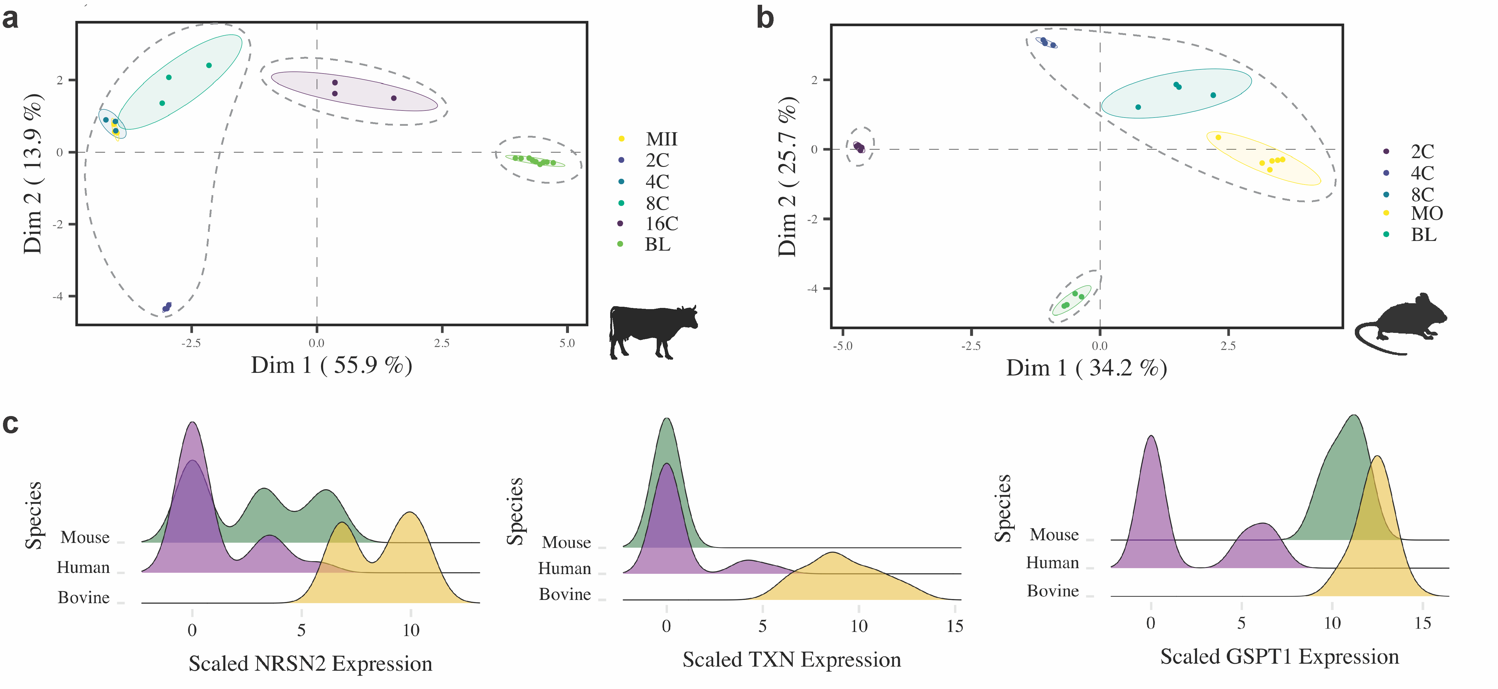

**Supplementary Figure 3.** General analysis of *in vitro* bovine and mouse mature oocyte (MII) and embryos (2C, 4C, 8C, 16C, MO and BL) gene expression. PCA of bovine (**a**) and mouse (**b**) *in vitro* developmental stages. In **c**, density plots of the genes of primary importance for classifying inter-species variation.

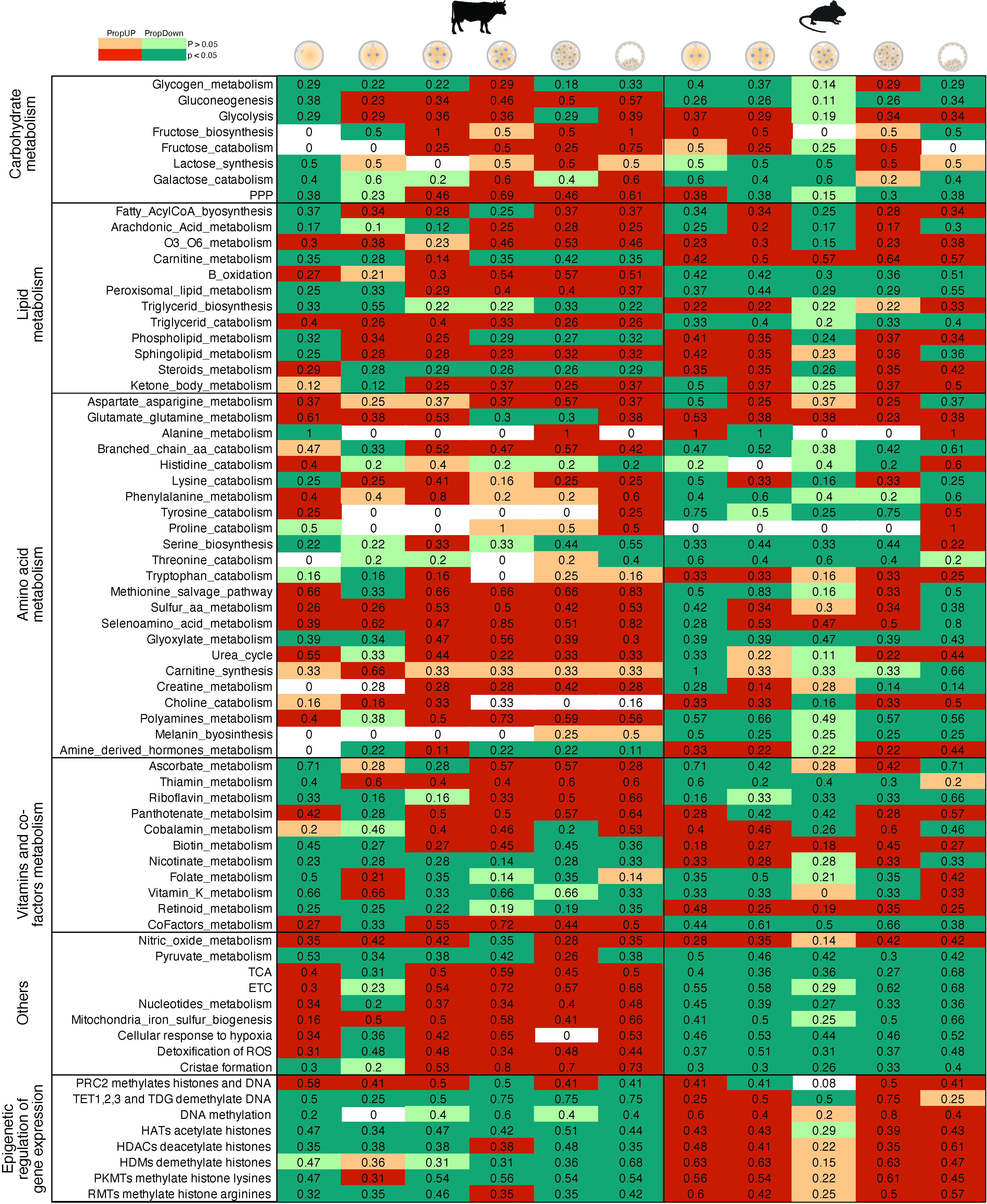

**Supplementary Figure 4.** General analysis mouse and bovine *in vito* mature oocyte (MII) and embryos (2C, 4C, 8C, 16C, MO and BL) gene expression of metabolism and epigenetic genes, showing proportion of up (PropUp, red) and down (PropDown, green) regulated metabolic pathways (part of Reactome terms “Metabolism” and “Epigenetic regulation of gene expression”) in 2C compared to MII, 4C compared to 2C, 8C compared to 4C, 16C compared to 8C and BL compared to 16C. Differences on PropUp and PropDown were calculated using the rotation gene set testing (ROAST), significant up and down regulated pathways presented the one-sided directional p-value < 0.05..

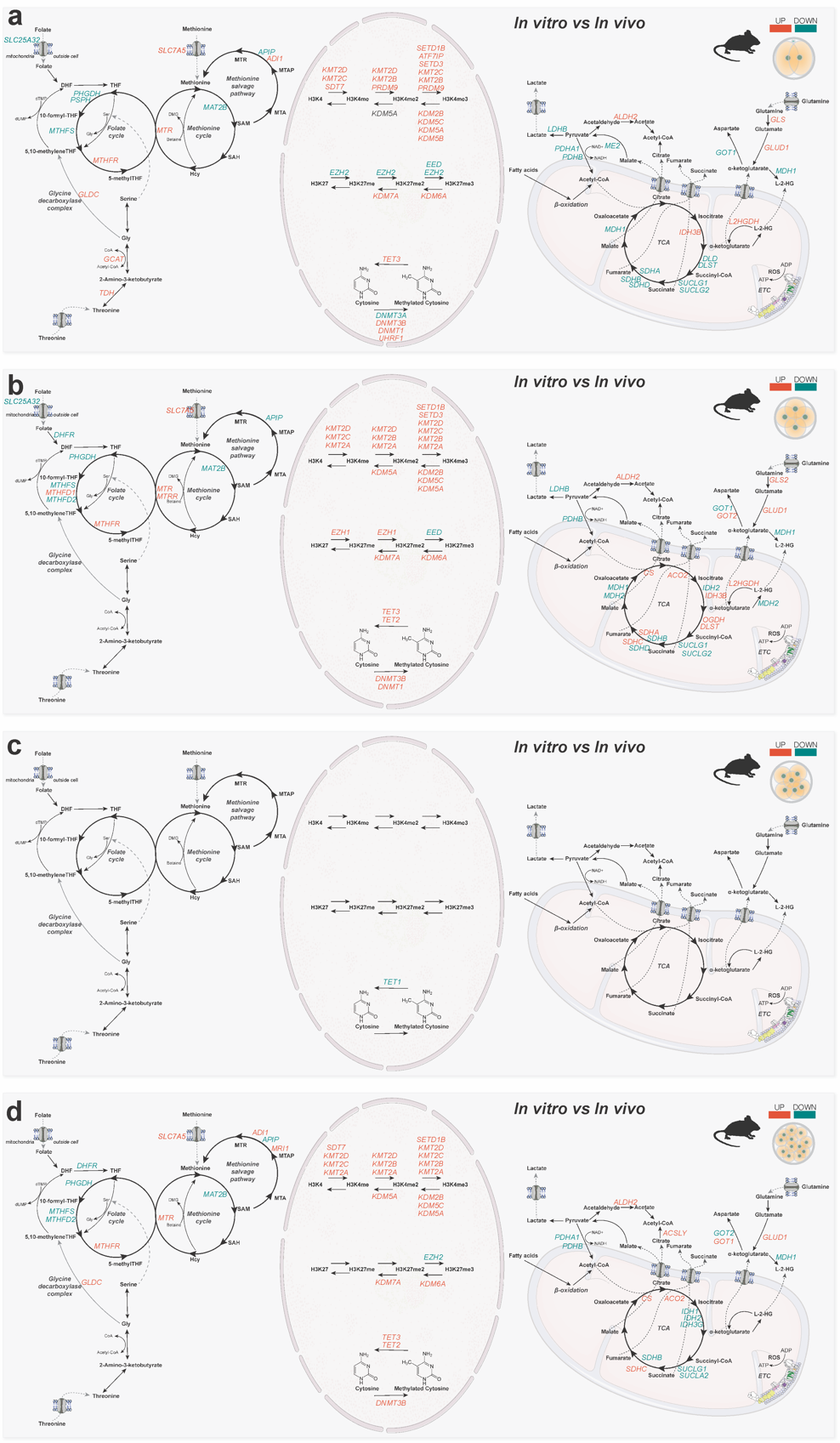

**Supplementary Figure 5.** Metaboloepigenetic genes up (red) and down (green) regulated (adjusted p-value < 0.05) comparing *in vitro vs in vivo* mouse MII (**a**), 2C (**b**), 4C (**c**), 8C (**d**) and 16C (**e**).

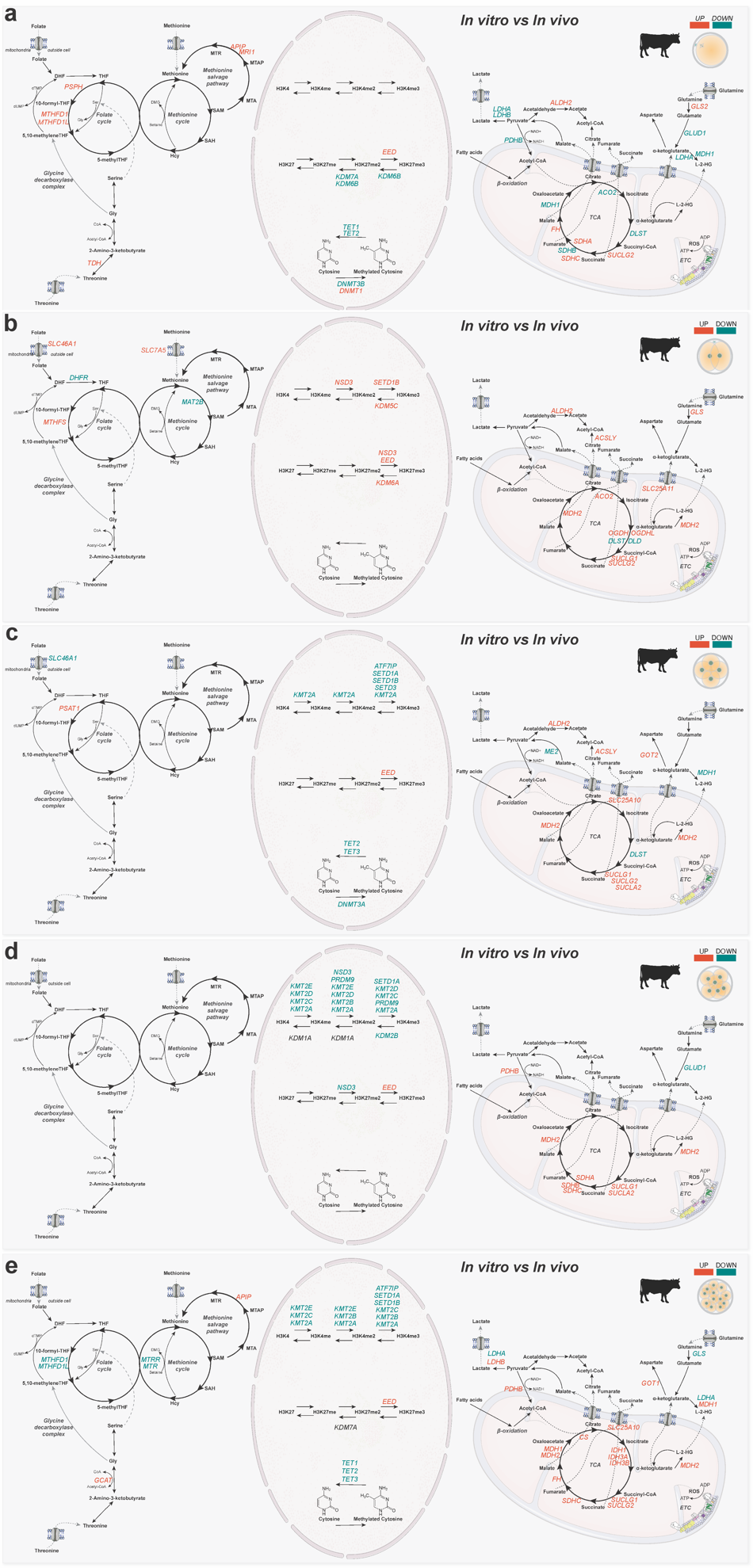

**Supplementary Figure 6.** Metaboloepigenetic genes up (red) and down (green) regulated (adjusted p-value < 0.05) comparing *in vitro vs in vivo* bovine MII (**a**), 2C (**b**), 4C (**c**), 8C (**d**) and 16C (**e**).
