## Supplementary file S2 for "RNAseq analysis reveals dynamic metaboloepigenetic profiles of human, mouse and bovine pre-implantation embryos": Supplementary file S2.html

Supplementary file S1 from: RNAseq analysis reveals different metaboloepigenetic reprogramming profile of human, mouse and bovine embryos


### Supplementary file S1 from: RNAseq analysis reveals different metaboloepigenetic reprogramming profile of human, mouse and bovine embryos

###### Marcella Pecora Milazzotto, Michael James Noonan, Marcia de Almeida Monteiro Melo Ferraz

#### 07/06/2021

---

#### Data scaling

The first step was to transform the raw data so that they would be on the same scale. To render the data comparable across species and developmental stages, EdgeR transformed data were scaled using Probabilistic Quotient Normalisation (PQN; Dieterle et al. 2006), which calibrates individual gene expression profiles against the median profile. Notably, analyses on PQN transformed data have been shown to have low false-positive rates, and can accurately recover groups of interest without introducing artefactual differences (Noonan et al. 2018).

```
#Load in the packages
library(FactoMineR)
library(ggplot2)
library(ellipse)
library(randomForest)
library(caret)
library(ggridges)
library(viridis)
library(gridExtra)
library(limma)

#Load in the raw data
data <- read.csv("All_Merged_Selected.csv")
data <- data[1:13134,]
SPECIES <- data[1,-1]
GROUP <- data[2,-1]
NAMES <- data[-c(1:2),1]
data <- data.frame(t(data[-c(1:2),-1]))

data <- as.data.frame(do.call(cbind,lapply(1:ncol(data), function(t)
  data[,t] <- as.numeric(data[,t])
)))

colnames(data) <- NAMES

#Plot the data to visualise the raw data
for(i in 1:nrow(data)){
  if(i == 1){plot(density(as.matrix(data[i,])),
                  ylim = c(0,0.8),
                  col = viridis::viridis(nrow(data))[i],
                  main = "",
                  xlab = "Raw expression values")
  } else {
    lines(density(as.matrix(data[i,])),
          ylim = c(0,0.7),
          col = viridis::viridis(nrow(data))[i])
  }
}
```

```
# PQN Normalisation of the data
#Calculate the median of each gene's expression to generate a reference sample
ref <- apply(data, 2, median)

for(i in 1:nrow(data)){
  QUOTIENTS <- data[i,]/ref
  m_j <- median(t(QUOTIENTS))
  data[i,] <- data[i,]/m_j
  data[i,][data[i,] == min(data[i,])] <- 0
}

#Plot the data to visualise the normalised data
for(i in 1:nrow(data)){
  if(i == 1){plot(density(as.matrix(data[i,])),
                  ylim = c(0,0.8),
                  col = viridis::viridis(nrow(data))[i],
                  main = "",
                  xlab = "Scaled expression")
  } else {
    lines(density(as.matrix(data[i,])),
          ylim = c(0,0.7),
          col = viridis::viridis(nrow(data))[i])
  }
}
```

---

#### PCA on all data

First step of the analyses was to run a PCA on all of the data to predict species and plot the results to visualise any clustering that might be occurring.

```
#Load in the meta data
IDs <- na.omit(read.csv("all_extra.csv"))

res.pca <- PCA(data, graph = FALSE) # Conduct a PCA on the data
PC1 <- res.pca$ind$coord[,1] #Store individual coordinates of PC1 as a vector
PC2 <- res.pca$ind$coord[,2] #Store individual coordinates of PC2 as a vector
PCs.ID <- data.frame(cbind(PC1,PC2)) #Bind the coordinates together as a dataframe
PCs.ID$Species <- (IDs$Species) #Add in Species to the df

#Define axis labels based on % of data explained across each dimension of the PCA
DIM_1 <- paste("PCA Dimension 1 (", round(res.pca$eig[1,2], 1), "%)")
DIM_2 <- paste("PCA Dimmension 2  (", round(res.pca$eig[2,2], 1), "%)")

#Draw ellipses around the cluster
centroids <- aggregate(cbind(PC1,PC2)~Species,PCs.ID,mean)
conf.rgn  <- do.call(rbind,lapply(unique(PCs.ID$Species),function(t)
  data.frame(Species=as.character(t),
             ellipse(cov(PCs.ID[PCs.ID$Species==t,1:2]),
                     centre=as.matrix(centroids[which(unique(PCs.ID$Species)==t),2:3]),
                     level=0.95),
             stringsAsFactors=FALSE)))

#Then make the figure
ggplot(PCs.ID, aes(x=PC1, y=PC2, color = Species), guide = FALSE) +
  geom_path(data=conf.rgn, alpha=0.2, size = 0, show.legend = FALSE) +
  geom_polygon(data=conf.rgn,
               aes(fill = Species),
               alpha=0.1, size = 0.1,
               show.legend = FALSE) + 
  geom_hline(aes(yintercept=0), linetype="dashed", lwd = 0.1) +
  geom_vline(aes(xintercept=0), linetype="dashed", lwd = 0.1) +
  theme_bw() +
  geom_point(size=0.4, aes(color = Species)) +
  scale_color_manual(labels=c("Bovine", "Human", "Mouse"),
                     values = c("#e6c141", "#8a3bb8", "#3c7a47")) +
  scale_fill_manual(labels=c("Bovine", "Human", "Mouse"),
                    values = c("#e6c141", "#8a3bb8", "#3c7a47"),
                    guide = FALSE) +
  ylab(DIM_2) +
  xlab(DIM_1) + 
  theme(panel.grid.major = element_blank(),
        panel.border = element_rect(colour = "black", size=1),
        panel.grid.minor = element_blank(),
        axis.title.x  = element_text(size=10, family = "serif"),
        axis.title.y  = element_text(size=10, family = "serif"),
        plot.title = element_text(size=8, hjust = 0, family = "serif"),
        axis.text.y  = element_text(size=5, family = "serif"),
        axis.text.x  = element_text(size=5, family = "serif"),
        legend.position=c(0.15,0.9),
        legend.background = element_blank(),
        legend.title = element_blank(),
        legend.text = element_text(size=8, family = "serif"),
        legend.key.size = unit(0.3, "cm"),
        legend.key = element_blank())
```

```
ggsave(file="Figures/Species_PCAF.png",
       width = 3.23,
       height=3,
       units = "in",
       dpi = 600)
```

---

#### Random Forest Classification

After this initial data visualisation, a random forest (RF) model (Ho 1995) was used to classify intra- and inter-species gene expression profiles according to developmental stages and collection method (in vivo or in vitro), with scaled gene expression values as the prediction variables. This allowed us to determine how well information contained within the expression data could be used to predict classes of interest. These analyses were conducted using the R package randomForest (RColorBrewer and Liaw, 2018). We chose RF modeling as it does not require any parameter reduction prior to analysis (Cutler et al. 2007), and has been shown to provide reliable results for biomarker identification (Chen et al. 2017; Noonan et al. 2018). Identification of genes important for classifying groups of interest in each RF model was carried out using RF variable importance values.

##### Classify species with in vivo data

Our first RF model aimed to classify species based on the data collected *in vivo*. Because data from humans were not collected *in vivo*, the *in vitro* data for humans were used here for comparison.

```
#Create a dataset that contains only the in vivo data (except for humans)
Species_Data <- data
Species_Data$Species <- as.factor(IDs$Species)
names(Species_Data) <- make.names(names(Species_Data))
Species_Data <- Species_Data[-which(IDs$Collection == "vitro" & IDs$Species == "mouse"),]
Species_Data <- Species_Data[-which(IDs$Collection == "vitro" & IDs$Species == "cow"),]

#Run the random forest model identifying species
Species.mod <- randomForest(y = Species_Data$Species,
                            x = Species_Data[, colnames(Species_Data) != "Species"],
                            mtry =  5,
                            ntree= 20000,
                            importance=TRUE,
                            proximity = TRUE,
                            keep.forest=TRUE,
                            replace = TRUE)

Species.mod
```

```
## 
## Call:
##  randomForest(x = Species_Data[, colnames(Species_Data) != "Species"],      y = Species_Data$Species, ntree = 20000, mtry = 5, replace = TRUE,      importance = TRUE, proximity = TRUE, keep.forest = TRUE) 
##                Type of random forest: classification
##                      Number of trees: 20000
## No. of variables tried at each split: 5
## 
##         OOB estimate of  error rate: 0%
## Confusion matrix:
##       cow human mouse class.error
## cow    12     0     0           0
## human   0    20     0           0
## mouse   0     0    17           0
```

```
varImpPlot(Species.mod, type=1, scale = FALSE)
```

```
#Get top 20 genes
PRIME_GENES <- row.names(Species.mod$importance)[order(Species.mod$importance[,"MeanDecreaseAccuracy"], decreasing = TRUE)][1:20]
PRIME_GENES
```

```
##  [1] "SLITRK2" "SMG1"    "SLITRK5" "ZNF597"  "CSNK1G3" "CCDC6"   "CALM1"  
##  [8] "BCL10"   "LYPD4"   "SPIDR"   "HCFC1"   "CRNKL1"  "JMJD6"   "ZNF330" 
## [15] "CENPT"   "IST1"    "FAM83D"  "FOXRED2" "HYPK"    "SLC35A3"
```

```
res.pca <- PCA(Species.mod$proximity, graph = FALSE) # Conduct a PCA on the proximity matrix
PC1 <- res.pca$ind$coord[,1] #Store individual coordinates of PC1 as a vector
PC2 <- res.pca$ind$coord[,2] #Store individual coordinates of PC2 as a vector
PCs.ID <- data.frame(cbind(PC1,PC2)) #Bind the coordinates together as a dataframe
PCs.ID$Subject <- Species_Data$Species #Add in species to the df

#Define axis labels based on % of data explained across each dimension of the PCA
DIM_1 <- paste("Dim 1 (", round(res.pca$eig[1,2], 1), "%)")
DIM_2 <- paste("Dim 2 (", round(res.pca$eig[2,2], 1), "%)")

#Draw ellipses around the clusters and highlighting the elipsoids rather than the data points
centroids <- aggregate(cbind(PC1,PC2)~Subject,PCs.ID,mean)
conf.rgn  <- do.call(rbind,lapply(unique(PCs.ID$Subject),function(t)
  data.frame(Subject=as.character(t),
             ellipse(cov(PCs.ID[PCs.ID$Subject==t,1:2]),
                     centre=as.matrix(centroids[t,2:3]),
                     level=0.95),
             stringsAsFactors=FALSE)))

#Then make the figure
PCA_FIG_Species <- 
  ggplot(PCs.ID, aes(x=PC1, y=PC2, color = Subject)) +
  geom_hline(aes(yintercept=0), linetype="dashed", lwd = 0.1) +
  geom_vline(aes(xintercept=0), linetype="dashed", lwd = 0.1) +
  geom_path(data=conf.rgn, alpha=0.2, size = 0) +
  geom_polygon(data=conf.rgn,
               aes(fill = Subject),
               alpha=0.1,
               size = 0.1,
               show.legend = FALSE) + 
  geom_point(size=0.4) +
  theme_bw() +
  ylab(DIM_2) +
  xlab(DIM_1) + 
  geom_point(size=0.4, aes(color = Subject)) +
  scale_color_manual(labels=c("Bovine", "Human", "Mouse"),
                     values = c("#e6c141", "#8a3bb8", "#3c7a47")) +
  scale_fill_manual(labels=c("Bovine", "Human", "Mouse"),
                    values = c("#e6c141", "#8a3bb8", "#3c7a47"),
                    guide = FALSE) +
  ylab(DIM_2) +
  xlab(DIM_1) + 
  theme(panel.grid.major = element_blank(),
        panel.border = element_rect(colour = "black", size=1),
        panel.grid.minor = element_blank(),
        axis.title.x  = element_text(size=10, family = "serif"),
        axis.title.y  = element_text(size=10, family = "serif"),
        plot.title = element_text(size=8, hjust = 0, family = "serif"),
        axis.text.y  = element_text(size=5, family = "serif"),
        axis.text.x  = element_text(size=5, family = "serif"),
        legend.position=c(0.8,0.15),
        legend.background = element_blank(),
        legend.title = element_blank(),
        legend.text = element_text(size=8, family = "serif"),
        legend.key.size = unit(0.3, "cm"),
        legend.key = element_blank())

ggsave(PCA_FIG_Species,
       file="Figures/Species_PCA_RF_Vivo.png",
       width = 3.23,
       height=3,
       units = "in",
       dpi = 600)

#Figure of the primary genes
X_LAB <- paste("Scaled", PRIME_GENES[1], "Expression")

a <- 
  ggplot(Species_Data, aes(x=Species_Data[,PRIME_GENES[1]], y=Species, fill = Species)) +
  geom_density_ridges(scale = 5, alpha=0.6, size = 0.2) +
  theme_ridges() +
  scale_fill_manual(labels=c("Bovine", "Human", "Mouse"),
                    values = c("#e6c141", "#8a3bb8", "#3c7a47")) +
  scale_y_discrete(expand = c(0.1, 0), labels=c("Bovine", "Human", "Mouse")) +
  labs(x=X_LAB, y="Species")+
  ggtitle("a)")+
  theme(plot.title = element_text(hjust = 0, size = 10, family = "serif"),
        panel.grid.major = element_blank(),
        panel.border = element_rect(colour = "black", size=1),
        panel.grid.minor = element_blank(),
        axis.title.x  = element_text(hjust = 0.5, size=10, family = "serif"),
        axis.title.y  = element_text(hjust = 0.5, size=10, family = "serif"),
        axis.text.y  = element_text(size=8, family = "serif"),
        axis.text.x  = element_text(size=8, family = "serif"),
        legend.position=c("none"))

X_LAB <- paste("Scaled", PRIME_GENES[2], "Expression")
b <- 
  ggplot(Species_Data, aes(x=Species_Data[,PRIME_GENES[2]], y=Species, fill = Species)) +
  geom_density_ridges(scale = 5, alpha=0.6, size = 0.2) +
  theme_ridges() +
  scale_fill_manual(labels=c("Bovine", "Human", "Mouse"),
                    values = c("#e6c141", "#8a3bb8", "#3c7a47")) +
  scale_y_discrete(expand = c(0.1, 0), labels=c("Bovine", "Human", "Mouse")) +
  labs(x=X_LAB, y="Species")+
  ggtitle("b)")+
  theme(plot.title = element_text(hjust = 0, size = 10, family = "serif"),
        panel.grid.major = element_blank(),
        panel.border = element_rect(colour = "black", size=1),
        panel.grid.minor = element_blank(),
        axis.title.x  = element_text(hjust = 0.5, size=10, family = "serif"),
        axis.title.y  = element_text(hjust = 0.5, size=10, family = "serif"),
        axis.text.y  = element_text(size=8, family = "serif"),
        axis.text.x  = element_text(size=8, family = "serif"),
        legend.position=c("none"))

X_LAB <- paste("Scaled", PRIME_GENES[3], "Expression")
c <- 
  ggplot(Species_Data, aes(x=Species_Data[,PRIME_GENES[3]], y=Species, fill = Species))+ #EXOSC1
  geom_density_ridges(scale = 5, alpha=0.6, size = 0.2) +
  theme_ridges() +
  scale_fill_manual(labels=c("Bovine", "Human", "Mouse"),
                    values = c("#e6c141", "#8a3bb8", "#3c7a47")) +
  scale_y_discrete(expand = c(0.1, 0), labels=c("Bovine", "Human", "Mouse")) +
  labs(x=X_LAB, y="Species")+
  ggtitle("c)")+
  theme(plot.title = element_text(hjust = 0, size = 10, family = "serif"),
        panel.grid.major = element_blank(),
        panel.border = element_rect(colour = "black", size=1),
        panel.grid.minor = element_blank(),
        axis.title.x  = element_text(hjust = 0.5, size=10, family = "serif"),
        axis.title.y  = element_text(hjust = 0.5, size=10, family = "serif"),
        axis.text.y  = element_text(size=8, family = "serif"),
        axis.text.x  = element_text(size=8, family = "serif"),
        legend.position=c("none"))

FIG <- arrangeGrob(a, b, c, ncol = 1)

grid.arrange(a, b, c, ncol = 1)
```

```
ggsave(FIG,
       file="Figures/Species_Density_Plot_Vivo.png",
       width = 3.23,
       height=6,
       units = "in",
       dpi = 600)
```

##### Classify species with in vitro data

Our next RF model aimed to classify species based on the data collected *in vitro*.

```
#Create a dataset that contains on the in vitro data
Species_Data <- data
Species_Data$Species <- as.factor(IDs$Species)
names(Species_Data) <- make.names(names(Species_Data))
Species_Data <- Species_Data[-which(IDs$Collection == "vivo" & IDs$Species == "mouse"),]
Species_Data <- Species_Data[-which(IDs$Collection == "vivo" & IDs$Species == "cow"),]

#Run the random forest model identifying species
Species.mod <- randomForest(y = Species_Data$Species, 
                            x = Species_Data[, colnames(Species_Data) != "Species"],
                            mtry =  5,
                            ntree= 20000,
                            importance=TRUE,
                            proximity = TRUE,
                            keep.forest=TRUE,
                            replace = TRUE)
Species.mod
```

```
## 
## Call:
##  randomForest(x = Species_Data[, colnames(Species_Data) != "Species"],      y = Species_Data$Species, ntree = 20000, mtry = 5, replace = TRUE,      importance = TRUE, proximity = TRUE, keep.forest = TRUE) 
##                Type of random forest: classification
##                      Number of trees: 20000
## No. of variables tried at each split: 5
## 
##         OOB estimate of  error rate: 0%
## Confusion matrix:
##       cow human mouse class.error
## cow    24     0     0           0
## human   0    20     0           0
## mouse   0     0    22           0
```

```
varImpPlot(Species.mod, type=1, scale = FALSE)
```

```
#Get top 20 genes
PRIME_GENES <- row.names(Species.mod$importance)[order(Species.mod$importance[,"MeanDecreaseAccuracy"], decreasing = TRUE)][1:20]
PRIME_GENES
```

```
##  [1] "NRSN2"   "TXN"     "GSPT1"   "YWHAH"   "KLHL18"  "REV3L"   "SRPK1"  
##  [8] "HIC2"    "ACADM"   "HHIP"    "BRPF3"   "ARMCX3"  "SMG7"    "RAC1"   
## [15] "GPR161"  "SLC25A3" "SELENOK" "ATP5IF1" "TDRD3"   "RALGAPB"
```

```
res.pca <- PCA(Species.mod$proximity, graph = FALSE) # Conduct a PCA on the proximity matrix
PC1 <- res.pca$ind$coord[,1] #Store individual coordinates of PC1 as a vector
PC2 <- res.pca$ind$coord[,2] #Store individual coordinates of PC2 as a vector
PCs.ID <- data.frame(cbind(PC1,PC2)) #Bind the coordinates together as a dataframe
PCs.ID$Subject <- Species_Data$Species #Add in species to the df

#Define axis labels based on % of data explained across each dimension of the PCA
DIM_1 <- paste("Dim 1 (", round(res.pca$eig[1,2], 1), "%)")
DIM_2 <- paste("Dim 2 (", round(res.pca$eig[2,2], 1), "%)")

#Draw ellipses around the clusters and highlighting the elipsoids rather than the data points
centroids <- aggregate(cbind(PC1,PC2)~Subject,PCs.ID,mean)
conf.rgn  <- do.call(rbind,lapply(unique(PCs.ID$Subject),function(t)
  data.frame(Subject=as.character(t),
             ellipse(cov(PCs.ID[PCs.ID$Subject==t,1:2]),
                     centre=as.matrix(centroids[t,2:3]),
                     level=0.95),
             stringsAsFactors=FALSE)))


#Then make the figure
ggplot(PCs.ID, aes(x=PC1, y=PC2, color = Subject)) +
  geom_hline(aes(yintercept=0), linetype="dashed", lwd = 0.1) +
  geom_vline(aes(xintercept=0), linetype="dashed", lwd = 0.1) +
  geom_path(data=conf.rgn, alpha=0.2, size = 0) +
  geom_polygon(data=conf.rgn,
               aes(fill = Subject),
               alpha=0.1,
               size = 0.1,
               show.legend = FALSE) + 
  geom_point(size=0.4) +
  theme_bw() +
  ylab(DIM_2) +
  xlab(DIM_1) + 
  geom_point(size=0.4, aes(color = Subject)) +
  scale_color_manual(labels=c("Bovine", "Human", "Mouse"),
                     values = c("#e6c141", "#8a3bb8", "#3c7a47")) +
  scale_fill_manual(labels=c("Bovine", "Human", "Mouse"),
                    values = c("#e6c141", "#8a3bb8", "#3c7a47"),
                    guide = FALSE) +
  ylab(DIM_2) +
  xlab(DIM_1) + 
  theme(panel.grid.major = element_blank(),
        panel.border = element_rect(colour = "black", size=1),
        panel.grid.minor = element_blank(),
        axis.title.x  = element_text(size=10, family = "serif"),
        axis.title.y  = element_text(size=10, family = "serif"),
        plot.title = element_text(size=8, hjust = 0, family = "serif"),
        axis.text.y  = element_text(size=5, family = "serif"),
        axis.text.x  = element_text(size=5, family = "serif"),
        legend.position=c(0.8,0.15),
        legend.background = element_blank(),
        legend.title = element_blank(),
        legend.text = element_text(size=8, family = "serif"),
        legend.key.size = unit(0.3, "cm"),
        legend.key = element_blank())
```

```
ggsave(file="Figures/Species_PCA_RF_Vitro.png",
       width = 3.23,
       height=3,
       units = "in",
       dpi = 600)

X_LAB <- paste("Scaled", PRIME_GENES[1], "Expression")
a <- 
  ggplot(Species_Data, aes(x=Species_Data[,PRIME_GENES[1]], y=Species, fill = Species))+ #MYL12A
  geom_density_ridges(scale = 5, alpha=0.6, size = 0.2) +
  theme_ridges() +
  scale_fill_manual(labels=c("Bovine", "Human", "Mouse"),
                    values = c("#e6c141", "#8a3bb8", "#3c7a47")) +
  scale_y_discrete(expand = c(0.1, 0), labels=c("Bovine", "Human", "Mouse")) +
  labs(x=X_LAB, y="Species")+
  ggtitle("a)")+
  theme(plot.title = element_text(hjust = 0, size = 10, family = "serif"),
        panel.grid.major = element_blank(),
        panel.border = element_rect(colour = "black", size=1),
        panel.grid.minor = element_blank(),
        axis.title.x  = element_text(hjust = 0.5, size=10, family = "serif"),
        axis.title.y  = element_text(hjust = 0.5, size=10, family = "serif"),
        axis.text.y  = element_text(size=8, family = "serif"),
        axis.text.x  = element_text(size=8, family = "serif"),
        legend.position=c("none"))

X_LAB <- paste("Scaled", PRIME_GENES[2], "Expression")
b <- 
  ggplot(Species_Data, aes(x=Species_Data[,PRIME_GENES[2]], y=Species, fill = Species))+ #MT.ND2
  geom_density_ridges(scale = 5, alpha=0.6, size = 0.2) +
  theme_ridges() +
  scale_fill_manual(labels=c("Bovine", "Human", "Mouse"),
                    values = c("#e6c141", "#8a3bb8", "#3c7a47")) +
  scale_y_discrete(expand = c(0.1, 0), labels=c("Bovine", "Human", "Mouse")) +
  labs(x=X_LAB, y="Species")+
  ggtitle("b)")+
  theme(plot.title = element_text(hjust = 0, size = 10, family = "serif"),
        panel.grid.major = element_blank(),
        panel.border = element_rect(colour = "black", size=1),
        panel.grid.minor = element_blank(),
        axis.title.x  = element_text(hjust = 0.5, size=10, family = "serif"),
        axis.title.y  = element_text(hjust = 0.5, size=10, family = "serif"),
        axis.text.y  = element_text(size=8, family = "serif"),
        axis.text.x  = element_text(size=8, family = "serif"),
        legend.position=c("none"))

X_LAB <- paste("Scaled", PRIME_GENES[3], "Expression")
c <- 
  ggplot(Species_Data, aes(x=Species_Data[,PRIME_GENES[3]], y=Species, fill = Species))+ #HNRPC
  geom_density_ridges(scale = 5, alpha=0.6, size = 0.2) +
  theme_ridges() +
  scale_fill_manual(labels=c("Bovine", "Human", "Mouse"),
                    values = c("#e6c141", "#8a3bb8", "#3c7a47")) +
  scale_y_discrete(expand = c(0.1, 0), labels=c("Bovine", "Human", "Mouse")) +
  labs(x=X_LAB, y="Species")+
  ggtitle("c)")+
  theme(plot.title = element_text(hjust = 0, size = 10, family = "serif"),
        panel.grid.major = element_blank(),
        panel.border = element_rect(colour = "black", size=1),
        panel.grid.minor = element_blank(),
        axis.title.x  = element_text(hjust = 0.5, size=10, family = "serif"),
        axis.title.y  = element_text(hjust = 0.5, size=10, family = "serif"),
        axis.text.y  = element_text(size=8, family = "serif"),
        axis.text.x  = element_text(size=8, family = "serif"),
        legend.position=c("none"))

FIG <- arrangeGrob(a, b, c, ncol = 1)

grid.arrange(a, b, c, ncol = 1)
```

```
ggsave(FIG,
       file="Figures/Species_Density_Plot_Vitro.png",
       width = 3.23,
       height=6,
       units = "in",
       dpi = 600)
```

##### Classify Collection type for each of the species

Our next RF model aimed to classify collection type (i.e., *in vitro* vs. *in vivo*) for data from mice and cows. Here data from each species were analysed separately. Because data from humans were not collected *in vivo*, they were excluded from these analyses.

```
collection_Data <- data
collection_Data$Collection <- as.factor(IDs$Collection)
names(collection_Data) <- make.names(names(collection_Data))

collection_Data_cows <- collection_Data[which(IDs$Species == "cow"),]
collection_Data_mice <- collection_Data[which(IDs$Species == "mouse"),]

#Run the random forest model identifying vitro/vivo for cows
Collection.mod.cows <- randomForest(y = collection_Data_cows$Collection,
                                    x = collection_Data_cows[, colnames(collection_Data_cows) != "Collection"],
                                    mtry =  14,
                                    ntree= 20000,
                                    importance=TRUE,
                                    proximity = TRUE,
                                    keep.forest=TRUE,
                                    replace = TRUE)
Collection.mod.cows
```

```
## 
## Call:
##  randomForest(x = collection_Data_cows[, colnames(collection_Data_cows) !=      "Collection"], y = collection_Data_cows$Collection, ntree = 20000,      mtry = 14, replace = TRUE, importance = TRUE, proximity = TRUE,      keep.forest = TRUE) 
##                Type of random forest: classification
##                      Number of trees: 20000
## No. of variables tried at each split: 14
## 
##         OOB estimate of  error rate: 0%
## Confusion matrix:
##       vitro vivo class.error
## vitro    24    0           0
## vivo      0   12           0
```

```
#Run the random forest model identifying vitro/vivo for mice
Collection.mod.mice <- randomForest(y = collection_Data_mice$Collection,
                                    x = collection_Data_mice[, colnames(collection_Data_mice) != "Collection"],
                                    mtry =  14,
                                    ntree= 20000,
                                    importance=TRUE,
                                    proximity = TRUE,
                                    keep.forest=TRUE,
                                    replace = TRUE)
Collection.mod.mice
```

```
## 
## Call:
##  randomForest(x = collection_Data_mice[, colnames(collection_Data_mice) !=      "Collection"], y = collection_Data_mice$Collection, ntree = 20000,      mtry = 14, replace = TRUE, importance = TRUE, proximity = TRUE,      keep.forest = TRUE) 
##                Type of random forest: classification
##                      Number of trees: 20000
## No. of variables tried at each split: 14
## 
##         OOB estimate of  error rate: 2.56%
## Confusion matrix:
##       vitro vivo class.error
## vitro    21    1  0.04545455
## vivo      0   17  0.00000000
```

```
####################################
# PCA on proximity matrix for cows

res.pca <- PCA(Collection.mod.cows$proximity, graph = FALSE) # Conduct a PCA on the proximity matrix
PC1 <- res.pca$ind$coord[,1] #Store individual coordinates of PC1 as a vector
PC2 <- res.pca$ind$coord[,2] #Store individual coordinates of PC2 as a vector
PCs.ID <- data.frame(cbind(PC1,PC2)) #Bind the coordinates together as a dataframe
PCs.ID$Subject <- collection_Data_cows$Collection #Add in collection type to the df

DIM_1 <- paste("Dim 1 (", round(res.pca$eig[1,2], 1), "%)")
DIM_2 <- paste("Dim 2 (", round(res.pca$eig[2,2], 1), "%)")

#Draw ellipses around the clusters and highlighting the elipsoids rather than the data points
centroids <- aggregate(cbind(PC1,PC2)~Subject,PCs.ID,mean)
conf.rgn  <- do.call(rbind,lapply(unique(PCs.ID$Subject),function(t)
  data.frame(Subject=as.character(t),
             ellipse(cov(PCs.ID[PCs.ID$Subject==t,1:2]),
                     centre=as.matrix(centroids[t,2:3]),
                     level=0.95),
             stringsAsFactors=FALSE)))

#Then make the figure
PCA_FIG_Cows <- 
  ggplot(PCs.ID, aes(x=PC1, y=PC2, color = Subject), guide = FALSE) +
  geom_hline(aes(yintercept=0), linetype="dashed", lwd = 0.1) +
  geom_vline(aes(xintercept=0), linetype="dashed", lwd = 0.1) +
  geom_path(data=conf.rgn, alpha=0.2, size = 0) +
  geom_polygon(data=conf.rgn, aes(fill = Subject), alpha=0.1, size = 0.1, show.legend = FALSE) + 
  geom_point(size=0.4) +
  theme_bw() +
  ggtitle("a) - Bovine") +
  scale_color_manual(labels=c("Vitro", "Vivo"), values = c("#046C9A", "red")) +
  scale_fill_manual(labels=c("Vitro", "Vivo"), values = c("#046C9A", "red"),guide = FALSE) +
  ylab(DIM_2) +
  xlab(DIM_1) + 
  theme(panel.grid.major = element_blank(),
        panel.border = element_rect(colour = "black", size=1),
        panel.grid.minor = element_blank(),
        axis.title.x  = element_text(size=10, family = "serif"),
        axis.title.y  = element_text(size=10, family = "serif"),
        plot.title = element_text(size=8, hjust = 0, family = "serif"),
        axis.text.y  = element_text(size=5, family = "serif"),
        axis.text.x  = element_text(size=5, family = "serif"),
        legend.position=c(0.8,0.15),
        legend.background = element_blank(),
        legend.title = element_blank(),
        legend.text = element_text(size=8, family = "serif"),
        legend.key.size = unit(0.3, "cm"),
        legend.key = element_blank())

####################################
# PCA on proximity matrix for mice

res.pca <- PCA(Collection.mod.mice$proximity, graph = FALSE) # Conduct a PCA on the proximity matrix
PC1 <- res.pca$ind$coord[,1] #Store individual coordinates of PC1 as a vector
PC2 <- res.pca$ind$coord[,2] #Store individual coordinates of PC2 as a vector
PCs.ID <- data.frame(cbind(PC1,PC2)) #Bind the coordinates together as a dataframe
PCs.ID$Subject <- collection_Data_mice$Collection #Add in Season to the df

DIM_1 <- paste("Dim 1 (", round(res.pca$eig[1,2], 1), "%)")
DIM_2 <- paste("Dim 2 (", round(res.pca$eig[2,2], 1), "%)")

#Draw ellipses around the clusters and highlighting the ellipsoids rather than the data points
centroids <- aggregate(cbind(PC1,PC2)~Subject,PCs.ID,mean)
conf.rgn  <- do.call(rbind,lapply(unique(PCs.ID$Subject),function(t)
  data.frame(Subject=as.character(t),
             ellipse(cov(PCs.ID[PCs.ID$Subject==t,1:2]),
                     centre=as.matrix(centroids[t,2:3]),
                     level=0.95),
             stringsAsFactors=FALSE)))

#Then make the figure
PCA_FIG_Mice <- 
  ggplot(PCs.ID, aes(x=PC1, y=PC2, color = Subject), guide = FALSE) +
  geom_hline(aes(yintercept=0), linetype="dashed", lwd = 0.1) +
  geom_vline(aes(xintercept=0), linetype="dashed", lwd = 0.1) +
  geom_path(data=conf.rgn, alpha=0.2, size = 0) +
  geom_polygon(data=conf.rgn, aes(fill = Subject),
               alpha=0.1,
               size = 0.1,
               show.legend = FALSE) + 
  geom_point(size=0.4) +
  theme_bw() +
  ggtitle("b) - Mouse") +
  scale_color_manual(labels=c("Vitro", "Vivo"),
                     values = c("#046C9A", "red")) +
  scale_fill_manual(labels=c("Vitro", "Vivo"),
                    values = c("#046C9A", "red"),
                    guide = FALSE) +
  ylab(DIM_2) +
  xlab(DIM_1) + 
  theme(panel.grid.major = element_blank(),
        panel.border = element_rect(colour = "black", size=1),
        panel.grid.minor = element_blank(),
        axis.title.x  = element_text(size=10, family = "serif"),
        axis.title.y  = element_text(size=10, family = "serif"),
        plot.title = element_text(size=8, hjust = 0, family = "serif"),
        axis.text.y  = element_text(size=5, family = "serif"),
        axis.text.x  = element_text(size=5, family = "serif"),
        legend.position="none",
        legend.background = element_blank(),
        legend.title = element_blank(),
        legend.text = element_text(size=8, family = "serif"),
        legend.key.size = unit(0.3, "cm"),
        legend.key = element_blank())

FIG <- arrangeGrob(PCA_FIG_Cows,
                   PCA_FIG_Mice,
                   ncol = 1)

grid.arrange(PCA_FIG_Cows, PCA_FIG_Mice,ncol = 1)
```

```
ggsave(FIG,
       file="Figures/Classification_Vitro_Vivo.png",
       width = 3.23,
       height=5,
       units = "in",
       dpi = 600)
```

##### Classify Stage for each of the species

Our last set of RF models aimed to classify developmental stage for data from mice and cows and humans. Here data from each collection type (i.e., *in vitro* vs. *in vivo*) were analysed separately.

###### In vitro samples

```
stage_Data <- data
stage_Data$Stage <- as.factor(IDs$Stage)
names(stage_Data) <- make.names(names(stage_Data))

stage_Data_cows <- stage_Data[which(IDs$Species == "cow" & IDs$Collection == "vitro"),]
stage_Data_cows$Stage <- factor(stage_Data_cows$Stage)
stage_Data_mice <- stage_Data[which(IDs$Species == "mouse" & IDs$Collection == "vitro"),]
stage_Data_mice$Stage <- factor(stage_Data_mice$Stage)
stage_Data_humans <- stage_Data[which(IDs$Species == "human"),]
stage_Data_humans$Stage <- factor(stage_Data_humans$Stage)


#Run the random forest model identifying stage in cows
stage.mod.cows <- randomForest(y = stage_Data_cows$Stage,
                               x = stage_Data_cows[, colnames(stage_Data_cows) != "Stage"],
                               mtry =  14,
                               ntree= 20000,
                               importance=TRUE,
                               proximity = TRUE,
                               keep.forest=TRUE,
                               replace = TRUE)
stage.mod.cows
```

```
## 
## Call:
##  randomForest(x = stage_Data_cows[, colnames(stage_Data_cows) !=      "Stage"], y = stage_Data_cows$Stage, ntree = 20000, mtry = 14,      replace = TRUE, importance = TRUE, proximity = TRUE, keep.forest = TRUE) 
##                Type of random forest: classification
##                      Number of trees: 20000
## No. of variables tried at each split: 14
## 
##         OOB estimate of  error rate: 20.83%
## Confusion matrix:
##     16C 2C 4C 8C BL MII class.error
## 16C   2  0  0  0  1   0   0.3333333
## 2C    0  3  0  0  0   0   0.0000000
## 4C    0  0  1  0  0   2   0.6666667
## 8C    0  0  1  2  0   0   0.3333333
## BL    0  0  0  0  9   0   0.0000000
## MII   0  0  1  0  0   2   0.3333333
```

```
#Run the random forest model identifying stage in mice
stage.mod.mice <- randomForest(y = stage_Data_mice$Stage,
                               x = stage_Data_mice[, colnames(stage_Data_mice) != "Stage"],
                               mtry =  14,
                               ntree= 20000,
                               importance=TRUE,
                               proximity = TRUE,
                               keep.forest=TRUE,
                               replace = TRUE)
stage.mod.mice
```

```
## 
## Call:
##  randomForest(x = stage_Data_mice[, colnames(stage_Data_mice) !=      "Stage"], y = stage_Data_mice$Stage, ntree = 20000, mtry = 14,      replace = TRUE, importance = TRUE, proximity = TRUE, keep.forest = TRUE) 
##                Type of random forest: classification
##                      Number of trees: 20000
## No. of variables tried at each split: 14
## 
##         OOB estimate of  error rate: 4.55%
## Confusion matrix:
##    2C 4C 8C BL MO class.error
## 2C  4  0  0  0  0        0.00
## 4C  0  4  0  0  0        0.00
## 8C  0  0  3  0  1        0.25
## BL  0  0  0  4  0        0.00
## MO  0  0  0  0  6        0.00
```

```
#Run the random forest model identifying stage in humans
stage.mod.humans <- randomForest(y = stage_Data_humans$Stage,
                                 x = stage_Data_humans[, colnames(stage_Data_humans) != "Stage"],
                                 mtry =  14,
                                 ntree= 20000,
                                 importance=TRUE,
                                 proximity = TRUE,
                                 keep.forest=TRUE,
                                 replace = TRUE)
stage.mod.humans
```

```
## 
## Call:
##  randomForest(x = stage_Data_humans[, colnames(stage_Data_humans) !=      "Stage"], y = stage_Data_humans$Stage, ntree = 20000, mtry = 14,      replace = TRUE, importance = TRUE, proximity = TRUE, keep.forest = TRUE) 
##                Type of random forest: classification
##                      Number of trees: 20000
## No. of variables tried at each split: 14
## 
##         OOB estimate of  error rate: 30%
## Confusion matrix:
##     2C 4C 8C BL MII MO class.error
## 2C   0  3  0  0   0  0   1.0000000
## 4C   0  5  0  0   0  0   0.0000000
## 8C   0  0  3  0   0  0   0.0000000
## BL   0  0  0  2   0  1   0.3333333
## MII  1  0  0  0   2  0   0.3333333
## MO   0  0  0  1   0  2   0.3333333
```

```
####################################
# PCA on proximity matrix for cows

res.pca <- PCA(stage.mod.cows$proximity, graph = FALSE) # Conduct a PCA on the proximity matrix
PC1 <- res.pca$ind$coord[,1] #Store individual coordinates of PC1 as a vector
PC2 <- res.pca$ind$coord[,2] #Store individual coordinates of PC2 as a vector
PCs.ID <- data.frame(cbind(PC1,PC2)) #Bind the coordinates together as a dataframe
PCs.ID$Subject <- stage_Data_cows$Stage #Add in Stage to the df
PCs.ID$Group <- factor(IDs[which(IDs$Species == "cow" & IDs$Collection == "vitro"),"Stage"])

DIM_1 <- paste("Dim 1 (", round(res.pca$eig[1,2], 1), "%)")
DIM_2 <- paste("Dim 2 (", round(res.pca$eig[2,2], 1), "%)")

#Draw ellipses around the clusters and highlighting the elipsoids rather than the data points
centroids <- aggregate(cbind(PC1,PC2)~Group,PCs.ID,mean)
conf.rgn  <- do.call(rbind,lapply(unique(PCs.ID$Group),function(t)
  data.frame(Group=as.character(t),
             ellipse(cov(PCs.ID[PCs.ID$Group==t,1:2]),
                     centre=as.matrix(centroids[t,2:3]),
                     level=0.95),
             stringsAsFactors=FALSE)))

#Then make the figure
PCA_FIG_Cows <- 
  ggplot(PCs.ID, aes(x=PC1, y=PC2), guide = FALSE) +
  geom_hline(aes(yintercept=0), linetype="dashed", lwd = 0.1) +
  geom_vline(aes(xintercept=0), linetype="dashed", lwd = 0.1) +
  geom_path(data=conf.rgn, alpha=0.2, size = 0) +
  geom_polygon(data=conf.rgn,
               aes(fill = Group, colour = Group),
               alpha=0.1,
               size = 0.1,
               show.legend = FALSE) + 
  geom_point(size=0.4, aes(color = Subject)) +
  theme_bw() +
  scale_color_viridis(discrete = TRUE) +
  scale_fill_viridis(guide = FALSE, discrete = T) +
  ggtitle("a) - Bovine") +
  ylab(DIM_2) +
  xlab(DIM_1) + 
  theme(panel.grid.major = element_blank(),
        panel.border = element_rect(colour = "black", size=1),
        panel.grid.minor = element_blank(),
        axis.title.x  = element_text(size=10, family = "serif"),
        axis.title.y  = element_text(size=10, family = "serif"),
        plot.title = element_text(size=8, hjust = 0, family = "serif"),
        axis.text.y  = element_text(size=5, family = "serif"),
        axis.text.x  = element_text(size=5, family = "serif"),
        legend.position= "right",
        legend.background = element_blank(),
        legend.title = element_blank(),
        legend.text = element_text(size=8, family = "serif"),
        legend.key.size = unit(0.3, "cm"),
        legend.key = element_blank())

####################################
# PCA on proximity matrix for mice

res.pca <- PCA(stage.mod.mice$proximity, graph = FALSE) # Conduct a PCA on the proximity matrix
PC1 <- res.pca$ind$coord[,1] #Store individual coordinates of PC1 as a vector
PC2 <- res.pca$ind$coord[,2] #Store individual coordinates of PC2 as a vector
PCs.ID <- data.frame(cbind(PC1,PC2)) #Bind the coordinates together as a dataframe
PCs.ID$Subject <- stage_Data_mice$Stage #Add in Season to the df
PCs.ID$Group <- factor(IDs[which(IDs$Species == "mouse" & IDs$Collection == "vitro"),"Stage"])

DIM_1 <- paste("Dim 1 (", round(res.pca$eig[1,2], 1), "%)")
DIM_2 <- paste("Dim 2 (", round(res.pca$eig[2,2], 1), "%)")

#Draw ellipses around the clusters and highlighting the elipsoids rather than the data points
centroids <- aggregate(cbind(PC1,PC2)~Group,PCs.ID,mean)
conf.rgn  <- do.call(rbind,lapply(unique(PCs.ID$Group),function(t)
  data.frame(Group=as.character(t),
             ellipse(cov(PCs.ID[PCs.ID$Group==t,1:2]),
                     centre=as.matrix(centroids[t,2:3]),
                     level=0.95),
             stringsAsFactors=FALSE)))

#Then make the figure
PCA_FIG_Mice <- 
  ggplot(PCs.ID, aes(x=PC1, y=PC2), guide = FALSE) +
  geom_hline(aes(yintercept=0), linetype="dashed", lwd = 0.1) +
  geom_vline(aes(xintercept=0), linetype="dashed", lwd = 0.1) +
  geom_path(data=conf.rgn, alpha=0.2, size = 0) +
  geom_polygon(data=conf.rgn,
               aes(fill = Group, colour = Group),
               alpha=0.1,
               size = 0.1,
               show.legend = FALSE) + 
  geom_point(size=0.4, aes(color = Subject)) +
  theme_bw() +
  scale_color_viridis(discrete = TRUE) +
  scale_fill_viridis(guide = FALSE, discrete = T) +
  ggtitle("b) - Mouse") +
  ylab(DIM_2) +
  xlab(DIM_1) + 
  theme(panel.grid.major = element_blank(),
        panel.border = element_rect(colour = "black", size=1),
        panel.grid.minor = element_blank(),
        axis.title.x  = element_text(size=10, family = "serif"),
        axis.title.y  = element_text(size=10, family = "serif"),
        plot.title = element_text(size=8, hjust = 0, family = "serif"),
        axis.text.y  = element_text(size=5, family = "serif"),
        axis.text.x  = element_text(size=5, family = "serif"),
        legend.position= "right",
        legend.background = element_blank(),
        legend.title = element_blank(),
        legend.text = element_text(size=8, family = "serif"),
        legend.key.size = unit(0.3, "cm"),
        legend.key = element_blank())

####################################
# PCA on proximity matrix for humans

res.pca <- PCA(stage.mod.humans$proximity, graph = FALSE) # Conduct a PCA on the proximity matrix
PC1 <- res.pca$ind$coord[,1] #Store individual coordinates of PC1 as a vector
PC2 <- res.pca$ind$coord[,2] #Store individual coordinates of PC2 as a vector
PCs.ID <- data.frame(cbind(PC1,PC2)) #Bind the coordinates together as a dataframe
PCs.ID$Subject <- stage_Data_humans$Stage #Add in Season to the df
PCs.ID$Group <- factor(IDs[which(IDs$Species == "human"),"Stage"])

DIM_1 <- paste("Dim 1 (", round(res.pca$eig[1,2], 1), "%)")
DIM_2 <- paste("Dim 2 (", round(res.pca$eig[2,2], 1), "%)")

#Draw ellipses around the clusters and highlighting the elipsoids rather than the data points
centroids <- aggregate(cbind(PC1,PC2)~Group,PCs.ID,mean)
conf.rgn  <- do.call(rbind,lapply(unique(PCs.ID$Group),function(t)
  data.frame(Group=as.character(t),
             ellipse(cov(PCs.ID[PCs.ID$Group==t,1:2]),
                     centre=as.matrix(centroids[t,2:3]),
                     level=0.95),
             stringsAsFactors=FALSE)))

#Then make the figure
PCA_FIG_Humans <- 
  ggplot(PCs.ID, aes(x=PC1, y=PC2), guide = FALSE) +
  geom_hline(aes(yintercept=0), linetype="dashed", lwd = 0.1) +
  geom_vline(aes(xintercept=0), linetype="dashed", lwd = 0.1) +
  geom_path(data=conf.rgn, alpha=0.2, size = 0) +
  geom_polygon(data=conf.rgn,
               aes(fill = Group, colour = Group),
               alpha=0.1,
               size = 0.1,
               show.legend = FALSE) + 
  geom_point(size=0.4, aes(color = Subject)) +
  theme_bw() +
  scale_color_viridis(discrete = TRUE) +
  scale_fill_viridis(guide = FALSE, discrete = T) +
  ggtitle("c) - Humans") +
  ylab(DIM_2) +
  xlab(DIM_1) + 
  theme(panel.grid.major = element_blank(),
        panel.border = element_rect(colour = "black", size=1),
        panel.grid.minor = element_blank(),
        axis.title.x  = element_text(size=10, family = "serif"),
        axis.title.y  = element_text(size=10, family = "serif"),
        plot.title = element_text(size=8, hjust = 0, family = "serif"),
        axis.text.y  = element_text(size=5, family = "serif"),
        axis.text.x  = element_text(size=5, family = "serif"),
        legend.position= "right",
        legend.background = element_blank(),
        legend.title = element_blank(),
        legend.text = element_text(size=8, family = "serif"),
        legend.key.size = unit(0.3, "cm"),
        legend.key = element_blank())

FIG <- arrangeGrob(PCA_FIG_Cows,
                   PCA_FIG_Mice,
                   PCA_FIG_Humans,
                   ncol = 1)

ggsave(FIG,
       file="Figures/Classification_Stage_Vitro.png",
       width = 4,
       height=7,
       units = "in",
       dpi = 600)
```

###### In vivo samples

```
stage_Data <- data
stage_Data$Stage <- as.factor(IDs$Stage)
names(stage_Data) <- make.names(names(stage_Data))

stage_Data_cows <- stage_Data[which(IDs$Species == "cow" & IDs$Collection == "vivo"),]
stage_Data_cows$Stage <- factor(stage_Data_cows$Stage)
stage_Data_mice <- stage_Data[which(IDs$Species == "mouse" & IDs$Collection == "vivo"),]
stage_Data_mice$Stage <- factor(stage_Data_mice$Stage)


#Run the random forest model identifying stage in cows
stage.mod.cows <- randomForest(y = stage_Data_cows$Stage,
                               x = stage_Data_cows[, colnames(stage_Data_cows) != "Stage"],
                               mtry =  14,
                               ntree= 20000,
                               importance=TRUE,
                               proximity = TRUE,
                               keep.forest=TRUE,
                               replace = TRUE)

stage.mod.cows
```

```
## 
## Call:
##  randomForest(x = stage_Data_cows[, colnames(stage_Data_cows) !=      "Stage"], y = stage_Data_cows$Stage, ntree = 20000, mtry = 14,      replace = TRUE, importance = TRUE, proximity = TRUE, keep.forest = TRUE) 
##                Type of random forest: classification
##                      Number of trees: 20000
## No. of variables tried at each split: 14
## 
##         OOB estimate of  error rate: 75%
## Confusion matrix:
##     16C 2C 4C 8C BL MII class.error
## 16C   1  0  0  1  0   0         0.5
## 2C    0  0  1  0  0   1         1.0
## 4C    0  2  0  0  0   0         1.0
## 8C    2  0  0  0  0   0         1.0
## BL    0  0  0  0  2   0         0.0
## MII   0  2  0  0  0   0         1.0
```

```
varImpPlot(stage.mod.cows, type=1, scale = FALSE)
```

```
#Run the random forest model identifying stage in mice
stage.mod.mice <- randomForest(y = stage_Data_mice$Stage,
                               x = stage_Data_mice[, colnames(stage_Data_mice) != "Stage"],
                               mtry =  14,
                               ntree= 20000,
                               importance=TRUE,
                               proximity = TRUE,
                               keep.forest=TRUE,
                               replace = TRUE)

stage.mod.mice
```

```
## 
## Call:
##  randomForest(x = stage_Data_mice[, colnames(stage_Data_mice) !=      "Stage"], y = stage_Data_mice$Stage, ntree = 20000, mtry = 14,      replace = TRUE, importance = TRUE, proximity = TRUE, keep.forest = TRUE) 
##                Type of random forest: classification
##                      Number of trees: 20000
## No. of variables tried at each split: 14
## 
##         OOB estimate of  error rate: 11.76%
## Confusion matrix:
##     2C 4C 8C BL MII MO class.error
## 2C   4  0  0  0   0  0           0
## 4C   0  4  0  0   0  0           0
## 8C   0  2  0  0   0  0           1
## BL   0  0  0  3   0  0           0
## MII  0  0  0  0   2  0           0
## MO   0  0  0  0   0  2           0
```

```
varImpPlot(stage.mod.mice, type=1, scale = FALSE)

####################################
# PCA on proximity matrix for cows

res.pca <- PCA(stage.mod.cows$proximity, graph = FALSE) # Conduct a PCA on the proximity matrix
PC1 <- res.pca$ind$coord[,1] #Store individual coordinates of PC1 as a vector
PC2 <- res.pca$ind$coord[,2] #Store individual coordinates of PC2 as a vector
PCs.ID <- data.frame(cbind(PC1,PC2)) #Bind the coordinates together as a dataframe
PCs.ID$Subject <- stage_Data_cows$Stage #Add in Season to the df
PCs.ID$Group <- factor(IDs[which(IDs$Species == "cow" & IDs$Collection == "vivo"),"Stage"])

DIM_1 <- paste("Dim 1 (", round(res.pca$eig[1,2], 1), "%)")
DIM_2 <- paste("Dim 2 (", round(res.pca$eig[2,2], 1), "%)")

#Draw ellipses around the clusters and highlighting the elipsoids rather than the data points
centroids <- aggregate(cbind(PC1,PC2)~Group,PCs.ID,mean)
conf.rgn  <- do.call(rbind,lapply(unique(PCs.ID$Group),function(t)
  data.frame(Group=as.character(t),
             ellipse(cov(PCs.ID[PCs.ID$Group==t,1:2]),
                     centre=as.matrix(centroids[t,2:3]),
                     level=0.95),
             stringsAsFactors=FALSE)))

#Then make the figure
PCA_FIG_Cows <- 
  ggplot(PCs.ID, aes(x=PC1, y=PC2), guide = FALSE) +
  geom_hline(aes(yintercept=0), linetype="dashed", lwd = 0.1) +
  geom_vline(aes(xintercept=0), linetype="dashed", lwd = 0.1) +
  geom_path(data=conf.rgn, alpha=0.2, size = 0) +
  geom_polygon(data=conf.rgn,
               aes(fill = Group, colour = Group),
               alpha=0.1,
               size = 0.1,
               show.legend = FALSE) + 
  geom_point(size=0.4, aes(color = Subject)) +
  theme_bw() +
  scale_color_viridis(discrete = TRUE) +
  scale_fill_viridis(guide = FALSE, discrete = T) +
  ggtitle("a) - Bovine") +
  ylab(DIM_2) +
  xlab(DIM_1) + 
  theme(panel.grid.major = element_blank(),
        panel.border = element_rect(colour = "black", size=1),
        panel.grid.minor = element_blank(),
        axis.title.x  = element_text(size=10, family = "serif"),
        axis.title.y  = element_text(size=10, family = "serif"),
        plot.title = element_text(size=8, hjust = 0, family = "serif"),
        axis.text.y  = element_text(size=5, family = "serif"),
        axis.text.x  = element_text(size=5, family = "serif"),
        legend.position= "right",
        legend.background = element_blank(),
        legend.title = element_blank(),
        legend.text = element_text(size=8, family = "serif"),
        legend.key.size = unit(0.3, "cm"),
        legend.key = element_blank())

####################################
# PCA on proximity matrix for mice

res.pca <- PCA(stage.mod.mice$proximity, graph = FALSE) # Conduct a PCA on the proximity matrix
PC1 <- res.pca$ind$coord[,1] #Store individual coordinates of PC1 as a vector
PC2 <- res.pca$ind$coord[,2] #Store individual coordinates of PC2 as a vector
PCs.ID <- data.frame(cbind(PC1,PC2)) #Bind the coordinates together as a dataframe
PCs.ID$Subject <- stage_Data_mice$Stage #Add in Season to the df
PCs.ID$Group <- factor(IDs[which(IDs$Species == "mouse" & IDs$Collection == "vivo"),"Stage"])

DIM_1 <- paste("Dim 1 (", round(res.pca$eig[1,2], 1), "%)")
DIM_2 <- paste("Dim 2 (", round(res.pca$eig[2,2], 1), "%)")

#Draw ellipses around the clusters and highlighting the elipsoids rather than the data points
centroids <- aggregate(cbind(PC1,PC2)~Group,PCs.ID,mean)
conf.rgn  <- do.call(rbind,lapply(unique(PCs.ID$Group),function(t)
  data.frame(Group=as.character(t),
             ellipse(cov(PCs.ID[PCs.ID$Group==t,1:2]),
                     centre=as.matrix(centroids[t,2:3]),
                     level=0.95),
             stringsAsFactors=FALSE)))

#Then make the figure
PCA_FIG_Mice <- 
  ggplot(PCs.ID, aes(x=PC1, y=PC2), guide = FALSE) +
  geom_hline(aes(yintercept=0), linetype="dashed", lwd = 0.1) +
  geom_vline(aes(xintercept=0), linetype="dashed", lwd = 0.1) +
  geom_path(data=conf.rgn, alpha=0.2, size = 0) +
  geom_polygon(data=conf.rgn,
               aes(fill = Group, colour = Group),
               alpha=0.1,
               size = 0.1,
               show.legend = FALSE) + 
  geom_point(size=0.4, aes(color = Subject)) +
  theme_bw() +
  scale_color_viridis(discrete = TRUE) +
  scale_fill_viridis(guide = FALSE, discrete = T) +
  ggtitle("b) - Mouse") +
  ylab(DIM_2) +
  xlab(DIM_1) + 
  theme(panel.grid.major = element_blank(),
        panel.border = element_rect(colour = "black", size=1),
        panel.grid.minor = element_blank(),
        axis.title.x  = element_text(size=10, family = "serif"),
        axis.title.y  = element_text(size=10, family = "serif"),
        plot.title = element_text(size=8, hjust = 0, family = "serif"),
        axis.text.y  = element_text(size=5, family = "serif"),
        axis.text.x  = element_text(size=5, family = "serif"),
        legend.position= "right",
        legend.background = element_blank(),
        legend.title = element_blank(),
        legend.text = element_text(size=8, family = "serif"),
        legend.key.size = unit(0.3, "cm"),
        legend.key = element_blank())

FIG <- arrangeGrob(PCA_FIG_Cows,
                   PCA_FIG_Mice,
                   ncol = 1)
```

```
ggsave(FIG,
       file="Figures/Classification_Stage_Vivo.png",
       width = 3.23,
       height=5,
       units = "in",
       dpi = 600)
```

---

#### Differentially Expressed Genes (DEG)

Differentially expressed genes (DEGs) of normalized data were identified using the R package `limma`. An adjusted p-value to correct for multiple testing was calculated using the Benjamini–Hochberg method. The most widely used correction for genomic studies is the Benjamini-Hochberg (BH) correction, that aims to control the FDR across significant genes.

##### Intra-specific DEGs

```
#Load in metadata
GENES <- read.csv("metabolism_genes.csv")
KEEPERS <- read.csv("metaboloepig_genes.csv")
#Convert to a numeric matrix
DATA <- as.matrix(t(data))


#Create the design matrix
design <- cbind(species = as.numeric(as.factor(IDs$Species)),
                stage = as.numeric(as.factor(IDs$Stage)),
                collection = as.numeric(as.factor(IDs$Collection)))

STAGE_NAMES <- unique(as.factor(IDs$Stage))

#Empty list to fill
RES <- list()

#Subset the data and create the appropriate design matrix
#Pick which species you're interested in cow = 1, human = 2, mouse = 3

for(i in 1:length(unique(IDs$Species))){
  SPECIES <- c(i)
  design2 <- design[which(design[,1] == SPECIES),]
  DATA_Species <- DATA[,which(design[,1] == SPECIES)]
  
  #################################
  # Do all the pairwise comparisons of the different stages for each collection type
  
  #The different combinations of stages to test
  stage_tests <- combn(unique(design2[,2]), 2)
  
  #Loop over the combinations of stage(s) 16C = 1; 2C = 2; 4C = 3; 8C = 4; BL = 5; MII = 6; MO = 7
  for(j in 1:ncol(stage_tests)){
    STAGES <- c(stage_tests[,j])
    design3 <- design2[which(design2[,2] %in% STAGES),]
    DATA_Stages <- DATA_Species[,which(design2[,2] %in% STAGES)]
    
    #Added this check because humans don't have in vivo
    if(SPECIES != 2){
      #Loop over collection type(s): vitro = 1; vivo = 2
      for(k in 1:2){
        COLLECTION <- k
        design4 <- design3[which(design3[,3] %in% COLLECTION),]
        DATA_test <- DATA_Stages[,which(design3[,3] %in% COLLECTION)]
        
        if(length(unique(design4[,2])) > 1){
          test <- lmFit(DATA_test, design = model.matrix(~ 1 + design4[,2]))
          test2 <- eBayes(test)
          test3 <- topTable(test2, number = length(test2$coefficients), sort.by = "none")
          results <- data.frame(test3$adj.P.Val)
          names(results) <- paste(unique(IDs$Species)[i],
                                  sort(unique(IDs$Collection))[k],
                                  paste(sort(unique(as.factor(IDs$Stage)))[STAGES][1], sort(unique(as.factor(IDs$Stage)))[STAGES][2], sep = "vs"),
                                  sep = "_")
          
          png(file=paste("Figures/Volcano_Plots/",
                         unique(IDs$Species)[i],
                         sort(unique(IDs$Collection))[k],
                         paste(sort(unique(as.factor(IDs$Stage)))[STAGES][1], sort(unique(as.factor(IDs$Stage)))[STAGES][2], sep = "vs"),
                         ".png", sep = ""),
              type="cairo",
              units = "in",
              width = 6, height = 6,
              res = 300) 
          volcanoplot(test2, col = ifelse(test3$adj.P.Val< 0.05, "red", "#046C9A"))
          dev.off()
          
          RES[[length(RES)+1]] <- results
        }
      } #Closes the loop over collection type
    } else {
      
      #Loop over collection type(s): vitro = 1; vivo = 2
      for(k in 1:2){
        COLLECTION <- k
        design4 <- design3[which(design3[,3] %in% COLLECTION),]
        DATA_test <- DATA_Stages[,which(design3[,3] %in% COLLECTION)]
        
        if(length(unique(design4[,2])) > 1){
          test <- lmFit(DATA_test, design = model.matrix(~ 1 + design4[,2]))
          test2 <- eBayes(test)
          test3 <- topTable(test2, number = length(test2$coefficients), sort.by = "none")
          results <- data.frame(test3$adj.P.Val)
          names(results) <- paste(unique(IDs$Species)[i],
                                  sort(unique(IDs$Collection))[k],
                                  paste(sort(unique(as.factor(IDs$Stage)))[STAGES][1], sort(unique(as.factor(IDs$Stage)))[STAGES][2], sep = "vs"),
                                  sep = "_")
          
          png(file=paste("Figures/Volcano_Plots/",
                         unique(IDs$Species)[i],
                         sort(unique(IDs$Collection))[k],
                         paste(sort(unique(as.factor(IDs$Stage)))[STAGES][1], sort(unique(as.factor(IDs$Stage)))[STAGES][2], sep = "vs"),
                         ".png", sep = ""),
              type="cairo",
              units = "in",
              width = 6, height = 6,
              res = 300) 
          volcanoplot(test2, col = ifelse(test3$adj.P.Val< 0.05, "red", "#046C9A"))
          dev.off()
          
          RES[[length(RES)+1]] <- results
        }
      } #Closes the loop over collection type
    }
  } #Closes the loop over the stage types
  
  
  #################################
  # Do all the pairwise comparisons of the different collection types for each DATA_Stages
  
  #Ignore these these for humans
  if(SPECIES != 2){
    #Loop over the combinations of stage(s) 16C = 1; 2C = 2; 4C = 3; 8C = 4; BL = 5; MII = 6; MO = 7
    for(m in 1:length(unique(design2[,2]))){
      STAGES <- unique(design2[,2])[m]
      design3 <- design2[which(design2[,2] %in% STAGES),]
      DATA_Stages <- DATA_Species[,which(design2[,2] %in% STAGES)]
      
      if(length(unique(design3[,3])) > 1){
        
        test <- lmFit(DATA_Stages, design = model.matrix(~ 1 + design3[,3]))
        test2 <- eBayes(test)
        test3 <- topTable(test2, number = length(test2$coefficients), sort.by = "none")
        results <- data.frame(test3$adj.P.Val)
        names(results) <- paste(unique(IDs$Species)[i],
                                sort(unique(as.factor(IDs$Stage)))[STAGES],
                                "vitro - vivo",
                                sep = "_")
        
        
        png(file=paste("Figures/Volcano_Plots/",
                       unique(IDs$Species)[i],
                       sort(unique(as.factor(IDs$Stage)))[STAGES],
                       "vitro - vivo",
                       ".png", sep = ""),
            type="cairo",
            units = "in",
            width = 6, height = 6,
            res = 300) 
        volcanoplot(test2, col = ifelse(test3$adj.P.Val< 0.05, "red", "#046C9A"))
        dev.off()
        
        
        RES[[length(RES)+1]] <- results
        
        
      }
    } #Closes the loop over the stage types
    
    
  }
} #Closes the top level loop


hist(unlist(RES), main = "Adjusted p-values")
```

```
RESULTS <- do.call(cbind, RES)
row.names(RESULTS) <- NAMES
write.csv(RESULTS, file = "DEG_Analysis_FINAL.csv", row.names = TRUE)
pvalues <- RESULTS
REDUCED <- RESULTS[(row.names(RESULTS) %in% stringr::str_trim(KEEPERS[,1])),]
write.csv(REDUCED, file = "DEG_Analysis_Reduced_FINAL.csv", row.names = TRUE)


############## Get info on up or down regulated
RES <- list()
#Subset the data and create the appropriate design matrix
#Pick which species you're interested in cow = 1, human = 2, mouse = 3
for(i in 1:length(unique(IDs$Species))){
  SPECIES <- c(i)
  design2 <- design[which(design[,1] == SPECIES),]
  DATA_Species <- DATA[,which(design[,1] == SPECIES)]
  
  #################################
  # Do all the pairwise comparisons of the different stages for each collection type
  
  #The different combinations of stages to test
  stage_tests <- combn(unique(design2[,2]), 2)
  
  #Loop over the combinations of stage(s) 16C = 1; 2C = 2; 4C = 3; 8C = 4; BL = 5; MII = 6; MO = 7
  for(j in 1:ncol(stage_tests)){
    STAGES <- c(stage_tests[,j])
    design3 <- design2[which(design2[,2] %in% STAGES),]
    DATA_Stages <- DATA_Species[,which(design2[,2] %in% STAGES)]
    
    
    #Added this because humans don't have in vivo
    if(SPECIES != 2){
      #Loop over collection type(s): vitro = 1; vivo = 2
      for(k in 1:2){
        COLLECTION <- k
        design4 <- design3[which(design3[,3] %in% COLLECTION),]
        DATA_test <- DATA_Stages[,which(design3[,3] %in% COLLECTION)]
        
        if(length(unique(design4[,2])) > 1){
          test <- lmFit(DATA_test, design = model.matrix(~ 1 + design4[,2]))
          test2 <- eBayes(test)
          test3 <- topTable(test2, number = length(test2$coefficients), sort.by = "none")
          results <- data.frame(test3$logFC)
          names(results) <- paste(unique(IDs$Species)[i],
                                  sort(unique(IDs$Collection))[k],
                                  paste(sort(unique(as.factor(IDs$Stage)))[STAGES][1], sort(unique(as.factor(IDs$Stage)))[STAGES][2], sep = "vs"),
                                  sep = "_")
          RES[[length(RES)+1]] <- results
        }
      } #Closes the loop over collection type
    } else {
      
      #Loop over collection type(s): vitro = 1; vivo = 2
      for(k in 1:2){
        COLLECTION <- k
        design4 <- design3[which(design3[,3] %in% COLLECTION),]
        DATA_test <- DATA_Stages[,which(design3[,3] %in% COLLECTION)]
        
        if(length(unique(design4[,2])) > 1){
          test <- lmFit(DATA_test, design = model.matrix(~ 1 + design4[,2]))
          test2 <- eBayes(test)
          test3 <- topTable(test2, number = length(test2$coefficients), sort.by = "none")
          results <- data.frame(test3$logFC)
          names(results) <- paste(unique(IDs$Species)[i],
                                  sort(unique(IDs$Collection))[k],
                                  paste(sort(unique(as.factor(IDs$Stage)))[STAGES][1], sort(unique(as.factor(IDs$Stage)))[STAGES][2], sep = "vs"),
                                  sep = "_")
          RES[[length(RES)+1]] <- results
        }
      } #Closes the loop over collection type
    }
  } #Closes the loop over the stage types
  
  
  
  
  
  #################################
  # Do all the pairwise comparisons of the different collection types for each DATA_Stages
  
  #Ignore these these for humans
  if(SPECIES != 2){
    #Loop over the combinations of stage(s) 16C = 1; 2C = 2; 4C = 3; 8C = 4; BL = 5; MII = 6; MO = 7
    for(m in 1:length(unique(design2[,2]))){
      STAGES <- unique(design2[,2])[m]
      design3 <- design2[which(design2[,2] %in% STAGES),]
      DATA_Stages <- DATA_Species[,which(design2[,2] %in% STAGES)]
      
      if(length(unique(design3[,3])) > 1){
        
        test <- lmFit(DATA_Stages, design = model.matrix(~ 1 + design3[,3]))
        test2 <- eBayes(test)
        test3 <- topTable(test2, number = length(test2$coefficients), sort.by = "none")
        results <- data.frame(test3$logFC)
        names(results) <- paste(unique(IDs$Species)[i],
                                sort(unique(as.factor(IDs$Stage)))[STAGES],
                                "vitro - vivo",
                                sep = "_")
        RES[[length(RES)+1]] <- results
        
      }
    } #Closes the loop over the stage types
  }
} #Closes the top level loop


RESULTS <- do.call(cbind, RES)
row.names(RESULTS) <- NAMES
write.csv(RESULTS, file = "DEG_Analysis_log2FC_FINAL.csv", row.names = TRUE)
REDUCED <- RESULTS[(row.names(RESULTS) %in% stringr::str_trim(KEEPERS[,1])),]
write.csv(REDUCED, file = "DEG_Analysis_log2FC_Reduced_FINAL.csv", row.names = TRUE)

# numbers of significant up and down regulated for each and volcano plots
REGULATION <- RESULTS[0,]

for(i in 1:ncol(RESULTS)){
  
  SIGS <- RESULTS[which(pvalues[,i] < 0.05),i]
  
  REGULATION[1,i] <- length(which(SIGS < 0))
  
  REGULATION[2,i] <- length(which(SIGS > 0))
}

row.names(REGULATION) <- c("down", "up")

write.csv(REGULATION, file = "Number_Sig_FINAL.csv", row.names = TRUE)
```

##### Inter-specific DEGs

```
RES <- list()

for(i in 1:length(unique(IDs$Stage))){
  STAGE <- c(i)
  design2 <- design[which(design[,2] == STAGE),]
  DATA_Stage <- DATA[,which(design[,2] == STAGE)]
  
  if(length(unique(design2[,1]))>1){
    #################################
    # Do all the pairwise comparisons of the different stages for each collection type
    
    #The different combinations of species to test
    species_tests <- combn(unique(design2[,1]), 2)
    
    for(j in 1:ncol(species_tests)){
      SPECIES <- c(species_tests[,j])
      design3 <- design2[which(design2[,1] %in% SPECIES),]
      DATA_Stages <- DATA_Stage[,which(design2[,1] %in% SPECIES)]
      
      
      design4 <- design3[which(design3[,3] == 2 & design3[,1] == 1),]
      design4 <- rbind(design4, design3[which(design3[,3] == 1 & design3[,1] == 2),])
      design4 <- rbind(design4, design3[which(design3[,3] == 2 & design3[,1] == 3),])
      DATA_Stages <- DATA_Stages[,as.numeric(row.names(plyr::match_df(as.data.frame(design3), as.data.frame(design4))))]
      
      
      test <- lmFit(DATA_Stages, design = model.matrix(~ 1 + design4[,1]))
      test2 <- eBayes(test)
      test3 <- topTable(test2, number = length(test2$coefficients), sort.by = "none")
      results <- data.frame(test3$adj.P.Val)
      names(results) <- paste(sort(unique(as.factor(IDs$Stage)))[i],
                              paste(sort(unique(as.factor(IDs$Species)))[SPECIES][1], sort(unique(as.factor(IDs$Species)))[SPECIES][2], sep = "vs"),
                              sep = "_")
      
      png(file=paste("Figures/Volcano_Plots/",
                     sort(unique(as.factor(IDs$Stage)))[i],
                     paste(sort(unique(as.factor(IDs$Species)))[SPECIES][1], sort(unique(as.factor(IDs$Species)))[SPECIES][2], sep = "vs"),
                     ".png", sep = ""),
          type="cairo",
          units = "in",
          width = 6, height = 6,
          res = 300) 
      volcanoplot(test2, col = ifelse(test3$adj.P.Val< 0.05, "red", "#046C9A"))
      dev.off()
      
      
      RES[[length(RES)+1]] <- results
      
    } # Ends loop over species to compare
  } #Closes the if statement to correct for stages that don't have potential inter-species comparisons
}


hist(unlist(RES), main = "Adjusted p-values")
```

```
RESULTS <- do.call(cbind, RES)
row.names(RESULTS) <- NAMES
write.csv(RESULTS, file = "DEG_Analysis_InterSpecies_FINAL.csv", row.names = TRUE)
pvalues <- RESULTS
REDUCED <- RESULTS[(row.names(RESULTS) %in% stringr::str_trim(KEEPERS[,1])),]
write.csv(REDUCED, file = "DEG_Analysis_InterSpecies_Reduced_FINAL.csv", row.names = TRUE)


#Empty list to fill with resuts
RES <- list()

for(i in 1:length(unique(IDs$Stage))){
  STAGE <- c(i)
  design2 <- design[which(design[,2] == STAGE),]
  DATA_Stage <- DATA[,which(design[,2] == STAGE)]
  
  if(length(unique(design2[,1]))>1){
    #################################
    # Do all the pairwise comparisons of the different stages for each collection type
    
    #The different combinations of species to test
    species_tests <- combn(unique(design2[,1]), 2)
    
    for(j in 1:ncol(species_tests)){
      SPECIES <- c(species_tests[,j])
      design3 <- design2[which(design2[,1] %in% SPECIES),]
      DATA_Stages <- DATA_Stage[,which(design2[,1] %in% SPECIES)]
      
      
      design4 <- design3[which(design3[,3] == 2 & design3[,1] == 1),]
      design4 <- rbind(design4, design3[which(design3[,3] == 1 & design3[,1] == 2),])
      design4 <- rbind(design4, design3[which(design3[,3] == 2 & design3[,1] == 3),])
      DATA_Stages <- DATA_Stages[,as.numeric(row.names(plyr::match_df(as.data.frame(design3), as.data.frame(design4))))]
      
      
      test <- lmFit(DATA_Stages, design = model.matrix(~ 1 + design4[,1]))
      test2 <- eBayes(test)
      test3 <- topTable(test2, number = length(test2$coefficients), sort.by = "none")
      results <- data.frame(test3$logFC)
      names(results) <- paste(sort(unique(as.factor(IDs$Stage)))[i],
                              paste(sort(unique(as.factor(IDs$Species)))[SPECIES][1], sort(unique(as.factor(IDs$Species)))[SPECIES][2], sep = "vs"),
                              sep = "_")
      
      png(file=paste("Figures/Volcano_Plots/",
                     sort(unique(as.factor(IDs$Stage)))[i],
                     paste(sort(unique(as.factor(IDs$Species)))[SPECIES][1], sort(unique(as.factor(IDs$Species)))[SPECIES][2], sep = "vs"),
                     ".png", sep = ""),
          type="cairo",
          units = "in",
          width = 6, height = 6,
          res = 300) 
      volcanoplot(test2, col = ifelse(test3$adj.P.Val< 0.05, "red", "#046C9A"))
      dev.off()
      
      RES[[length(RES)+1]] <- results
      
    } # Ends loop over species to compare
  } #Closes the if statement to correct for stages that don't have potential inter-species comparisons
}


RESULTS <- do.call(cbind, RES)
row.names(RESULTS) <- NAMES
write.csv(RESULTS, file = "DEG_Analysis_InterSpecies_logFC_FINAL.csv", row.names = TRUE)
REDUCED <- RESULTS[(row.names(RESULTS) %in% stringr::str_trim(KEEPERS[,1])),]
write.csv(REDUCED, file = "DEG_Analysis_InterSpecies_logFC_Reduced_FINAL.csv", row.names = TRUE)


# numbers of significant up and down regulated for each and volcano plots
REGULATION <- RESULTS[0,]

for(i in 1:ncol(RESULTS)){
  
  SIGS <- RESULTS[which(pvalues[,i] < 0.05),i]
  
  REGULATION[1,i] <- length(which(SIGS < 0))
  
  REGULATION[2,i] <- length(which(SIGS > 0))
}

row.names(REGULATION) <- c("down", "up")

write.csv(REGULATION, file = "InterSpecies_Number_Sig_FINAL.csv", row.names = TRUE)
```

---

#### ROAST

We accessed information of all epigenetic and metabolic pathways for humans (Reactome terms: “epigenetic regulation of gene expression”, “metabolism”, “metabolism of proteins” and “metabolism of RNA”) from the Reactome pathways database version 74 (Jassal et al., 2020). Reactome pathways are arranged into several tiers, the Reactome term “epigenetic regulation of gene expression” (Reactome ID: R-HSA-212165.2), included curated pathways involving 122 genes; the Reactome term “metabolism” (Reactome ID: R-HSA-1430728.10) involved 2,210 genes; the curated pathways of the Reactome term “metabolism of proteins” (Reactome ID: R-HSA-392499.7) involved 2,095 genes; and the curated pathways of the Reactome term “metabolism of RNA” (Reactome ID: R-HSA-8953854.4) involved 739 genes. The rotation gene set (ROAST) algorithm was used to perform self-contained gene set analysis of each metabolic pathway, for the different developmental stages and species (Wu et al., 2010).

```
RES <- list()

#Subset the data and create the appropriate design matrix
#Pick which species you're interested in cow = 1, human = 2, mouse = 3

for(i in 1:length(unique(IDs$Species))){
  SPECIES <- c(i)
  design2 <- design[which(design[,1] == SPECIES),]
  DATA_Species <- DATA[,which(design[,1] == SPECIES)]
  
  #################################
  # Do all the pairwise comparisons of the different stages for each collection type
  
  #The different combinations of stages to test
  stage_tests <- combn(unique(design2[,2]), 2)
  
  #Loop over the combinations of stage(s) 16C = 1; 2C = 2; 4C = 3; 8C = 4; BL = 5; MII = 6; MO = 7
  for(j in 1:ncol(stage_tests)){
    STAGES <- c(stage_tests[,j])
    design3 <- design2[which(design2[,2] %in% STAGES),]
    DATA_Stages <- DATA_Species[,which(design2[,2] %in% STAGES)]
    
    #Added this because humans don't have in vivo
    if(SPECIES != 2){
      #Loop over collection type(s): vitro = 1; vivo = 2
      for(k in 1:2){
        COLLECTION <- k
        design4 <- design3[which(design3[,3] %in% COLLECTION),]
        DATA_test <- DATA_Stages[,which(design3[,3] %in% COLLECTION)]
        
        if(length(unique(design4[,2])) > 1){
          # Loop over the different pathways that are going to be tested
          for(l in 1:ncol(GENES)){
            #Select genes in the pathway
            Index <- which(NAMES %in% GENES[,l])
            
            #Conduct the test (REMEBER TO CHANGE CONTRAST 1 = species, 2 = stages, 3 = collection type)
            ROAST <- mroast(DATA_test,Index,design4[,1:2],contrast=2)
            ROAST$Stages <- paste(sort(unique(as.factor(IDs$Stage)))[STAGES][1], sort(unique(as.factor(IDs$Stage)))[STAGES][2], sep = "_")
            ROAST$Collection <- sort(unique(IDs$Collection))[k]
            ROAST$Pathway <- names(GENES)[l]
            ROAST$Species <- unique(IDs$Species)[i]
            
            #Save the results
            RES[[length(RES)+1]] <- ROAST
            
          } }#Closes the loop over the different pathways
      } #Closes the loop over collection type
    } else {
      
      #Loop over collection type(s): vitro = 1; vivo = 2
      for(k in 1){
        COLLECTION <- k
        design4 <- design3[which(design3[,3] %in% COLLECTION),]
        DATA_test <- DATA_Stages[,which(design3[,3] %in% COLLECTION)]
        
        # Loop over the different pathways that are going to be tested
        for(l in 1:ncol(GENES)){
          #Select genes in the pathway
          Index <- which(NAMES %in% GENES[,l])
          
          #Conduct the test (REMEBER TO CHANGE CONTRAST 1 = species, 2 = stages, 3 = collection type)
          ROAST <- mroast(DATA_test,Index,design4[,1:2],contrast=2)
          ROAST$Stages <- paste(sort(unique(as.factor(IDs$Stage)))[STAGES][1], sort(unique(as.factor(IDs$Stage)))[STAGES][2], sep = "_")
          ROAST$Collection <- rev(unique(IDs$Collection))[k]
          ROAST$Pathway <- names(GENES)[l]
          ROAST$Species <- unique(IDs$Species)[i]
          
          #Save the results
          RES[[length(RES)+1]] <- ROAST
          
        } #Closes the loop over the different pathways
      } #Closes the loop over collection type
    } # Closes the else from the ifelse
  } #Closes the loop over the stage types
  
  #################################
  # Do all the pairwise comparisons of the different stages for each collection type
  
  #Ignore these these for humans
  if(SPECIES != 2){
    #Loop over the combinations of stage(s) 16C = 1; 2C = 2; 4C = 3; 8C = 4; BL = 5; MII = 6; MO = 7
    for(m in 1:length(unique(design2[,2]))){
      STAGES <- unique(design2[,2])[m]
      design3 <- design2[which(design2[,2] %in% STAGES),]
      DATA_Stages <- DATA_Species[,which(design2[,2] %in% STAGES)]
      
      if(length(unique(design3[,3])) > 1){
        # Loop over the different pathways that are going to be tested
        for(n in 1:ncol(GENES)){
          #Select genes in the pathway
          Index <- which(NAMES %in% GENES[,n])
          
          #Conduct the test (REMEBER TO CHANGE CONTRAST 1 = species, 2 = stages, 3 = collection type)
          ROAST <- mroast(DATA_Stages,Index,design3[,c(1,3)],contrast=3)
          
          ROAST$Stages <- sort(unique(as.factor(IDs$Stage)))[STAGES]
          ROAST$Collection <- "vitro - vivo"
          ROAST$Pathway <- names(GENES)[n]
          ROAST$Species <- unique(IDs$Species)[i]
          
          #Save the results
          RES[[length(RES)+1]] <- ROAST
          
        } } #Closes the loop over the different pathways
    } #Closes the loop over the stage types
  } #Closes the if statement
} #Closes the top level loop

test <- do.call(rbind, RES)

write.csv(test, file = "Roast_FINAL.csv", row.names = FALSE)
```

#### Expression Figures

The following script recreates the expression pannels shown in Figure 4 in the main text.

```
IDs_2 <- read.csv("all_extra2.csv")

for(i in 1:length(NAMES)){
  MEANS <- aggregate(DATA[i,], by = IDs_2, FUN = "mean")
  CI <- function(x) sqrt(var(x)/length(x))*1.96
  CIs <- aggregate(DATA[i,], by = IDs_2, FUN = "CI")
  MEANS$CIs <- CIs$x
  MEANS <- MEANS[-which(MEANS$Species == "cow" & MEANS$Collection == "vitro"),]
  MEANS <- MEANS[-which(MEANS$Species == "mouse" & MEANS$Collection == "vitro"),]
  
  path <- paste("Figures/Expression_Figures/", NAMES[i], "_Expression.png", sep = "")
  
  YLAB <- paste(NAMES[i], " Adjusted expression", sep = "")
  
  FIG <- 
    ggplot(MEANS, aes(x=Stage, y=x, color = Species), guide = FALSE) +
    geom_line(size = 1.5) +
    geom_errorbar(aes(ymin=x-CIs, ymax=x+CIs), width=.1) +
    theme_bw() +
    scale_color_manual(labels=c("Bovine", "Human", "Mouse"), values = c("#e6c141", "#8a3bb8", "#3c7a47")) +
    ylab(YLAB) +
    xlab("Stage") + 
    theme(panel.grid.major = element_blank(),
          panel.border = element_rect(colour = "black", size=1),
          panel.grid.minor = element_blank(),
          axis.title.x  = element_blank(),
          axis.title.y  = element_text(size=10),
          plot.title = element_text(size=8, hjust = 0),
          axis.text.y  = element_text(size=5),
          axis.text.x  = element_text(size=7),
          legend.position="top",
          legend.background = element_blank(),
          legend.title = element_blank(),
          legend.text = element_text(size=8),
          legend.key.size = unit(0.3, "cm"),
          legend.key = element_blank(),
          panel.background = element_rect(fill = "transparent"),
          plot.background = element_rect(fill = "transparent", color = NA)) +
    scale_x_continuous(breaks = 1:6, labels = c("MII", "2C", "4C", "8C", "16C/MO", "BL"))
  
  ggsave(FIG,
         file=path,
         bg = "transparent",
         width = 3.23,
         height=3,
         units = "in",
         dpi = 600)
}


for(i in 1:length(NAMES)){
  MEANS <- aggregate(DATA[i,], by = IDs_2, FUN = "mean")
  CI <- function(x) sqrt(var(x)/length(x))*1.96
  CIs <- aggregate(DATA[i,], by = IDs_2, FUN = "CI")
  MEANS$CIs <- CIs$x
  
  path <- paste("Figures/Expression_Figures_2/", NAMES[i], "_Expression.png", sep = "")
  
  YLAB <- paste(NAMES[i], " Adjusted expression", sep = "")
  
  FIG <- 
    ggplot(MEANS, aes(x=Stage, y=x, color = Species), guide = FALSE) +
    geom_line(size = 1, aes(linetype = Collection)) +
    geom_errorbar(aes(ymin=x-CIs, ymax=x+CIs), width=.1) +
    theme_bw() +
    scale_color_manual(labels=c("Bovine", "Human", "Mouse"), values = c("#e6c141", "#8a3bb8", "#3c7a47")) +
    scale_linetype_manual(values = c("dashed", "solid"), guide = FALSE) +
    ylab(YLAB) +
    xlab("Stage") + 
    theme(panel.grid.major = element_blank(),
          panel.border = element_rect(colour = "black", size=1),
          panel.grid.minor = element_blank(),
          axis.title.x  = element_blank(),
          axis.title.y  = element_text(size=10),
          plot.title = element_text(size=8, hjust = 0),
          axis.text.y  = element_text(size=5),
          axis.text.x  = element_text(size=7),
          legend.position="top",
          legend.background = element_blank(),
          legend.title = element_blank(),
          legend.text = element_text(size=8),
          legend.key.size = unit(0.3, "cm"),
          legend.key = element_blank(),
          panel.background = element_rect(fill = "transparent"),
          plot.background = element_rect(fill = "transparent", color = NA)) +
    scale_x_continuous(breaks = 1:6, labels = c("MII", "2C", "4C", "8C", "16C/MO", "BL"))
  
  ggsave(FIG,
         file=path,
         bg = "transparent",
         width = 3.23,
         height=3,
         units = "in",
         dpi = 600)
}
```

---

#### Session Info

Detail of the `R` session info for reproducibility.

```
sessionInfo()
```

```
## R version 4.0.2 (2020-06-22)
## Platform: x86_64-apple-darwin17.0 (64-bit)
## Running under: macOS  10.16
## 
## Matrix products: default
## BLAS:   /Library/Frameworks/R.framework/Versions/4.0/Resources/lib/libRblas.dylib
## LAPACK: /Library/Frameworks/R.framework/Versions/4.0/Resources/lib/libRlapack.dylib
## 
## locale:
## [1] en_US.UTF-8/en_US.UTF-8/en_US.UTF-8/C/en_US.UTF-8/en_US.UTF-8
## 
## attached base packages:
## [1] stats     graphics  grDevices utils     datasets  methods   base     
## 
## other attached packages:
##  [1] limma_3.46.0        gridExtra_2.3       viridis_0.5.1      
##  [4] viridisLite_0.4.0   ggridges_0.5.3      caret_6.0-86       
##  [7] lattice_0.20-41     randomForest_4.6-14 ellipse_0.4.2      
## [10] ggplot2_3.3.3       FactoMineR_2.4     
## 
## loaded via a namespace (and not attached):
##  [1] sass_0.3.1           jsonlite_1.7.2       splines_4.0.2       
##  [4] foreach_1.5.1        prodlim_2019.11.13   bslib_0.2.4         
##  [7] assertthat_0.2.1     highr_0.8            stats4_4.0.2        
## [10] yaml_2.2.1           ggrepel_0.9.1        ipred_0.9-11        
## [13] pillar_1.6.1         glue_1.4.2           pROC_1.17.0.1       
## [16] digest_0.6.27        colorspace_2.0-1     recipes_0.1.15      
## [19] htmltools_0.5.1.1    Matrix_1.3-2         plyr_1.8.6          
## [22] timeDate_3043.102    pkgconfig_2.0.3      purrr_0.3.4         
## [25] scales_1.1.1         gower_0.2.2          lava_1.6.9          
## [28] tibble_3.1.2         farver_2.1.0         generics_0.1.0      
## [31] ellipsis_0.3.2       DT_0.17              withr_2.4.2         
## [34] nnet_7.3-15          survival_3.2-7       magrittr_2.0.1      
## [37] crayon_1.4.1         evaluate_0.14        fansi_0.4.2         
## [40] nlme_3.1-152         MASS_7.3-53.1        class_7.3-18        
## [43] tools_4.0.2          data.table_1.14.0    lifecycle_1.0.0     
## [46] stringr_1.4.0        munsell_0.5.0        cluster_2.1.2       
## [49] flashClust_1.01-2    compiler_4.0.2       jquerylib_0.1.3     
## [52] rlang_0.4.11         grid_4.0.2           iterators_1.0.13    
## [55] htmlwidgets_1.5.3    leaps_3.1            labeling_0.4.2      
## [58] rmarkdown_2.7        gtable_0.3.0         ModelMetrics_1.2.2.2
## [61] codetools_0.2-18     DBI_1.1.1            reshape2_1.4.4      
## [64] R6_2.5.0             lubridate_1.7.10     knitr_1.31          
## [67] dplyr_1.0.5          utf8_1.2.1           stringi_1.5.3       
## [70] Rcpp_1.0.6           vctrs_0.3.8          rpart_4.1-15        
## [73] scatterplot3d_0.3-41 tidyselect_1.1.0     xfun_0.22
```
